# Spatiotemporal coordination at the maternal-fetal interface promotes trophoblast invasion and vascular remodeling in the first half of human pregnancy

**DOI:** 10.1101/2021.09.08.459490

**Authors:** Shirley Greenbaum, Inna Averbukh, Erin Soon, Gabrielle Rizzuto, Alex Baranski, Noah F. Greenwald, Adam Kagel, Marc Bosse, Eleni G. Jaswa, Zumana Khair, Shirley Kwok, Shiri Warshawsky, Hadeesha Piyadasa, Geneva Miller, Morgan Schwartz, Will Graf, David Van Valen, Virginia D. Winn, Travis Hollmann, Leeat Keren, Matt van de Rijn, Michael Angelo

## Abstract

Beginning in the first trimester, fetally derived extravillous trophoblasts (EVTs) invade the uterus and remodel its spiral arteries, transforming them into large, dilated blood vessels left with a thin, discontinuous smooth muscle layer and partially lined with EVTs. Several mechanisms have been proposed to explain how EVTs coordinate with the maternal decidua to promote a tissue microenvironment conducive to spiral artery remodeling (SAR). However, it remains a matter of debate which immune and stromal cell types participate in these interactions and how this process evolves with respect to gestational age. Here, we used a multiomic approach that combined the strengths of spatial proteomics and transcriptomics to construct the first spatiotemporal atlas of the human maternal-fetal interface in the first half of pregnancy. We used multiplexed ion beam imaging by time of flight (MIBI-TOF) and a 37-plex antibody panel to analyze ∼500,000 cells and 588 spiral arteries within intact decidua from 66 patients between 6-20 weeks of gestation, integrating this with coregistered transcriptomic profiles. Gestational age substantially influenced the frequency of many maternal immune and stromal cells, with tolerogenic subsets expressing CD206, CD163, TIM-3, Galectin-9, and IDO-1 increasingly enriched and colocalized at later time points. In contrast, SAR progression preferentially correlated with EVT invasion and was transcriptionally defined by 78 gene ontology pathways exhibiting unique monotonic and biphasic trends. Lastly, we developed an integrated model of SAR supporting an intravasation mechanism where invasion is accompanied by upregulation of pro-angiogenic, immunoregulatory EVT programs that promote interactions with vascular endothelium while avoiding activation of immune cells in circulating maternal blood. Taken together, these results support a coordinated model of decidualization in which increasing gestational age drives a transition in maternal decidua towards a tolerogenic niche conducive to locally regulated, EVT-dependent SAR.

## Introduction

Normal development during healthy pregnancy depends on a complex interplay between maternal cells and placental trophoblasts that ultimately transforms the womb into a specialized niche capable of meeting the metabolic demands of a growing semi-allogeneic fetus while maintaining maternal tolerance^1–5^. Rather than being a single monotonic trend, this process is multifaceted and dynamic with respect to both tissue structure and gestational age (GA). After implantation, decidual cellular composition shifts to one that is enriched for invasive extravillous trophoblasts (EVTs)^6^. During this transition, maternal and fetal cells remodel uterine spiral arteries into highly dilated vessels with minimal smooth muscle where EVTs have partially replaced the maternal endothelium within the arterial lumen^7–9^. Spiral artery remodeling (SAR) in healthy pregnancies results in low-resistance vessels that can deliver blood to the intervillous space at low flow velocities that prevent damage to the placental architecture^10, 11^. Conversely, impaired SAR, fewer tolerogenic maternal cells, and abnormal decidual invasion of EVTs have each been implicated in placenta-related obstetric complications, including preeclampsia, intrauterine growth restriction, and preterm birth^12, 13^. Therefore, a detailed investigation of the cell population dynamics at the maternal-fetal interface is key to understanding the biology of normal pregnancy and the pathophysiology of placenta-related obstetric complications.

Due to the poor feasibility of controlled studies in pregnant humans, much of what is known about maternal-fetal tolerance and SAR is based on pregnancy in small mammals^14^. Although some similarities exist, key facets of hemochorial placentation in humans are primate-specific, and in some cases are restricted even further to great apes^15–17^. For example, giant cells in mice only invade the superficial decidua, do not replace the vascular endothelium, and are thought to play a minor role in SAR compared to maternal uterine natural killer (NK) cells^18^. In contrast, EVTs in humans invade completely through the decidua into the inner third of the myometrium and are considered to be vital for adequate SAR^3, 19, 20^. Since the most extensive EVT invasion has been observed in humans, it may be a key adaptation that permitted upright, bipedal locomotion while maintaining adequate blood flow in the third trimester when development of the large fetal brain accounts for 60% of metabolic needs ^21, 22^.

The study of human decidual remodeling is further complicated by additional inherent challenges. First, cell composition and structure are temporally dynamic; aggregating data across different GAs or observing a single time point may be misleading. As the decidua develops^23, 24^, the functions of maternal NK cells, T cells, and macrophages change dynamically in the first and second trimesters to promote a permissive niche conducive to villus attachment and invasion. This process necessarily establishes a gradient of EVT invasion that advances inward from the superficial decidua. Consequently, decidual structure and composition in focal regions can differ significantly from its bulk attributes.

A second major challenge arises in understanding how these global dynamics are coupled to processes requiring spatial coordination, such as those between maternal and placental cells in the local tissue microenvironment. For example, periarterial decidual NK cells are thought to contribute to SAR both by initiating smooth muscle breakdown and by secreting chemokines that attract invading EVTs, while phagocytic macrophages are thought to facilitate clearance of the resultant apoptotic debris^25–27^. Overall, formation of the human maternal-fetal interface involves sophisticated spatiotemporal coordination such that tissue composition, structure, and function are inextricably coupled. Unraveling this interdependence requires an approach that can ascertain how these facets change over time in intact human tissue.

With this in mind, we constructed the first high dimensional spatio-temporal atlas of the human maternal-fetal interface. We leveraged archival tissue banks to assemble a cohort of maternal decidua from 66 women, who underwent elective terminations of otherwise healthy pregnancies at 6-20 weeks gestation, the largest single-cell study of the maternal-fetal interface to date^4, 28^. We performed high dimensional, subcellular imaging with multiplexed ion beam imaging by time of flight (MIBI-TOF) ^29^ using a 37-plex antibody panel designed to comprehensively identify the location, lineage, and function of all major maternal and placental cells. Building upon previous studies of aggregate tissue across the entire first or second trimester^4, 28^, we were able to examine dynamic changes with respect to GA, allowing us to elucidate important temporal patterns not previously observed.

We also profiled the transcriptome of arteries, adjacent decidua and EVT on serial sections with Nanostring DSP, generating a multi-modal atlas of decidual remodeling. To understand how SAR relates to local decidual composition, we developed new algorithms for quantifying vascular morphology that enabled us to assign a remodeling score to each individual artery. We discerned which changes in decidual composition, transcriptome and structure were preferentially driven by GA, SAR, or both. Overall, the frequencies, relative proportions and spatial arrangement of maternal immune cells exhibited a robust temporal dependence that permitted us to predict GA based on immune cellular composition alone.

In contrast, we found that EVT invasion and perivascular localization were the dominant drivers of SAR in the tissue microenvironment and that these processes correlated with extensive shifts in arterial transcription. Given these findings, we then used our atlas to compare two models for the path of EVT migration from the cytotrophoblast anchoring cell columns to maternal spiral arteries that have been proposed previously: (1) intravasation, where EVTs first invade the decidua and then enter arteries by traversing the arterial wall, and (2) extravasation, where EVTs enter arteries directly at the basal plate^9^. Using statistical analyses correlating EVT phenotype and location with the extent of arterial smooth muscle and endothelial loss, we found that our spatiotemporal atlas was most consistent with an intravasation model. Leveraging the spatial nature of our data, we were able to characterize the differential gene expression between interstitial and intravascular EVT populations and predict how these drive SAR via cell-cell interactions between intravascular EVTs and arteries. Taken together, these investigations support a cooperative interplay in the first half of pregnancy in which temporally dependent changes in decidual function permit placental EVTs to extensively alter the maternal uterine vasculature.

## Results

### Multiplexed imaging of human decidua reveals the tolerogenic composition of the maternal-fetal interface

As part of the Human BioMolecular Atlas Program (HuBMAP) initiative, we created the first spatio-temporal tissue atlas of the human maternal-fetal interface in the first 20 weeks of pregnancy (Fig. 1a). The goal of this study was to comprehensively define the structure and composition of decidua and to understand how it changes during the first two trimesters with respect to two axes: GA and maternal SAR. To examine these questions, we first assembled a large retrospective cohort of archival formalin-fixed, paraffin-embedded placenta and decidua tissue from 66 patients who underwent elective termination of pregnancies with no known fetal abnormalities. Archival tissue blocks were manually screened by a perinatal pathologist in hematoxylin and eosin (H&E) stained tissue sections to determine which samples contained decidua. Then, regions of decidua that contained spiral arteries were demarcated, cored, and assembled into two tissue microarrays (TMAs) of 1mm and 1.5 mm cores. The final dataset included samples for 6-20 weeks of gestation (13.72±4.8 weeks) from 66 patients of varying parity (1.45±1.72), age (28.17±5.9 years), body mass index (28.19±7.3 kg/m^2^), and ethnicity (Fig. 1b-f, Supplementary Table 1). Due to inherent limitations in how the tissue was procured, precise anatomic locations could not be determined. However, 61 out 66 tissue blocks contained placental villi, suggesting that the vast majority of this cohort was sampled from decidua basalis (Supplementary Table 2, see Methods).

**Figure 1.**
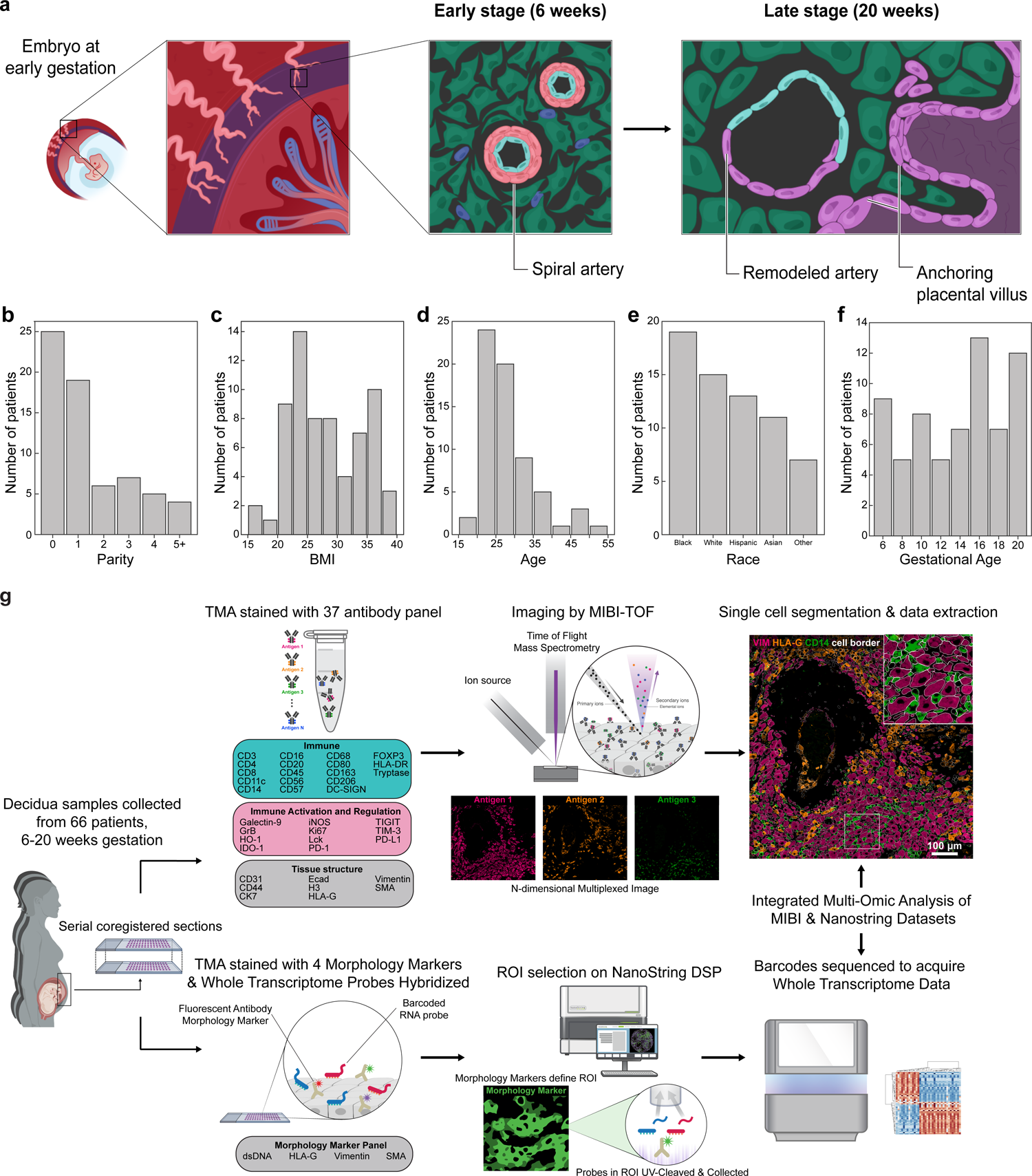
I Study design and workflow. **a.** Diagram of a human embryo in utero at 6 weeks gestation. First inset: the maternal-fetal interface consisting of decidua basalis (purple) with maternal spiral arteries (light pink) and fetal chorionic villi in the intervillous space (bottom right corner). Second inset: early stage (6 weeks) unremodeled spiral artery and progression to late stage (20 weeks) remodeled artery and anchoring fetal villi. **b.** Cohort parity distribution. **c.** Cohort distribution of body mass index (BMI). **d.** Cohort age distribution. **e.** Cohort ethnicity distribution. **f.** Cohort distribution of gestational age (GA). GA in days is binned to weeks for visualization. **g.** TMA construction and serial sections for multi-omic workflow. Upper: antibody panel, MIBI workflow, and single-cell segmentation. Bottom: morphology marker panel and probe diagram, Nanostring DSP workflow, and barcode sequencing for acquisition of Whole Transcriptome data.

Previous studies of intact tissue examining only one or a few cell populations at a time reported shifts in maternal immune cells towards tolerogenic states that are permissive to invasion by fetally derived EVTs^19^. To gain a more complete picture of the complex cell-cell interactions that establish maternal tolerance in the first half of pregnancy, we combined the strengths of targeted subcellular imaging with antibodies and spatial transcriptomics on serial co-registered sections to construct a comprehensive composite model of SAR and decidual remodeling (Fig. 1g).

For MIBI-TOF, we designed and validated a 37-plex antibody panel for simultaneously mapping the functional state and location of all major maternal and fetal cell populations simultaneously (Fig. 1g, see Methods, Extended Data Fig. S1). In addition to canonical lineage defining markers for fetal cells, maternal immune cells, fibroblasts, smooth muscle, endothelium, and epithelium, we also quantified 10 functional markers previously implicated in maternal immune tolerance, including TIM-3, Galectin-9, PD-1, PDL-1, and IDO-1 (Fig. 1g, Extended Data Fig. S1)^30–35^. All TMA sections were stained simultaneously with this antibody panel and subsequently imaged at 500 nm resolution (Fig. 1g).

For spatial transcriptomics, we used the NanoString GeoMx™ Digital Spatial Profiler (DSP) for whole transcriptome analysis of decidua, arteries, and EVTs (Fig. 1g). Immunofluorescence imaging of TMAs stained with antibodies for HLA-G, VIM, SMA, and a nucleic acid stain were used to precisely define regions of interest (ROIs) specific for each of these histologic features (see Methods). Barcodes bound to hybridized RNA probes in each ROI were then selectively photocleaved, collected, and sequenced, generating a spatially resolved map of transcription within the tissue (Fig. 1g). In total, we collected whole transcriptome data from 13 individual arteries, their adjacent decidua, 5 samples of interstitial and 3 samples of intravascular EVT (19 cores from 17 patients, see Methods).

MIBI images were denoised with a low-level image analysis pipeline as described previously (Fig. 1g)^36^. To accurately capture the unique diversity of morphologically distinct maternal and fetal cells, we used our previously validated custom whole cell convolutional neural network, Mesmer^37^ (see Methods, Extended Data Fig. S2a). We optimized this neural network for decidua-specific segmentation by training with 93,000 manually annotated single cell events from 25 decidual images. Applying this segmentation algorithm to our cohort images yielded 495,349 segmented cells in total, identified across 211 images (800μmx800μm, 2347±783 cells per image). FlowSOM clustering^38^ was used to assign 92% of whole cell segmented events to 25 cell populations (Fig. 2a, b, see Methods, Extended Data Fig. S2b-f). These data (Fig. 2c-g) were then combined with whole-cell segmentation masks to generate cell phenotype maps (CPM) in which each cell is colored categorically by its respective population (Fig. 2h, Extended Data Fig. S2g). We then determined whether cells expressed the functional markers by applying manually derived per-marker binary expression thresholds (see Methods). Noteworthy histological features—such as arteries, vessels, glands, the cell columns, and decidual tissue boundaries—were manually annotated in collaboration with a perinatal pathologist.

**Figure 2.**
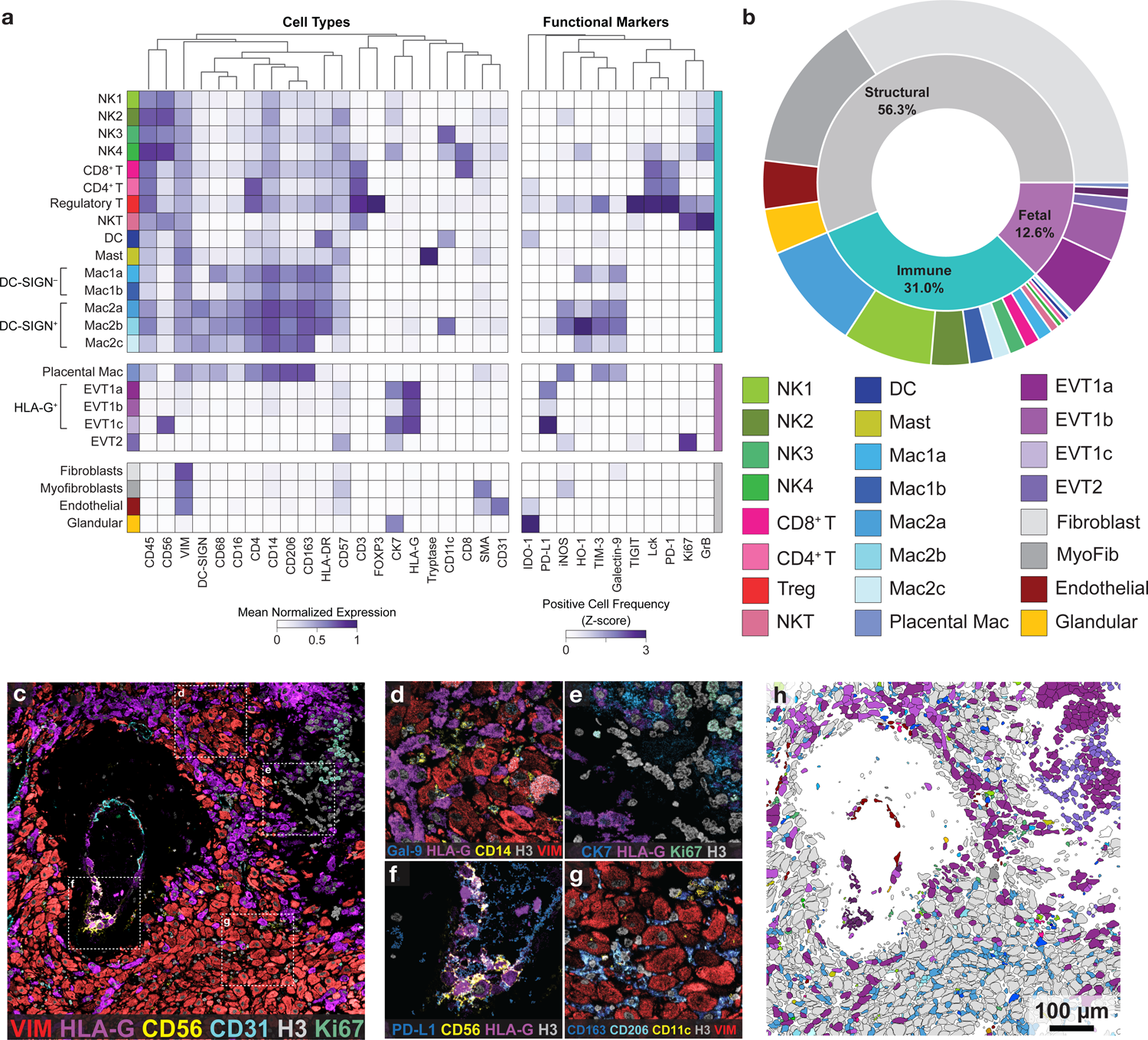
I Multiplexed imaging of human decidua reveals the immune tolerance-conducive composition of the maternal fetal interface. **a.** Cell lineage assignments showing mean normalized expression of lineage markers (left) and functional marker positive cell frequency (right, Z-score). Columns (markers) are hierarchically clustered. **b.** Cell lineage abundances across our cohort. Placental mac: Placental macrophage; MyoFib: Myofibroblast. **c.** Representative MIBI field of view color overlay of a 20 week sample. Red = VIM, vimentin, purple = HLA-G, yellow = CD56, cyan = CD31, grey = H3, green = Ki67. **d.** Inset of **c**, interstitial fetal EVTs. Blue = Galectin-9, purple = HLA-G, yellow = CD14, grey = H3, red = VIM **e.** Inset of **c**, showing anchoring villous cell column to decidua interface. Blue = CK7, cytokeratin7, purple = HLA-G, green = Ki67, grey = H3. **f.** Inset of **c**, showing intravascular EVTs. Blue = PD-L1, purple = HLA-G, yellow = CD56, grey = H3. **g.** Inset of **c**, showing decidual stromal cells (fibroblasts) and macrophages. Blue = CD163, cyan = CD206, yellow = CD11c, grey = H3, red = vimentin. **h.** Cell lineage assignments overlaid onto the cell-segmentation output to produce a cell phenotype map.

Non-immune maternal (structural) cells accounted for the majority (56.3%) of all segmented events in the decidua and were predominantly composed of decidual fibroblasts (60.5%) and myofibroblasts (24.8%) with smaller contributions from vascular endothelium (7.6%) and glandular epithelial cells (7.1%, Fig. 2b). Interestingly, we observed a novel, rare subset of TIGIT+ glandular cells (0.34% of glandular cells, Extended Data Fig. S2h). Consistent with work^39^ quantifying maternal populations *in situ*, maternal immune cells (31% of all cells) were dominated by macrophages (47.6% of immune) and NK cells (42.6% of immune) with minor contributions from T (8% of immune), dendritic (1.3% of immune), and mast cells (0.5% of immune).

Decidual macrophages (CD14^+^) ubiquitously co-expressed CD163 and CD206, consistent with an M2-polarized, tolerogenic phenotype^40^ (Fig. 2g). In line with previous work showing pregnancy-specific recruitment, 77% of macrophages expressed DC-SIGN^41^ (Fig. 2a). We further classified DC-SIGN^+^ macrophages into three subsets (Mac2a, 2b, 2c) based on expression of CD11c (Mac2b, 2.7% of macrophages) or absence of HLA-DR (Mac2c, 10.3% of macrophages). The majority (64%) of macrophages were CD11c^-^HLA-DR^+^ (Mac2a). DC-SIGN-macrophages were subsetted based on CD68 expression (CD68^-^ Mac1b and CD68^+^ Mac1a) (Fig. 2a, b). Notably, Mac2 subsets were also found to express multiple immune regulatory proteins, including TIM-3, Galectin-9, iNOS, and HO-1

Four subsets of NK cells (CD3^-^CD56^+^) were identified based on combinatorial co-expression of CD57, CD11c, and CD8. NK1 (CD57^-^CD16^low^) was the largest NK cell population present, making up 59.7% of NK cells (Fig. 2a, b). We found a novel CD57+ population of decidual NK cells (NK2, 25.8% of NK cells), which had only been previously identified in peripheral blood during pregnancy^42^. As described below, its frequency and spatial distribution suggest NK2s play a distinct role in spiral artery remodeling (Extended Data Fig. S3j). The remaining two subsets could be distinguished based on expression of CD11c (NK3, 11.3% of NK cells), or CD8 (NK4, 3.2% of NK cells) (Fig. 2a, b).

T cells consisted of CD8^+^ (53.2% of T cells), CD56^+^ NKTs (28.8% of T cells), CD4^+^ (17.1% of T cells), and sparse numbers of regulatory T (Treg) cells (CD4^+^FOXP3^+^, 0.7% of T cells), while no B cells were observed (Fig. 2a, b). We identified several populations of decidual Tregs, including a PD-1^+^ activated population (31% PD-1^+^ Tregs), and an intriguing TIM-3^+^Lck^+^ subset that accounted for >50% of this population (Fig. 2a, b). Interestingly, both Tregs and NKT cells were highly proliferative (13.7% Ki67^+^ Tregs, 17% Ki67^+^ NKT cells), and with the notable exception of CD8^+^ NK cells (22.9% GrB^+^), had higher frequencies of GrB (Granzyme B) expressing cells than any NK cell subset (33.7% GrB^+^ NKT, 19.5% GrB^+^ Treg). TIGIT was most frequently expressed by Tregs—a rare subset that has been suggested to bind PVR (CD155) on EVTs^4^; this interaction has been observed in the tumor microenvironment^43^ and may contribute to driving Treg mediated immunosuppression at the maternal-fetal interface.

Fetal cells (12.6% of all cells) were primarily comprised of four subsets of EVTs that were delineated based on combinatorial expression of HLA-G, CK7, CD57, and CD56 (Fig. 2a). HLA-G^+^ interstitial EVT populations were CK7^+^ (EVT1a, 44.6% of fetal cells), CK7^-^ (EVT1b, 35.3% of fetal cells), or CD56^+^ (EVT1c, 6.9% of fetal cells) (Fig. 2c-f). EVT2 lacked HLA-G and were CD57^-^CK7^low^ and were located predominantly at the base of attaching cell columns (EVT2, 9.4% of fetal cells). Notably, placental macrophages (Hoffbauer cells) located in chorionic villi constituted the remainder (4.1%) of fetal cells and exhibited a cellular phenotype similar to that of Mac2c (DC-SIGN^+^HLA-DR^-^) decidual macrophages (Fig. 2a).Taken together, these data provide spatial context to prior work using dissociated samples ^28, 33, 41, 44^ where an ensemble of functional states in fetal and maternal cells are collectively aligned in maintaining a tolerogenic niche.

### SAR progression is tightly correlated with the local tissue microenvironment

Perfusion of the intervillous space by uterine spiral arteries is the sole source of oxygen and nutrients to the growing fetus after the establishment of arterial flow. During the first half of pregnancy, these vessels undergo an extensive remodeling process that culminates in dilated, non-contractile vessels depleted of smooth muscle where the maternal endothelium has been partially replaced by EVTs. While abnormal SAR is associated with obstetric complications, such as intrauterine growth restriction and preeclampsia^12, 13^, it is still not fully understood which cell populations directly participate in SAR, how this process is locally regulated, and to what extent these changes are synchronized with GA.

We therefore used our spatiotemporal atlas of decidua to construct a SAR trajectory and understand how this process relates to temporal changes in decidua cell composition and structure. Using artery size, smooth muscle disruption, endothelial continuity, and EVT infiltration, we manually assigned each artery to one of five sequential remodeling stages based on previously published criteria^45^ (Fig. 3a). To ensure scoring was not biased by patient demographics or the composition of neighboring arteries and stroma, scoring was performed on cropped images by blinded experts where only the artery of interest was visible. Out of 588 arteries, 186 were unremodeled and assigned to Stage 1 (Fig. 3b, c). Stage 2 arteries (300 arteries) were characterized by moderate smooth muscle disruption and endothelial swelling (Fig. 3d, e). Stage 3 arteries (43 arteries) exhibited more dilation, smooth muscle loss, and early endothelial disruption (Fig. 3f, g). Progression to Stage 4 (34 arteries) was marked by the presence of EVTs within the arterial lumen (Fig. 3h,i), while fully remodeled Stage 5 arteries (25 arteries) were identified based on their very large size, near-complete smooth muscle loss, and EVT endothelization (Fig. 3j, k, see Methods, Extended Data Fig. S3a, Supplementary Table 3).

**Figure 3.**
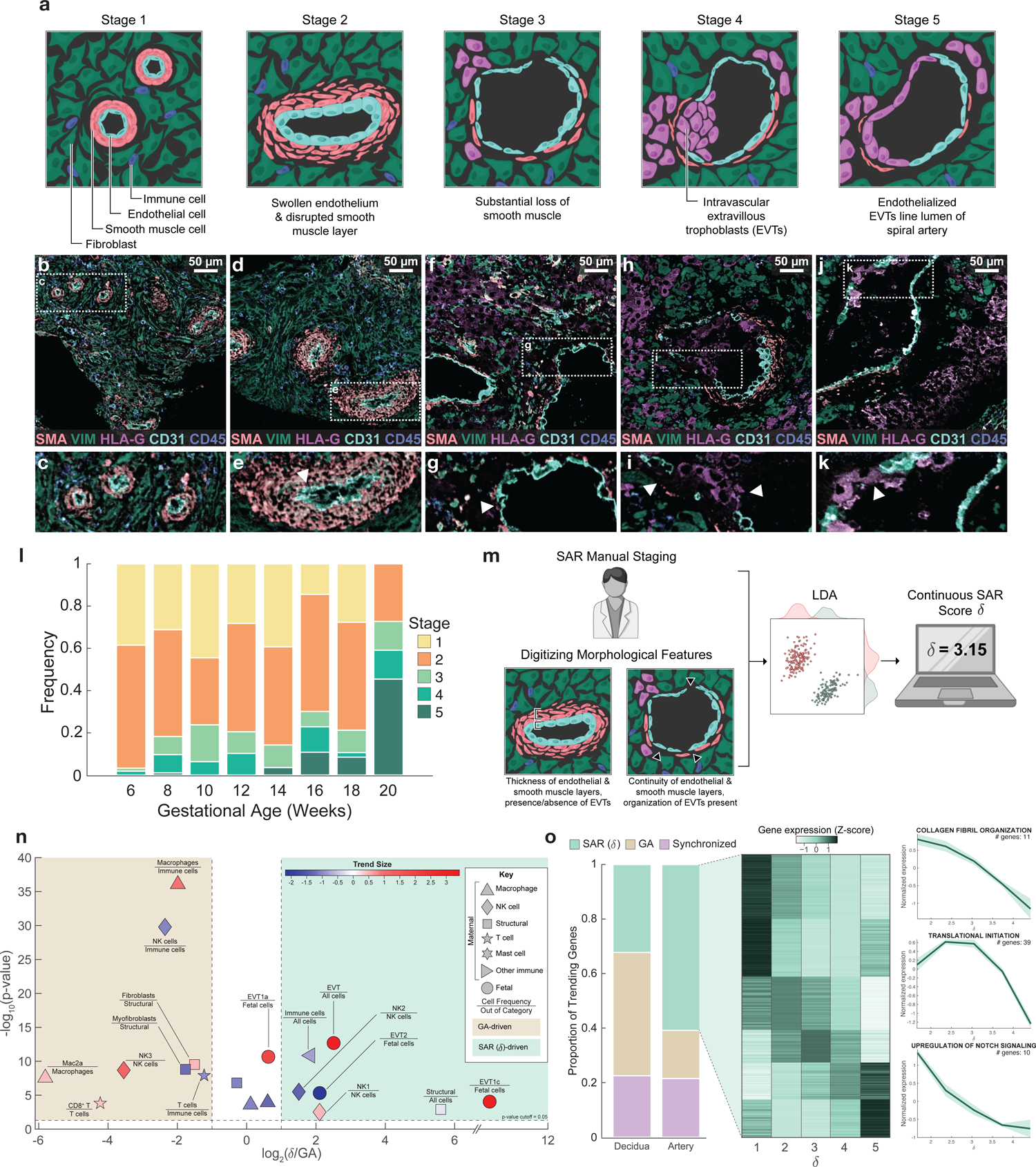
I Spiral Artery Remodeling (SAR) progression significantly influences maternal-fetal interface composition. **a.** Diagram showing key characteristics of SAR Stages 1-5, assessed manually. **b.** Representative MIBI color overlay of SAR manual Stage 1 arteries. VIM, vimentin; SMA, smooth muscle actin. **c.** Inset of **b**, showing SAR manual Stage 1 arteries. **d.** Representative MIBI color overlay of SAR manual Stage 2 arteries. **e.** Inset of **d**, showing one SAR manual Stage 2 artery. Arrowhead; swollen endothelial cells. **f.** Representative MIBI color overlay of SAR manual Stage 3 arteries. **g.** Inset of **f**, showing one SAR manual Stage 3 artery. Arrowhead; substantial loss of smooth muscle **h.** Representative MIBI color overlay of one SAR manual Stage 4 artery. **i.** Inset of **h**, showing one SAR manual Stage 4 artery. Arrowheads; intravascular EVTs. **j.** Representative MIBI color overlay of one SAR manual Stage 5 artery. **k.** Inset of **j**, showing one SAR manual Stage 5 artery. Arrowhead; endothelialized intravascular EVTs lining the spiral artery lumen. **l.** Distribution of SAR manual stages by gestational age (GA). GA in days is binned to weeks for visualization. **m.** Schematic of calculating the continuous SAR remodeling score (δ). Manual stages along with quantified digitized morphological features were used to construct a trajectory of SAR using LDA from which the continuous SAR score δ was calculated. **n.** Volcano plot distinguishing GA-driven from SAR (δ)-driven cell-type frequencies. X axis: log_2_ ratio of R^2^ derived from linear regression against SAR (δ) and GA. Y axis: -log_10_ of the p-value for the better-fitting regression model. Points are color coded by the trend size observed in the better-fitting regression model. **o.** Proportion of genes in artery (2932 total) and decidua (1722 total) tissue where expression changes significantly correlate with GA (517 artery, 778 decidua), SAR (δ) (1785 artery, 555 decidua), or both (633 artery, 389 decidua). Center inset: 778 SAR (δ)-correlated genes in artery tissue showing mean normalized expression (Z-score) by SAR (δ) stage. Right inset: 3 SAR (δ)-trending gene ontology pathways, showing normalized expression of genes in the GO pathway by SAR (δ)

Although SAR correlated with GA to some extent (Spearman’s ρ=0.28, p-value = 1.5*10^-12^), artery staging and GA in many cases were discordant. For example, at least one late-stage artery (Stage 4-5) was present in 40% of week 8 samples, while minimally remodeled arteries were present throughout (Fig. 3l). Moreover, SAR staging of arteries from the same patient often varied significantly between tissue cores (32% of patients had arteries that differed by at least two stages), suggesting that this discordance could be highlighting aspects of SAR that are locally regulated by the tissue microenvironment (Fig. 3l, Extended Data Fig. S3b).

This decoupling of SAR and GA permitted us to identify changes in decidual composition that were predominantly driven by one or the other. We first developed a quantitative staging scheme for automatically assigning a continuous remodeling score. For each artery, we extracted 35 parameters describing the same aspects of arterial morphology that were used for manual scoring (Fig. 3m, see Methods, Extended Data Fig. S3c,d). Together with manual staging, we used this quantitative morphologic profile to construct a highly resolved pseudotime trajectory of SAR using linear discriminant analysis (LDA)^46^ (Fig. 3m, see Methods). We generated this trajectory by combining the 35 morphological features with our manually defined stage labels and applying LDA to project each artery with respect to a two-dimensional LDA space in which separation of arteries by their manually assigned stages is optimal (see Methods). We then defined a remodeling trajectory as the polynomial fit to artery points in this space and subsequently mapped each artery (a_i_) to the nearest point along this curve (b_i_, Extended Data Fig. S3e, see Methods). Finally, a remodeling score (δ) was determined by calculating the distance along this curve from the point of origin (x_o_) to b_i_ for each artery (See integral in Extended Data Fig. S3e, Extended Data Fig. S3f-g, Supplementary Table 3).

With our continuous remodeling score δ, we next defined a simple scheme to differentiate GA- and SAR-driven trends by performing linear regressions of cell frequency per image both as a function of GA and as a function of δ. Regression R^2^ and p-values were used as proxies for trend quality and to assess significance, respectively. Trends where R^2^ for GA and SAR differed by at least two-fold were classified as being driven predominately by a single process, while ones falling below this cutoff were classified as synchronized (Fig. 3n, see Methods, Extended Data Fig. S3h, i).

The frequency of decidual EVTs was better correlated with SAR (Log_2_ R^2^ ratio(δ:GA) = 2.5, p-value for δ =1e^-13^) while changes in the proportion of maternal immune cells were mostly driven by GA (Fig. 3n). One notable exception to the latter was observed within the NK cell compartment, where the ratio of NK2 (CD57^+^) to NK1 (CD57^-^) was found to drop in an SAR-dependent manner (Log_2_ R^2^ ratio(δ:GA) ≥ 1.5, p-value for δ ≤ 0.003, Fig. 3n).

To further investigate this finding, we examined how the spatial distribution of NK cells near arteries changed as SAR progressed (see Methods). Remarkably, we found that NK2s were the only subset of maternal immune cells to preferentially localize around arteries (Supplementary Table 4). NK2 accumulation spiked specifically at stage 2 of SAR when smooth muscle swelling and disruption are maximal (p=2e^-3, Extended Data Fig. S3j). Interestingly, CD57 expression in human NKs is associated with a cytotoxic phenotype in tumors ^42^, suggesting that this subset could be serving a similar role in mediating early smooth muscle disruption during SAR.

To create a transcriptional trajectory that is integrated with our proteomics atlas, we used Nanostring DSP on serial sections of the TMA imaged by MIBI-TOF. We collected paired, whole transcriptome tissue profiles of 13 individual arteries and adjacent decidua of varying stages of remodeling (26 ROIs total) using patient-matched tissue cores (see Methods). Paring these samples with their respective MIBI-TOF images allowed us to assign a remodeling score and gestational age to each of these transcriptome profiles (see Methods). In a similar fashion to the approach used in Fig. 3n, we set out to divide genes based on their association with SAR or GA (see Methods). Changes in expression of 2935 out of 18695 genes were found to significantly correlate with SAR, GA, or both (i.e. synchronized).

Despite their close spatial proximity, the proportion of GA or SAR associated transcriptional changes in arteries and decidua markedly differed. A plurality of decidual genes and majority of arterial genes were preferentially correlated with GA or SAR, respectively (Fig. 3o, Extended Data Fig. S3k). With respect to the latter, we identified 78 functional, temporally synchronized gene ontology pathways, including modules related to vessel remodeling processes and translation (see Methods, Supplementary Table 5, Fig. 3o). These pathways were found to exhibit both monotonic and biphasic trends (Fig. 3o) showing that SAR is a composite of interrelated processes occurring continuously and episodically. For instance, we identified 185 genes that were maximally expressed in stage 2 before subsequently declining. These genes correlated with perivascular enrichment of NK2s found in our MIBI data (Extended Data Fig. S3j) and are enriched for pathways related to collagen fibril organization (Fig. 3o) and responses to Bone Morphogenic Protein (Extended Data Fig. S3i). Consistent with cell growth and subsequent apoptosis of arterial smooth muscle, translation related pathways followed a biphasic trend, peaking around stage 2-3 of remodeling (Fig. 3o). We also observed continual downregulation of genes involved in Notch signaling as SAR progressed (Fig. 3o). Taken together, these multi-modal data provide a fully integrated atlas of decidual remodeling describing tissue structure, single cell function, and changes in transcriptional programs.

### Changes in immune cell composition and spatial enrichment are tightly correlated with Gestational Age

We next interrogated these data to identify temporal changes in decidual composition, which revealed a robust, GA-dependent shift from a lymphoid to myeloid dominated immune landscape, characterized by fewer NK and T cells and a concomitant increase in macrophage frequency (Log_2_ R^2^ ratio(δ:GA) ≤ −1.2, p-value for δ ≤ 1.2e^-8^, Figs. 3n, 4a, b). Images at weeks 6-8 (Fig. 4c, e), show NK cells and T cells, including those exhibiting cytotoxic (Fig. 4d) and immunosuppressive (Fig. 4e) phenotypes, greatly outnumbered macrophages (Fig. 4b, c). Contrastingly, images from weeks 16-20 were dominated by interstitial EVTs (Figs. 4a, 4f-g) and an accompanying increase in tolerogenic macrophage populations (Fig. 4h) compared to NK and T cells. To further evaluate this relationship, we asked whether immune cell composition in the decidua alone could be used to predict GA. Using immune features that were preferentially associated with GA (Fig. 3n), we trained and validated a ridge regression model on a per-image basis using a random 70/30 test-train split (Extended Data Fig. S4a). Remarkably, the trained model predicted GA in the withheld test set within 19 days of the ground-truth value (R^2^=0.7, Fig. 4i). On inspecting the model weights, we found that the relative contribution of decidual immune cells was consistent with the observed shift in the proportion of myeloid and lymphoid cells. In particular, the relative frequencies of T and NK cells were negatively correlated with GA, while total macrophage frequency was positively correlated with GA (Fig. 4j). Notably, a modified regression model for predicting SAR (δ) based on the same immune cell population parameters performed poorly (R^2^=0.05, RMSE=0.85, Extended Data Fig. S4b), reinforcing our hypothesis that these immune correlates are driven by GA and not SAR.

**Figure 4.**
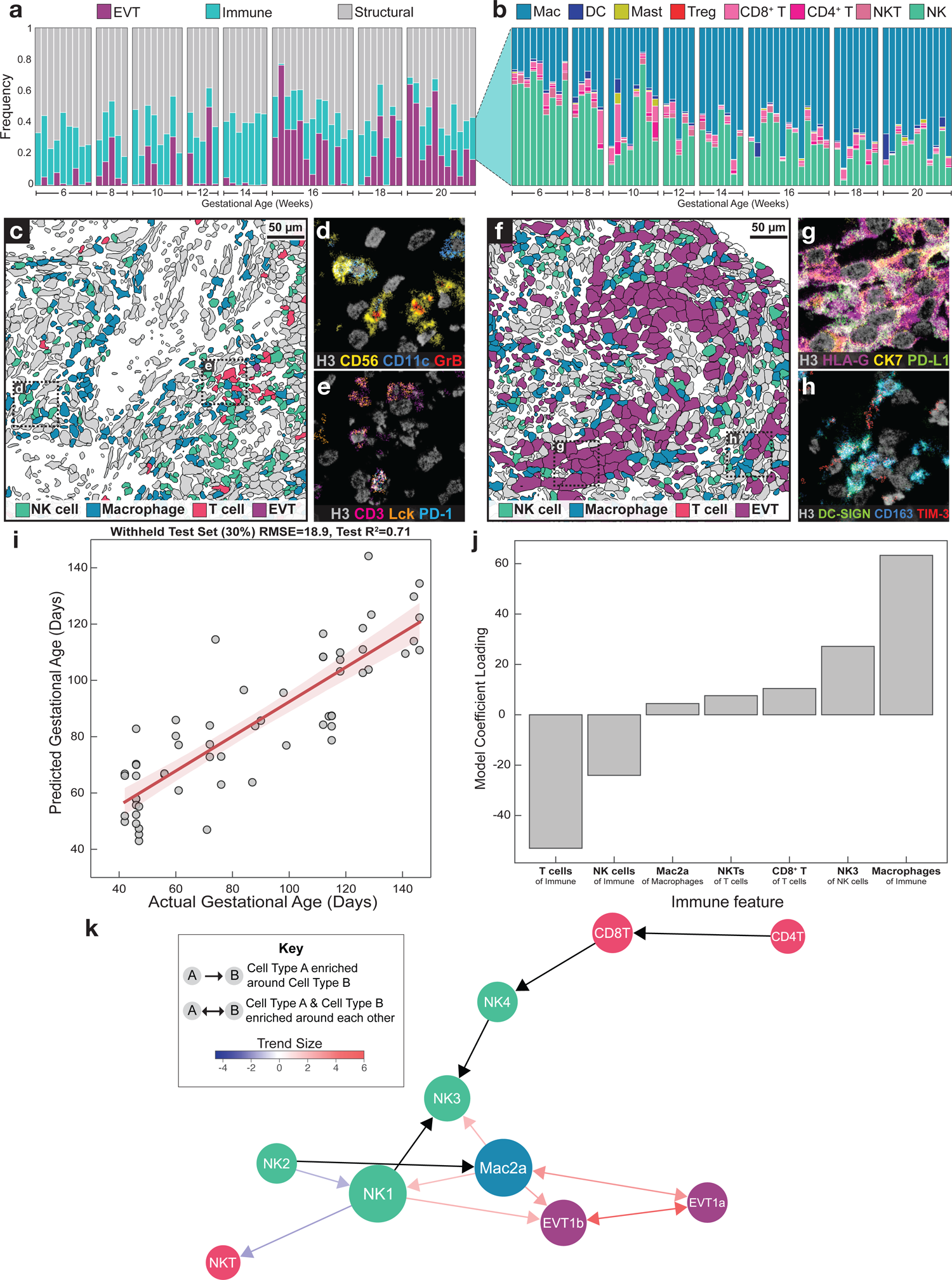
I A lymphoid-to-myeloid shift in immune-compartment composition is tightly correlated with gestational age (GA). **a.** Frequency of EVT, immune, and structural cell populations per patient, with patients ordered by GA. GA in days is binned to weeks for visualization. **b.** Frequency of immune cell populations per patient, by GA. MAC, macrophage; NKT, NK T cells; NK, total NK cells. **c.** Representative cell phenotype map of immune composition in decidual tissue in an early (6 weeks GA) sample. Green = NK, blue = macrophage, pink = T cell, purple = EVT, grey = other. **d.** Inset of **c**, showing a MIBI color overlay of NK cells with GrB expression. Grey = H3, yellow = CD56, blue = CD11c, red = GrB. **e.** Inset of **c**, showing a MIBI color overlay of T cells with PD-1 and Lck expression. Grey = H3, pink = CD3, orange = Lck, blue = PD-1. **f.** Representative cell phenotype map of immune composition in decidual tissue in a late (16 weeks GA) sample. Green = NK, blue = macrophage, pink = T cell, purple = EVT, grey = other. **g.** Inset of **f**, showing a MIBI color overlay of EVTs (1a, 1b) with PD-L1 expression. Grey = H3, purple = HLA-G, yellow = CD56, green = PD-L1. **h.** Inset of **f**, showing a MIBI color overlay of macrophages with TIM-3 expression. Grey = H3, green = DC-SIGN, blue = CD163, red = TIM-3. **i.** Predicted versus actual GA in days for a ridge regression model trained on GA-associated immune features, for a withheld test set (30%). Shaded region; 1 standard deviation. **j.** Ridge regression model coefficient loadings for GA-associated immune features. **k.** Pairwise spatial enrichment relationships including those temporally coordinated with GA or GA/SAR synchronized. Trend size for coordination with GA or GA/SAR synchronized, where a more positive trend (red) indicates increasing spatial enrichment with GA or GA/SAR synchronized, and a more negative trend (blue) indicates decreasing spatial enrichment between cell types. Arrow length represents mean spatial enrichment (Z-score) for cell pairs across the entire dataset. Circle size represents the number of coordinated pairwise relationships for each given cell type.

Using computational approaches validated in previous work for identifying statistically significant enrichment of two cell types (see Methods)^47–49^, we observed that the majority of significant pairwise enrichments involved EVT, NK cells, and macrophages (Fig. 4k, Supplementary Table 6). Once again, by examining these relationships on a per image basis, we were able to distinguish spatial relationships that evolved dynamically with respect to GA (see Methods). Of these relationships, the pregnancy-specific Mac2a population was involved in the largest number of pairwise enrichments, becoming more enriched around several NK and EVT subsets. Interestingly, pairwise interactions between Mac2a and CD11c+ NK cells (NK3) become increasingly prevalent with GA (Fig. 4k), even though NK cells as a whole are in decline (Fig. 4b-j).

### Coordinated up-regulation of tolerogenic functional markers with GA

Having examined the influence of GA and SAR in driving changes in the frequency of cell populations in the decidua, we next employed a similar approach to understand how these two time axes correlate with shifts in decidual function. Using the same method as our analysis of cell frequencies, we classified the temporal dynamics of functional marker expression as GA-driven, SAR-driven, or synchronized (comparably correlated with both GA and SAR) (Fig. 3n). Out of the 48-cell population-functional marker combinations that were evaluated, 16 exhibited functional marker expression that significantly correlated with one or both axes (Fig. 5a, see Methods).

**Figure 5.**
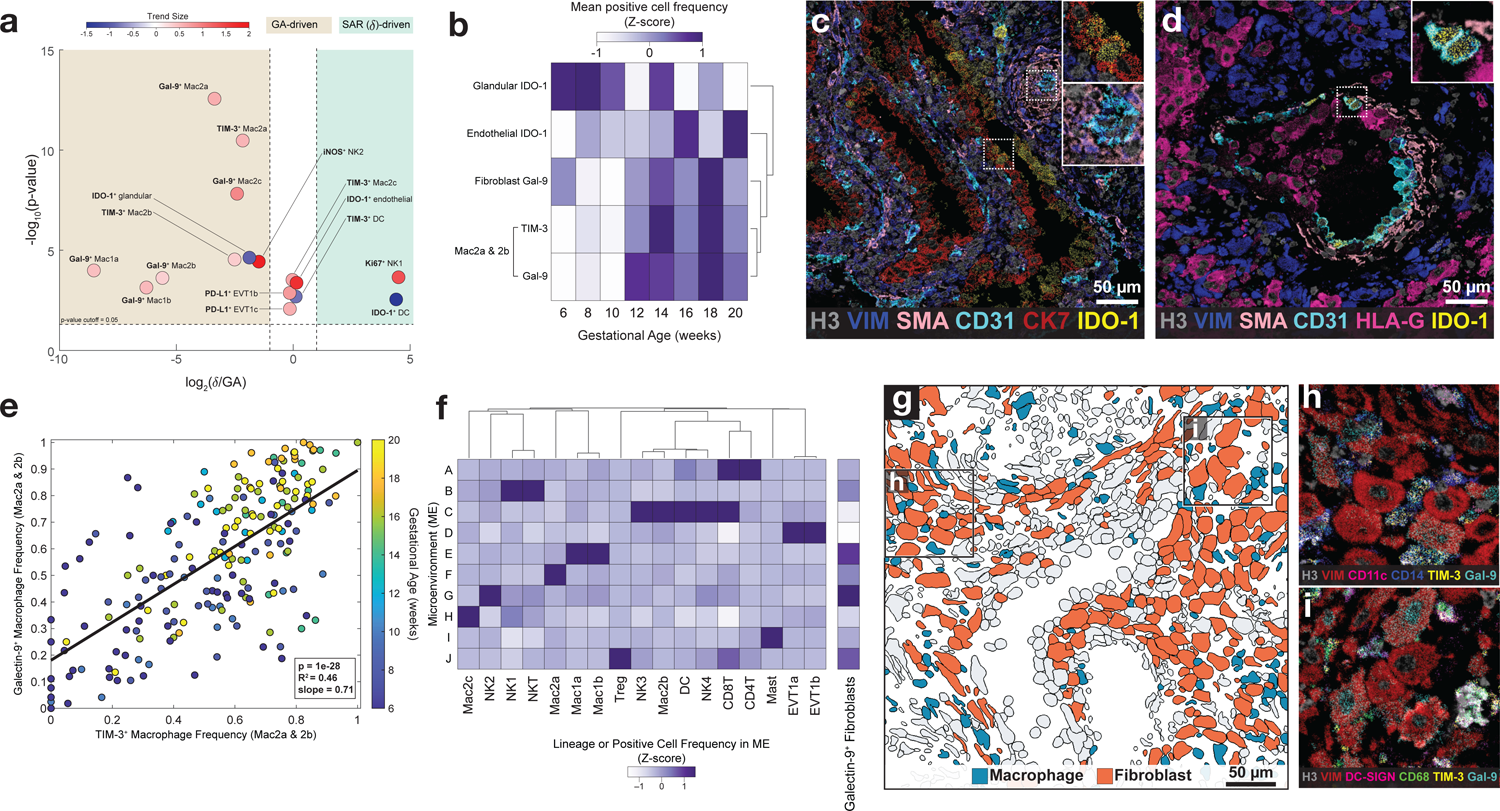
I Coordinated up-regulation of tolerogenic functional markers with gestational age (GA). **a.** Volcano plot distinguishing GA-driven from SAR (δ)-driven cell type-specific functional marker positivity fraction. X axis: log_2_ ratio of trend size is a relative measurement of R^2^ derived from linear regression against GA or SAR (δ) and GA. Y axis: -log_10_ of the p-value for the better-fitting regression model. Points are color coded by the trend size observed in the better-fitting regression model **b.** Heatmap of changes in a subset of GA-driven functional markers as a function of GA in weeks. GA in days is binned to weeks for visualization. **c.** MIBI color overlay of IDO-1 expression in glandular cells (top inset) and endothelial cells (bottom inset) in an early (6 weeks GA) sample. Grey = H3, blue = VIM (vimentin), peach = SMA (smooth muscle actin), cyan = CD31, red = CK7, yellow = IDO-1. **d.** MIBI color overlay of IDO-1 expression in endothelial cells (inset) in spiral artery (SAR manual Stage 4) of a late (16 weeks GA) sample. Grey = H3, blue = vimentin, peach = SMA, cyan = CD31, magenta = HLA-G, yellow = IDO-1. **e.** Per-image Mac2a and Mac2b TIM-3^+^ cell frequency versus Mac2a and Mac2b galectin-9^+^ frequency, colored by GA. **f.** Lineage composition of cellular microenvironments (ME) across the cohort, and frequency of Galectin-9^+^ fibroblasts in each ME. **g.** Cell phenotype map of macrophages and decidual fibroblasts. **h.** Inset of **g**; MIBI color overlay of TIM-3^+^ and galectin-9^+^ Mac2b and fibroblast cells. **i.** Inset of **g**; MIBI color overlay of TIM-3^+^ and galectin-9^+^ Mac1a, Mac1b, Mac2a, and fibroblast cells. Grey = H3, red = vimentin, pink = DC-SIGN, green = CD68, yellow = TIM-3, turquoise = Galectin-9.

These data revealed three overarching trends. First, both SAR and GA are associated with dynamic changes in IDO-1 expression. We identified a GA-driven decline in IDO-1^+^ glandular cells (Log_2_ R^2^ ratio(δ:GA) = −1.8, p-value for GA = 2.3e^-5^) in line with previous observation of IDO-1^+^ glandular cells in the first trimester but not at term^35^. Interestingly, this trend in IDO-1 glandular expression coincides with an additional immune-modulatory protein, glycodelin-A, which has been suggested to facilitate placentation^50, 51^. We’ve also observed a SAR-driven decline in IDO-1^+^ dendritic cells (Log_2_ R^2^ ratio(δ:GA) = 4.4, p-value for δ = 3e^-3^), and an increase in IDO-1^+^ vascular endothelium (p-value = 4e^-4^, Fig. 5c, d) that was comparably correlated with both GA and SAR (Fig. 5b, d). Second, consistent with the cell frequency analysis (Fig. 3n) in which NK1 exhibited a preferential increase with SAR, NK1 also exhibited a similar increase in Ki67^+^ frequency (Log_2_ R^2^ ratio(δ:GA) = 4.5, p-value for δ = 2e^-4^) becoming more proliferative as arterial remodeling progresses (Fig. 5a). Third, functional shifts in innate immunity were preferentially correlated with GA. All five macrophage populations upregulated either TIM-3 and/or its cognate ligand Galectin-9 with GA (Fig. 5a, b). This trend was most prominent in the Mac2a and Mac2b populations, where a tightly correlated up-regulation of both TIM-3 and Galectin-9 was observed (Fig. 5e, g, h, Extended Data Fig. S5a).

Interestingly, Galectin-9 upregulation was also detected in fibroblasts and found to be higher at 12-20 weeks GA (Fig. 5b, g-i). In previous work, interactions between maternal immune and stromal cell populations have been implicated in promoting fetal tolerance^52, 53^. With this in mind, we next sought to determine whether this subset was biased to colocalize within specific spatial niches. To answer this question, we quantified their frequency within 10 tissue microenvironments that were identified by clustering the cell type compositions of each cell’s closest neighbors (see Methods). Galectin 9+ fibroblasts were strongly biased to colocalize with CD57+ NK cells (NK2, Microenvironment G Fig. 5f). Notably, this trend was accompanied by GA-dependent increase in the expression of iNOS in NK2 (Fig 5a). Both TIM-3 and Galectin-9 have been implicated in suppressing anti-tumor surveillance by impairing the activity of cytotoxic NK and T cells in various human cancers^31, 54–56^. Taken with the transient perivascular enrichment of NK2s observed in early SAR, these findings suggest that expression of these proteins by macrophages and fibroblasts could play a concerted tolerizing role with fetal EVTs to attenuate immune cytotoxicity subsequent to NK-dependent early disruption of arterial smooth muscle.

### Spatio-temporal EVT distribution and transcriptional patterns suggest that intravasation is the predominant route of EVT invasion in superficial decidua

Although intravascular EVTs are known to originate from the cytotrophoblast cell columns, their path of migration remains a subject of debate primarily revolving around two models: intravasation and extravasation (Fig. 6a). In the intravasation model, EVTs detach from the cell columns and migrate through the decidua to first localize around the spiral arteries. These perivascular EVTs then enter spiral arteries by migrating through the arterial wall. In contrast, in the extravasation model, EVTs do not traverse the arterial wall from within the decidua. Instead, detaching EVTs migrate retrograde against arterial blood flow after entering at the basal plate where spiral arteries empty and merge into the intervillous space^9^.

**Figure 6.**
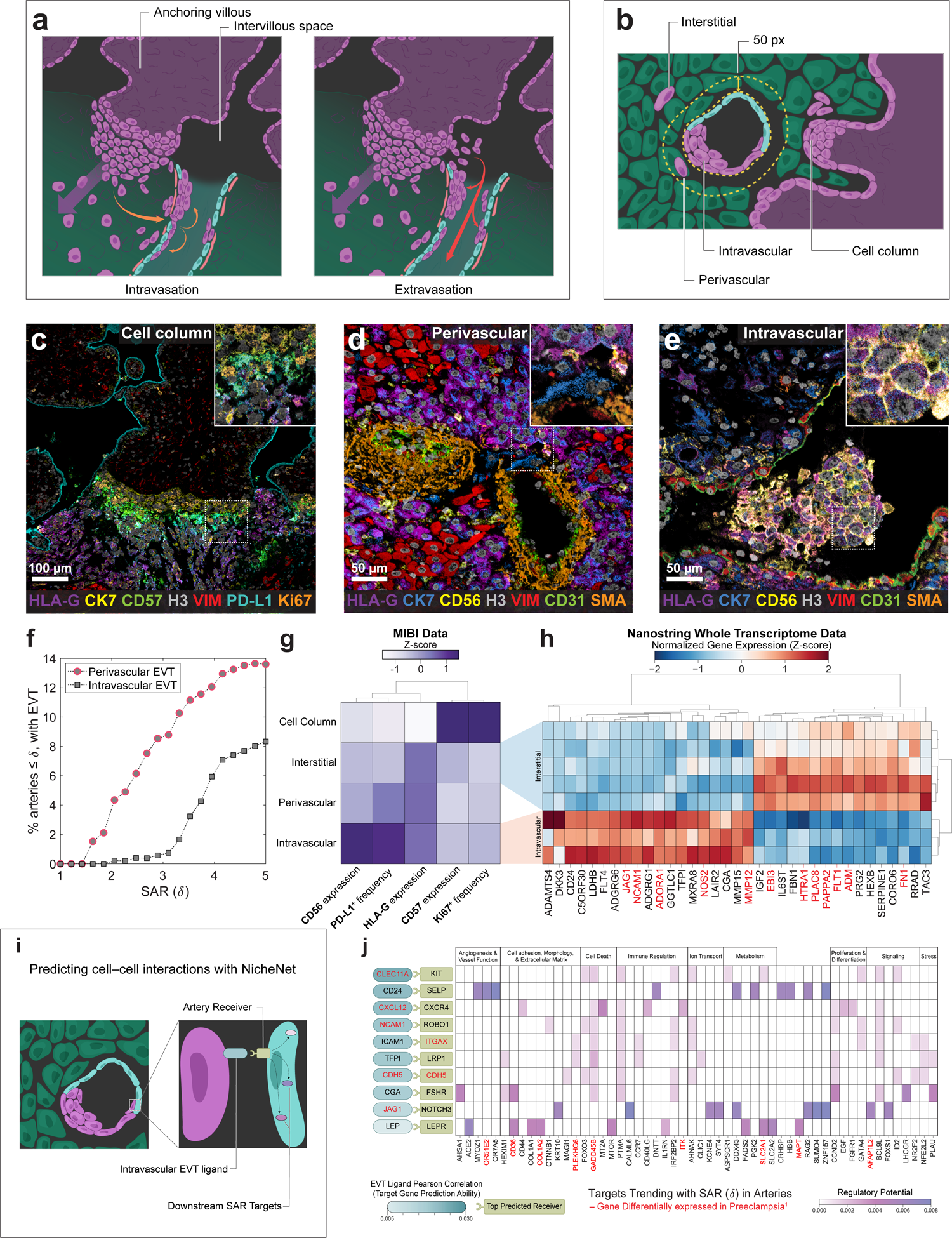
I Spatiotemporal EVT distributions suggest that intravasation is the predominant route of EVT invasion in superficial decidua. **a.** Two hypotheses for intravascular EVT invasion. (Left) Intravasation: orange arrows indicate movements of EVTs from the cell column of the anchoring villi into the decidua and through the wall of the artery and into the lumen. (Right) Extravasation: red arrows indicate movement of EVTs from the fetal villi through the intervillous space into the artery. **b.** Anatomical locations of interstitial, intravascular, perivascular, and cell column EVT populations in the decidua. **c.** MIBI overlay of anchoring villous and associated cell column EVT populations. Inset: cell column EVTs. Purple = HLA-G, yellow = CK7, green = CD57, grey = H3, red = VIM (vimentin), cyan = PD-L1, orange = Ki67 **d.** MIBI overlay of spiral arteries and associated perivascular EVT populations. Inset: perivascular EVT breaching artery wall. Purple = HLA-G, blue = CK7, yellow = CD56, grey = H3, red = VIM, green = CD31, orange = SMA (smooth muscle actin). **e.** MIBI overlay of remodeled spiral arteries and associated intravascular EVT populations. Inset; intravascular EVTs in a clump. Purple = HLA-G, blue = CK7, yellow = CD56, grey = H3, red = VIM, green = CD31, orange = SMA. **f.** Percentage of arteries with scores less than or equal to a given SAR (δ) threshold, by perivascular or intravascular EVTs present. Arteries were considered to have perivascular or intravascular EVT if the number of EVT in the appropriate artery compartment was >5. **g.** Lineage and functional marker trends of EVT populations by anatomical location. Lineage marker (CD57, HLA-G, CD56) trends are mean expression values of EVT populations. Functional marker (Ki67, PD-L1) trends are mean positive cell frequencies of EVT populations. Columns are z-scored and hierarchically clustered. **h.** Expression (Z-score) of the top 35 differentially expressed genes by logFC (adj p-value <0.05) between interstitial and intravascular EVT populations. Genes also differentially expressed in Preeclamptic decidua samples^1^ indicated in red. **i.** Application of NicheNet algorithm to artery and intravascular EVT Whole Transcriptome Data to predict EVT-artery interactions and downstream signaling targets. **j.** Outcome of NicheNet’s ligand activity prediction on differentially expressed genes on intravascular EVTs: results are shown for the 10 EVT-ligands best predicting receivers expressed in arteries, ranked by Pearson correlation coefficient or the EVT ligand activity ranking metric. Targets in arteries shown are those with at least one interaction of regulatory potential > 0.025. Ligands, receivers, and targets also differentially expressed in Preeclamptic decidua samples^1^ indicated in red.

To determine which model best explains arterial invasion, we used our spatiotemporal atlas to quantify how the phenotype and spatial distribution of EVTs evolve with respect to SAR. First, we manually defined feature masks demarcating cell column anchoring villi and three decidual compartments—interstitial, perivascular, and intravascular in our images (Fig. 6b), and then quantified EVT frequency in each (Fig. 6c-e). Together with our SAR temporal trajectory, we used these data to ask a question that has been qualitatively explored in previous work^20^: Where do EVTs accumulate first—in the perivascular compartment (directly proximal to arteries) or within the intravascular compartment? We quantified peri- and intravascular EVTs on a per-artery basis with respect to their remodeling score δ and found that perivascular EVTs began accumulating around less remodeled arteries in the decidua (Fig. 6f) and were consistently present at earlier remodeling stages than intravascular EVTs (median δ = 2.2 vs. 3.2, Kruskal-Wallis p-value = 5e^-8^, Extended Data Fig. S6a). Furthermore, out of all arteries with intravascular EVT present, 75% also had perivascular EVT-a higher percentage than would be expected if retrograde migration were the primary source of proximal intravascular EVTs. For arteries in which both perivascular and intravascular EVTs were present, the Log_2_ ratio of EVTs present in these two compartments followed a continuous and smooth trend as remodeling progressed, with intravascular EVTs increasing at the expense of perivascular EVTs (R^2^ = 0.5, p-value = 9e^-12^, Extended Data Fig. S6b). For a small number of arteries, we were able to capture perivascular EVTs breaching the artery wall, suggesting they are in the process of invading the artery lumen (Fig. 6d).

Loss of smooth muscle and endothelium have defining roles in determining the extent of SAR. Using morphometrics to quantify the extent of these concentric layers of the arterial wall (see Methods), we examined how their integrity relates to EVT enrichment in the perivascular and intravascular compartments. Similar to the trend seen with respect to remodeling score δ, accumulation of perivascular EVTs was consistently present around arteries at an earlier stage, with intravascular EVTs only appearing after 80% smooth muscle loss (median smooth muscle loss for arteries with at least five intravascular EVT present: 98%, Extended Data Fig. S6c). Perivascular EVTs were present around arteries irrespective of the degree of endothelium loss, while intravascular EVTs increased with endothelium loss (Extended Data Fig. S6d, linear regression on Log transformed intravascular EVT as a function of endothelium loss: R^2^=0.13, p-value =3e^-4^) indicating that endothelial disruption is a precursor or occurs concomitant with EVT entry into the arterial lumen. This conclusion further supports the intravasation model, in which EVTs must transverse the endothelial barrier to enter the arterial lumen.

Taken together, these data are consistent with a sequential process in which EVTs detach from the cytotrophoblast cell columns and migrate through the decidua as interstitial EVTs to accumulate in the perivascular compartment prior to intravasation, as suggested previously^20^. To further evaluate this model, we posed the following questions: Does EVT phenotype shift in a progressive manner that is consistent with this stepwise intravasation model? If so, do intravascular EVTs more closely resemble the cell column or perivascular compartment?

We found that the composition of cell column, interstitial, perivascular, and intravascular compartments shifted in a systematic manner along the proposed path of migration: cell columns consisted primarily of EVT1a, EVT1b and EVT2 subsets with few CD56^+^ EVT1c cells (99% vs. 1%), and EVT1c cells were further enriched within the perivascular compartment (7.3%) but most prevalent in the intravascular compartment (54%, Extended Data Fig. S6e). EVT1c cells were also found significantly closer to arteries than EVT2 cells (Kruskal Wallis p-value = 1.15e^-19^, Extended Data Fig. S6f). Examining phenotypic marker expression within each compartment once again revealed a progressive shift in EVT phenotype that best aligned with an intravasation model. Cell columns were uniquely enriched for proliferative (Ki67^+^), CD57^+^ EVTs (Fig. 6g), and a precipitous drop in CD57 and Ki67 expression was accompanied by a progressive increase in PD-L1 that peaked in the intravascular compartment (Fig. 6g). While functional marker expression in the perivascular compartment is most similar to the intravascular compartment (Fig. 6g, Extended Data Fig. S6g), a noticeable difference between the two compartments is driven by the prevalence of CD56^+^ PDL-1^+^ EVT1c cells in the intravascular compartment, which further increases with SAR (Extended Data Fig. S6h).

Note that given the observational nature of this study, neither model can be definitively ruled in or out. However, taken together these analyses best align with an intravasation model in which decidual invasion of cell column EVTs is accompanied by pronounced downregulation of CD57 and Ki67 and upregulation of HLA-G. Perivascular accumulation of EVTs occurs early in SAR, preceding the appearance of intravascular EVTs and any loss in endothelium. In this model, as the endothelial barrier is lost, perivascular EVTs invade the artery lumen and upregulate CD56 (see Methods, Extended Data Fig. S6i-l).

Irrespective of the route of migration, the distinct changes in phenotypic markers measured by MIBI-TOF suggest that arterial invasion is accompanied by a shift in EVT transcriptional programs. With this in mind, we used Nanostring DSP to reveal differentially expressed genes (DEGs) between interstitial and intravascular EVTs, leveraging the spatial information preserved on serial sections to specifically select and measure the transcriptomes of these subsets for the first time. We found 274 genes that were differentially expressed between interstitial and intravascular EVTs (Fig. 6h, Extended Data Fig. S6m,n, see Methods). In addition to confirming previous work noting upregulation of NCAM1, JAG1, and LAIR2 specifically in intravascular EVTs^57–59^, we identified transcriptional changes in genes important for ECM remodeling and angiogenesis (Fig. 6h). For example, MMP12, MMP15 and ADAMTS4^60–62^ were specifically upregulated in intravascular EVTs (mean logFC; MMP12: 11.67, MMP15: 9.74, ADAMTS4: 7.88), suggesting that these proteins play a significant role in late stage SAR. Additionally, arterial invasion was accompanied by a shift from VEGFR1 (FLT1) to VEGFR3 (FLT4) (Fig. 6h).

Intravascular EVTs were also found to upregulate DKK3, C5orf30 (MACIR), and CD24 (Fig. 6h) which have each been shown in previous work to play roles in fetal viability, tumor invasion, or immune tolerance^63–66^. Similarly, we observe an accompanying downregulation of genes associated with invasion in intravascular EVTs, such as MGAT5. With respect to immune modulation, C5orf30 is a potent immunometabolic regulator that has been shown to inhibit macrophage-mediated tissue damage in rheumatoid arthritis^63^. Similarly, CD24 binding to Siglec-10 was recently found in many cancers to promote immune evasion by serving as an anti-phagocytic, “don’t eat me” signal^66^. Taken together, our multi-modal approach paints a picture of a highly regulated and controlled process. We observe a transcriptional shift away from a more invasive phenotype (SERPINE1, CORO6) in interstitial EVTs towards genes implicated in vascular remodeling in intravascular EVTs. Importantly, this is accompanied by an increase in immunoregulatory modules that allow EVTs to be in continuous contact with maternal blood while avoiding immune activation (i.e. C5orf30, CD24, PD-L1) (Fig. 6g-h).

To better understand how these changes promote SAR, we investigated potential cell-cell interactions between intravascular EVTs and arterial cells using NicheNet ^67^ (Fig. 6i). This analysis identified 10 protein-protein interactions that were predicted to drive changes in 121 downstream targets (see Methods, Extended Data Fig. S6o). For example, interactions between EVT JAG1 and arterial NOTCH were predicted to drive downstream changes in arterial MEOX1 and MT2A, which have been implicated in endothelial dysfunction and apoptosis^68, 69^. Similarly, CGA-FSHR and LEP-LEPR interactions were correlated with changes in arterial hormone receptors (LHCGR) and several cell adhesion targets, respectively (Fig. 6j, Extended Data Fig. S6o, Supplementary Table 7). Intriguingly, among the most prominent downstream targets were olfactory receptors: OR51E2 and the human specific OR7A5, expression of which outside of the olfactory bulb has been thought to regulate blood pressure and angiogenesis^70, 71^.

CD24-SELP was the second most significant interaction and had several targets related to blood vessel function and formation (Fig. 6j, Extended Data Fig. S6o, Supplementary Table 7). Interestingly, reduced global placental CD24 expression has been associated with an increased risk of developing preterm preeclampsia, yet EVT-specific expression of CD24 transcripts has not yet been reported^65, 72^. Given that abnormal decidual and spiral artery remodeling are thought to play a major role in preeclampsia^73^, we sought to determine if other genes involved in EVT invasion and vascular remodeling had previously been implicated as well.

To do this, we first compared our list of EVT DEGs with genes found previously to be differentially expressed in decidua samples from women diagnosed with preeclampsia^74^. We found that 31% of EVT DEGs (12 genes) are differentially expressed in preeclamptic decidua^75–78^ (Fig. 6h, Supplementary Table 8). Notably, FN1 and FLT1, which have been proposed as biomarkers for early prediction of preeclampsia^79, 80^,were markedly downregulated in intravascular EVTs.

Similarly, half of the NichNet interactions and 19 downstream targets overlapped with this list of preeclampsia DEGs^74^ (Fig. 6j, Extended Data Fig. S6o). These included WNT10B—a novel accelerator of EVT invasion^81^ (Extended Data Fig. S6o)—and OR51E2, a target of CD24-SELP signaling that also exhibited the highest regulatory potential. With respect to the latter, SELP is notable for being differentially expressed in peripheral blood cell free RNA from patients with preeclampsia^82^. Taken together, our transcriptomic approach validates and compliments the stepwise changes in EVT phenotype seen in our spatial atlas, as well as highlighting signaling pathways that could serve as novel candidates for understanding how the functional dynamics of EVT migration and invasion are perturbed in pregnancy-related disorders.

## Discussion

Decidualization is a fascinating process with no other normative precedent in human biology, where the structure and function of the maternal endometrium transforms to promote tightly regulated invasion of genetically dissimilar placental cells. In this process, the decidua plays a dual role by permitting EVT invasion in the first trimester and later limiting it by inducing EVT apoptosis^83^. EVT invasion can also be limited by morphological changes such as EVT fusion leading to polyploidization which limits invasion due to nuclear size^84^. Given the lack of tractable and relevant animal models and the inability to study decidualization prospectively, our understanding of it is immature relative to other areas of human physiology. With this in mind, we used MIBI-TOF and Nanostring DSP on archival human tissue to generate the first multimodal, spatiotemporal atlas of the maternal-fetal interface during 6-20 weeks gestation. The central focus of our study was to understand how global, temporally dependent changes in decidual composition are coupled to local regulation of vascular remodeling in pregnancy. While initial invasion of placental EVTs is prompted by a shift towards a permissive milieu, progression of SAR is dependent on subsequent migration and perivascular accumulation of EVTs, where they are thought to participate in cooperative cell-cell interactions with maternal fibroblasts, NK cells, and macrophages ^4,5^. Thus, formation of the maternal-fetal interface is mediated by global, temporally dependent cues that serve as a gating function for remodeling processes that are regulated in the local tissue microenvironment.

With this paradigm in mind, we set out to delineate which aspects of the first half of pregnancy are driven globally by GA and how this relates to SAR. To achieve this, we mapped the spatial distribution, composition, and functional state of ∼500,000 maternal cells and fetal EVTs with respect to glands, anchoring cell column villi, and spiral arteries in >200 images from the decidua of 66 patients. In studying placentation and SAR, an ideal sampling strategy might use elective cesarian hysterectomies from normal pregnancies performed across GA in an ethnically diverse patient population. As ethical considerations prohibit this, previous work has employed a range of sample types that each have their own strengths and weaknesses. Here, we utilized archival tissue from elective terminations of uncomplicated pregnancies to examine these questions in a large, ethnically diverse patient cohort that is well distributed with respect to GA. Since tissue procured during terminations is fragmented, anatomic registration for determining if these tissue blocks were sampled from central or peripheral regions of the decidua basalis was not feasible.

Using LDA, image morphometrics, and expert annotations, we assigned quantitative remodeling scores to every spiral artery in these images. These targeted multiplexed imaging data were complimented by spatially coregistered tissue transcriptomics. This multi-modal dataset permitted us to reveal how cell frequency and function, tissue organization and transcriptional programs in maternal decidua, arteries, and EVTs change with SAR and GA.

Our analysis of these changes determined GA to be the predominant driver of maternal immune cell composition (Figs. 3n, 4i, j). Progressive decreases in NK and T cells drive a transition at 12-14 weeks GA from a lymphoid to myeloid predominant decidua enriched for iNOS^+^ NK cells, IDO-1^+^ vascular endothelium, and DC-SIGN^+^ macrophages that co-express TIM-3 and Galectin-9 (Figs. 4b, 5a, b). Notably, this relationship between immune composition and GA was robust enough to allow us to predict GA within 19 days based exclusively on immune population frequencies (Fig. 4i).

In contrast, all EVT subsets and only two maternal cell populations (NK1 and NK2) preferentially correlated with progression of SAR. Higher remodeling scores were correlated with more EVTs, more NK1s, and fewer NK2s. NK2s were preferentially depleted from the perivascular region, after initial accumulation around lowly remodeled arteries. NK1 and NK2 primarily differ in that the latter express CD57—a marker associated with a cytotoxic phenotype. Higher proportions of presumptively more reactive NK2s early in SAR around arteries aligns well with previous studies that have suggested that decidual NKs initiate early disruption of arterial smooth muscle through secretion of GrB, MMP2, and MMP9^26, 85^. Likewise, the proportional gains seen here as SAR progresses of less reactive NK1s and invasive EVTs are consistent with the tolerizing effects of HLA-G, which has been shown previously to decrease NK cell cytotoxicity and induce production of IL-6 and IL-8 via binding of HLA-G to KIR2DL4, LILRB1, and LILRB2^86, 87^. Taken together, these data suggest that maternal and fetal cells play cooperative, interdependent roles with SAR transitioning through NK- and EVT-dependent phases.

We also examined a long-standing open question^9^ in the field: What is the predominant path of migration taken by EVTs that invade spiral arteries? In line with early work based on studies of 8-18 week hysterectomy specimens processed in toto^9,88^, we believe our work using material from elective terminations provides compelling evidence for EVT intravasation of arteries in the decidua basalis. On comparing cellular composition within cytotrophoblast cell columns of anchoring villi, decidua, and arteries, we found EVT frequency and phenotype to shift in a sequential, coordinated manner consistent with an intravasation model where EVTs within the decidua enter spiral arteries through the arterial wall. Notably, previous studies of cesarian hysterectomies also found morphologic evidence of arterial extravasation. Given the observational nature of this study, we cannot rule out an extravasation model in which EVTs migrate in a retrograde manner after entering spiral arteries directly at the basal plate. We also note the possibility that upon intravasating arteries in the decidua, EVTs could migrate upstream within the artery to reach the upper third of the myometrium. This would be consistent with previous studies where perivascular trophoblasts were found to become increasingly scarce as a function of myometrial depth^89–92^. With this limitation in mind, in our model EVTs detaching from proliferative cytotrophoblast cell columns first invade the decidua and transition to a CD57^-^ CK7^+^ HLA-G^+^ phenotype in our proposed model. In line with previous work demonstrating EVT expression of MMP2 and MMP9^93^, these cells migrate through the decidua and accumulate around spiral arteries where they participate in removal of arterial smooth muscle. As this layer is depleted, perivascular EVTs disrupt the underlying vascular endothelium and invade the arterial lumen where they form multicellular clumps. Intravascular invasion is accompanied by EVT upregulation of CD56, a homophilic binding molecule that has been suggested to be necessary for heterotypic cell adhesion to endothelial cells^94^. Finally, these multicellular clumps in fully remodeled arteries recede and are replaced by EVTs that have partially displaced the maternal endothelium.

While previous single cell and bulk sequencing studies of decidua have characterized the transcriptome of decidual cells in great detail, they were performed on dissociated tissue, agnostic to spatial context and the local extent of SAR. Correlating spatial morphology and tissue composition with targeted tissue transcriptomics in coregistered, serially sectioned tissue, allowed us to observe for the first time how the transcriptome evolves with respect to SAR. In arteries, our analysis revealed a down-regulation with SAR of Notch signaling, tissue organization and cohesion, accompanied by a burst of translation-related activity around stage 2 of remodeling. In comparing interstitial with intravascular EVTs, our analyses revealed genes upregulated in the interstitial populations that shed light on novel aspects on how EVTs contribute to facilitating immune tolerance. We observe a compelling upregulation of genes typically associated with cancer progression in intravascular EVTs, such as DKK3, CD24, ADORA1, and JAG1, suggesting that there may be similarities between mechanisms of immune evasion in tumors and at the maternal-fetal interface. Almost a third of DEGs between interstitial and intravascular EVTs overlap with differentially expressed genes in preeclamptic decidua samples. Given the significant contribution abnormal vascular remodeling and EVT invasion are thought to play in preeclampsia, this work should serve as a valuable community resource that can be used to contextualize preeclampsia-related changes for future studies.

Formation of the maternal-fetal interface is an organized and controlled invasive process that is sometimes viewed as a template for understanding invasive and immunosuppressive properties of tumors^95^. Both processes involve a genetically dissimilar invasive cell type (haploidentical EVTs vs. clonal, mutated cancer cells), extracellular matrix remodeling, and recruitment of a wide variety of tolerogenic immune cells, including M2 polarized macrophages and proliferating Tregs. The intersection of anchoring placental villi and maternal decidua morphologically resembles the invasive margin of carcinomas and contains trophoblast cells expressing high levels of immunomodulatory proteins and growth factors implicated in tumor severity including PD-L1, IDO-1, TIM3, Her2, and EGFR^31, 54, 96, 97^. In addition to these phenotypic and structural similarities, recent work revealing mosaicism and clonal mutations in normal term placentas demonstrate that this phenotypic overlap is even manifest at a genomic level^98^.

Overall, we anticipate that this spatio-temporal atlas of the early human maternal-fetal interface will provide a normative framework for elucidating etiological perturbations in maternal-fetal tolerance and SAR in pregnancy complications. Likewise, this work may also serve as a template for understanding how immune tolerance, tissue remodeling, and angiogenesis are aberrantly recruited and synergized during tumor progression. With this in mind, we plan in future studies to extend this comparative approach to archival tissue from patients with preeclampsia, placenta accreta, and choriocarcinoma to further elucidate cellular interactions involved in regulating SAR and EVT invasion.

## Methods

### Retrospective cohort design

The study cohort comprised decidua tissue from archival formalin-fixed, paraffin embedded (FFPE) blocks, sampled after elective pregnancy terminations at the Women Options Center at Zuckerberg San Francisco General Hospital, an outpatient clinic located within a large public hospital affiliated with an academic medical center. Patients at this clinic reflect a diverse population. The clinic serves women in the Bay Area as well as referrals from California and out of state. While the patient population is predominantly low-income mainly Medi-Cal patients, women of all economic backgrounds are cared for at the clinic.

In the clinic, an ultrasound examination is performed to estimate GA, and medical history is taken and logged as Electronic Medical Record– (‘eCW’ - electronic clinical works) or handwritten forms. A board-certified gynecologist reviewed medical records and specifically extracted the following details: age, ethnicity, body mass index, gravidity, parity, prior terminations, smoking, medications, HIV status, history of preeclampsia, chronic hypertension, diabetes mellitus, renal disease, autoimmune disease, multifetal pregnancy, and congenital anomalies (Supplementary Table 1). For procedures occurring at less <14 weeks GA, suction aspiration is routinely used. For procedures at >14 weeks GA, a combination of suction aspiration and grasping forceps is used. After the procedure, tissue samples are routinely sent to pathology.

### TMA construction

Whole tissue sections from patients who underwent elective termination at San Francisco General Hospital at 6-20 weeks gestation were first reviewed by H&E staining to identify samples containing decidual tissue and spiral arteries. These regions were manually demarcated and assessed for suitability, blocks containing decidua with vessels were selected, cored with a bore needle, and assembled into the TMA used in this study.

Archival tissue blocks from 74 patients were initially selected by a board-certified perinatal pathologist (G.R.) to be included in the TMAs. The first TMA consisted of 205 cores (including 3 tonsil cores, 1 endometrium core, 1 myometrium core) of 1 mm in diameter and the second contained 86 cores of 1.5 mm in diameter). Unfortunately, cores from 8 patients did not end up containing decidua, and there was not sufficient tissue in the block for additional re-coring. We therefore had to exclude these patients from the analysis. The final cohort included 66 patients, an exhaustive list of which is provided in Supplementary Table 1. Images from 6 patients had no arteries and therefore were not included in analyses related to spiral arteries. Information on the histological characteristics of the blocks retrieved, including the presence of cell column anchoring villi, is in Supplementary Table 2. High resolution scans of each core were uploaded to the Stanford Tissue Microarray Database (URL: http://tma.im/cgi-bin/home.pl), a collaborative internal platform for designing, viewing, scoring, and analyzing TMAs. Sequential recuts of the main experiment were stained with H&E, to aid in choosing the imaging regions of interest (ROIs) and analyzing data.

### Antibody preparation

Antibody staining was validated as described previously^29, 47–49, 99^. Briefly, each reagent was first tested using single plex chromogenic IHC using multiple positive and negative FFPE tissue controls prior to metal conjugation. Antibodies were then conjugated to isotopic metal reporters as described previously^29, 47–49, 99^ with the exception of biotin-conjugated anti-PD-L1, for which a metal-conjugated secondary antibody was used. Performance of metal conjugated antibody reagents were then tested within the complete MIBI-TOF staining panel, under conditions identical to those in the main study and compared with representative single plex chromogenic IHC to confirm equivalent performance. Representative stains and information for each marker can be found in Extended Data Fig. S1 and Supplementary Table 9 respectively. After conjugation, antibodies were diluted in Candor PBS Antibody Stabilization solution (Candor Bioscience). Antibodies were either stored at 4°C or lyophilized in 100 mM D-(+)-Trehalose dehydrate (Sigma Aldrich) with ultrapure distilled H_2_O for storage at −20°C. Before staining, lyophilized antibodies were reconstituted in a buffer of Tris (Thermo Fisher Scientific), sodium azide (Sigma Aldrich), ultrapure water (Thermo Fisher Scientific), and antibody stabilizer (Candor Bioscience) to a concentration of 0.05 mg/mL. Information on the antibodies, metal reporters, and staining concentrations is in Supplementary Table 9.

### Tissue staining

Tissues were sectioned (4 μm in thickness) from tissue blocks on gold and tantalum-sputtered microscope slides. Slides were baked at 70°C for 20 minutes followed by deparaffinization and rehydration with washes in xylene (3x), 100% ethanol (2x), 95% ethanol (2x), 80% ethanol (1x), 70% ethanol (1x), and ddH_2_O with a Leica ST4020 Linear Stainer (Leica Biosystems). Tissues next underwent antigen retrieval was carried out by submerging sides in 3-in-1 Target Retrieval Solution (pH 9, DAKO Agilent) and incubating them at 97°C for 40 minutes in a Lab Vision PT Module (Thermo Fisher Scientific). After cooling to room temperature slides were washed in 1x PBS IHC Washer Buffer with Tween 20 (Cell Marque) with 0.1% (w/v) bovine serum albumin (Thermo Fisher). Next, all tissues underwent two rounds of blocking, the first to block endogenous biotin and avidin with an Avidin/Biotin Blocking Kit (Biolegend). Tissues were then washed with wash buffer and blocked for 1 hour at room temperature with 1x TBS IHC Wash Buffer with Tween 20 with 3% (v/v) normal donkey serum (Sigma-Aldrich), 0.1% (v/v) cold fish skin gelatin (Sigma Aldrich), 0.1% (v/v) Triton X-100, and 0.05% (v/v) Sodium Azide. The first antibody cocktail was prepared in 1x TBS IHC Wash Buffer with Tween 20 with 3% (v/v) normal donkey serum (Sigma-Aldrich) and filtered through a 0.1 μm centrifugal filter (Millipore) prior to incubation with tissue overnight at 4°C in a humidity chamber. After the overnight incubation slides were washed for 2 minutes in wash buffer. The second day, the antibody cocktail was prepared as described (Supplementary Table 9) and incubated with the tissues for 1 hour at 4°C in a humidity chamber. After staining, slides were washed twice for 5 minutes in wash buffer and fixed in a solution of 2% glutaraldehyde (Electron Microscopy Sciences) solution in low-barium PBS for 5 minutes. Slides were washed in low-barium PBS for 20 seconds then, using a linear stainer, through 0.1 M Tris at pH 8.5 (3x), ddH2O (2x), and then dehydrated by washing in 70% ethanol (1x), 80% ethanol (1x), 95% ethanol (2x), and 100% ethanol (2x). Slides were dried under vacuum prior to imaging.

### MIBI-TOF imaging

Imaging was performed using a custom MIBI-TOF instrument with a Xe+ primary ion source, as described previously^29, 47^. 222 808 x 808um Fields of View (FOVs) were acquired at approximately 600 nm resolution using an ion dose of 7nA*hr/mm^2^. After excluding 11 FOVs that contained necrotic or non-decidual tissue, or consisted of duplicate tissue regions, the final dataset consisted of 211 FOVs from 66 patients.

### Low-level image processing

Multiplexed image sets were extracted, slide background-subtracted, denoised, and aggregate filtered as previously described^36, 47–49^. For several markers, a “background” channel consisting of signal from the mass 128 channel was used. All parameters used as inputs for low-level processing are listed in Supplementary Table 9.

### Feature annotation

Large tissue features were manually annotated in collaboration with a perinatal pathologist. Pseudo-colored MIBI images with H3 to identify cell nuclei, vimentin for decidual stromal cells, smooth muscle actin and CD31 for vessels, cytokeratin 7 (CK7) for glands and the fetal cell columns, and HLA-G for EVTs were used to guide annotation. Serial H&E sections, and an H&E recut of the entire block, if necessary, were additionally used to supplement annotation. Labelling was performed in ImageJ and the annotated features were exported as binary TIF masks.

### Single cell segmentation

The Mesmer segmentation algorithm^37^ was adapted specifically to segment the cells in our dataset. First, training data were generated using a subset of 15 images out of 211 in our cohort, in addition to 10 decidua MIBI-TOF images from titration data. 1024 x 1024 pixel crops were selected to encompass the range of different cell morphologies present. The markers H3, vimentin, HLA-G, CD3, CD14 and CD56 were used to capture the major cell lineages present. Subsequently, a team of annotators parsed these images to identify the location of each unique cell using DeepCell Label, custom annotation software specifically developed for this task^37^ (code URL: https://github.com/vanvalenlab/deepcell-label). The manually annotated images were used to generate partially overlapping crops of 256 x 256 pixels from each image. In total, training data included 1600 distinct crops with 93,000 cells. This dataset was used to retrain the Mesmer segmentation model, modifying the architecture to accept six distinct channels of input. The output from the network was then post-processed using the default model settings (Extended Data Fig. S2a).

### Segmentation post-processing

Examination of the images revealed that glandular cells and chorionic villus trophoblasts did not express any markers included in the training data; namely these cells were predominantly CK7^+^. This resulted in effectively nuclear-only segmentation being predicted by the CNN within these features. To account for this, segmented cells that overlapped with the gland mask were expanded radially by 5 pixels, and those in the cell column mask by 2 pixels. This approach accounted for glandular cells and cell column anchoring trophoblasts that were not expressing any markers but were included in the training data, resulting in effectively nuclear-only segmentation being predicted by the convolutional neural network. The number of pixels used for expansion was optimized to approximate the observed cell size, based on systematic inspection of three images per GA. Objects <100 pixels in area were deemed noncellular and excluded from subsequent analyses. Finall number of segmented events per FOV can be found in Supplementary Table 10.

### Single-cell phenotyping and composition

Single cell expression data were extracted for all cell objects and area-normalized. Single-cell data were linearly scaled with a scaling factor of 100 and ArcSinh-transformed with a co-factor of 5. All mass channels were normalized ^to^ the 99th percentile. To assign decidual cell populations (≥ 70% cell area in decidua) to a lineage, the clustering algorithm FlowSOM (Bioconductor “FlowSOM” package in R)^38^ was used, which separated cells into 100 clusters based on the expression of 19 canonical lineage defining markers (Extended Data Fig. S2b). Clusters were further classified into 21 cell populations, with proper lineage assignments ensured by manual examination of overlayed FlowSOM cluster identity with lineage-specific markers. Clusters containing non biologically meaningful or distinct signals were assigned the label ‘other’. Tregs were identified by thresholding T cells (FlowSOM clusters 43, 53, 63) with CD3 signal ≥ the mean CD3 expression of CD4^+^ T cells and > 0.5 normalized expression of FOXP3. Mast cells were identified as cells for which normalized expression of tryptase was >0.9. Mac2b (CD11c^+^) cells were identified as macrophages with >0.5 normalized expression of CD11c. Placental macrophages (Hofbauer cells) were defined as CD14^+^ >0.5 cells located within the cell column. Cells from FlowSOM clusters 4, 5, and 15 ubiquitously and predominantly expressed CK7 and were reassigned to the EVT2 subset if located within the cell column feature mask, or as glandular cells otherwise (Extended Data Fig. S2b). These thresholds were selected based on the distribution of lineage marker expression (Extended Data Fig. S2d) as well as on systematic examination of the images by eye since expression patterns varied significantly between markers. For a comprehensive list of all single cells, their morphological features, markers expression, lineage classification et cetera – see Data Availability Statement.

### Definition of thresholds for functional marker positivity

Cells were considered positive for a functional marker if their scaled expression level was ≥ a set threshold, as described previously^47^. Thresholds for individual functional markers were determined based on examining the images by eye, as expression patterns varied significantly between markers (Supplementary Table 11, Extended Data Fig. S2h). To set the per marker thresholds, 5 images for each functional marker were reviewed and increasing threshold values were examined using custom software. Subsequently, cells defined as negative for a marker based on the determined threshold value were re-examined to ensure the thresholds were representative. For Ki67 positivity, only cells that had a nucleus in the image were considered. Ki67 values were not cell size normalized because the Ki67 signal is exclusive to nuclei.

### Blinded manual artery staging

Arteries were categorized into 5 remodeling stages based on criteria adapted from the 4-stage model proposed by Smith et al^45^. These criteria were previously used to describe spiral arteries observed in H&E and single channel IHC images and were adapted to suit multiplexed MIBI data (Fig. 3a, details in Extended data Fig. S3a). 600 arteries were categorized according to these criteria by a single reviewer using exclusively crops of MIBI pseudocolor overlays (SMA, Vimentin, CD31, H3, and HLA-G) including only the artery (as defined by feature mask) and any EVTs in the lumen. The reviewer was blinded to the rest of the image, serial H&E sections, gestational age, and any clinical data. 12 partially captured arteries were excluded from the final dataset of 588 arteries.

### Automated digitization of artery morphological features

The same format of cropped artery MIBI images that were manually scored by the reviewer were used to calculate a set of geometric parameters for several selected features. These features described the organization and structure of the vessel wall, the continuity of the endothelium and its thickness, and the presence and structure of intravascular EVTs. In order to capture these features, a structure of concentric circles we termed the “onion” structure is defined, with the outer circle of this structure enclosing the artery and the inner circles dividing it into layers. This structure is described below using the two-dimensional cylindrical coordinate system with the radial axis r, azimuthal (angular) axis ø, and origin of the axis at point (x,y). Point (x,y) is the user defined artery center. For an artery in the binary mask M, the following algorithm was used to create the “onion” structure (Extended Data Fig. S3c):

- Define a circle enclosing the artery, centered at point (x,y) with radius a as follows:

- (x,y) was taken as the user-defined artery center point
- a, the radius is defined as the maximum distance between (x,y) and the edge of M – rounded up to the nearest integer multiple of n, such that a=I*n for an integer I. n is a user defined thickness parameter for the “onion” layers.
- Define the inner circles comprising the “onion” layers:

- Divide the radius a of the outer circle into I equal sections of length n, creating layers along the radial r axis.
- The radii of the inner circles are then defined as: 0,1*n,2*n,…(I-1)*n.
- Divide the “onion” into k equal sectors along the ø axis, k is a user defined integer.
- Subdivide each sector into segments:

- The sectors are internally divided by the circles, creating parts with 4 corners and 4 sides, with the 2 sides being straight (sector dividers), and the 2 sides being arcs (parts of ellipse circumferences).
- The arcs are replaced with secants (straight line connecting the ends of the arc), turning the segment into a trapezoid.
- The parameters n=10 pixels and k=100 were used to allow for segments large enough to contain a sufficient number of pixels to average expression over.

The following features are then extracted for each artery “onion”:

1. Geometrical features:

a. radius - the maximum distance between any pixel within the mask and the closest pixel on the edge of the mask.
b. perimeter – the Euclidean distance between all adjacent pixels on the edge of the artery mask
c. area – the total number of pixels within the artery mask
2. Protein morphology features, for each of the following markers: CD31, CK7, H3, HLA-G, SMA, VIM

a. Average signal – weighted-average over segments of marker expression, where the weight of a segment corresponds to the number of pixels it contains. Weighted average was used to avoid smaller inner segments having disproportionate effect on the average.
b. Thickness –

i. For each sector we calculate the distance d between the inner-most segment positive for the marker and the outer-most positive segment. Positivity is measured by comparing the mean signal over pixels the segment to a user defined threshold.
ii. The mean and standard deviation of thickness are calculated as the mean and standard deviation of d over all sectors.
c. Radial coverage - the percentage of sectors positive for marker signal. A sector is considered positive if the mean signal over sector pixels accedes a user defined threshold.
d. Jaggedness – This feature measures the extent jaggedness of an artery outline. To do so, first a skeletonization function written by Nicholas R. Howe ^100^ is applied to the artery mask, this function returns a “skeleton” of the artery outline. This “skeleton” also assigns values to the outline pixels based on their distance from the core shape. Then, two different binarization thresholds are chosen: a “non-branch” threshold (a high value = 60 pixels, indicating greater topological distance and a “branch” threshold (a low value = 5 pixels, indicating smaller topological distance). The ratio between the total number of “non branch” and “branch” pixels is the jaggedness.

### Calculation of continuous SAR remodeling score δ

A supervised dimensionality reduction technique based on linear discriminant analysis (LDA)^46^ (code URL: https://github.com/davidrglass) was employed using the per artery digitized morphological features and manually assigned remodeling stage labels as inputs. All artery morphology feature values were standardized (mean subtracted and divided by the standard deviation) and all arteries were used as training data. The LDA output was:

a. The optimal linear combination of a subset of features, that maximized the separation by manual stage between arteries in LDA space (Supplementary Table 12)
b. The coordinates of each artery in LDA space (Supplementary Table 3)

In order to define the SAR trajectory, a fourth-degree polynomial was fitted to the artery coordinates in LDA space. To determine the optimal degree of the polynomial, polynomials with degrees 1-6 were fitted and the degree that minimized the p-value for separating δ distributions between arteries grouped by manual remodeling stage (Extended Data Fig. S3g) was selected. The polynomial fit was implemented using the MATLAB function fit and resulted in the following polynomial: f(x) = 0.0005*x^4^ −0.01227*x^3^ + 0.1363*x^2^ −0.4354*x −0.7425. The polynomial was then numerically interpolated on a dense 10^4^ point grid and the distance from each artery point in LDA space to the polynomial was calculated using this grid and the MATLAB exchange function distance2curve ^101^. δ per artery was then calculated as the line integral from the curve origin to closest point to the artery on the curve (Extended Data Fig. S3e, inset). This integral was numerically calculated using a custom MATLAB script. δ values were linearly rescaled to the range 1-5 using the MATLAB function rescale.

### Cell type frequency as function of GA and SAR

For examining cell type frequencies within the decidua as function of GA and SAR (Fig. 3, Fig. 4), per image cell frequency tables were constructed in which cell type frequencies were calculated as the proportion of cells in the decidua feature mask of that image. Cells located in other feature masks (artery, gland, vessel, or cell column masks) were not counted, nor were cells of an unassigned type (‘other’). In order to focus these analyses on cell populations strictly found in the decidua, muscle and glandular cells were also excluded; these cell types occasionally extended outside of their artery and gland feature masks, respectively. Cell frequency as a function of GA for a cell type was defined as the per image proportion values for that cell type, as function of the GAs associated with the images. Similarly, cell frequency as a function of SAR for a cell type was defined as the per image proportions of that cell type, as function of the mean δ values per image. For the volcano plot in Fig. 3n, we fitted a linear regression model to the two above-described functions. All linear regression models were implemented using the MATLAB function fitlm and the volcano plot only shows points for which regression R^2^ ≥ 0.05. R^2^ and p-values for all δ and GA based regressions can be found in Supplementary Table 13. The ratio between R^2^ in the two regression models was used to classify trends as GA-driven, SAR-driven or synchronized. For example, the increase in EVTs out of all cells, R_EVT, was classified as GA-driven because R^2^ for R_EVT as a function of δ was 0.3, but only 0.1 for R_EVT as a function of GA (Extended Data Fig. S3h, Supplementary Table 13). Another example is the increase in macrophages out of immune cells, I_sumMac: it was classified as GA-driven since R^2^ for I_sumMac as a function of GA was 0.6 but only 0.1 for I_sumMac as a function of δ (Extended Data Fig. S3i, Supplementary Table 13). For determining trend sizes depicted in Fig. 3n, the following calculation was used: denote the per image frequencies of a cell type as V, and the corresponding per image temporal stamps (either GA or mean image δ) as X. Trend size is then calculated as the difference between the first and last time point in units of the mean: 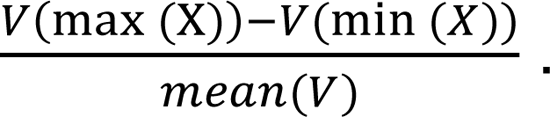

### NanoString GeoMx™ Digital Spatial Profiler (DSP)

The experiment was performed by NanoString Technologies, according to company manuals, details below.

#### Slide preparation

Serial sections of the TMAs were cut into 5 µm FFPE sections and were mounted on SuperFrost Plus slides (Fisher Scientific, 12-550-15), air dried and baked overnight in 60°C. Slides were then processed as specified by the NanoString GeoMx DSP Slide Preparation User Manual (NanoString Technologies, MAN-100 7). Briefly, slides were dewaxed, underwent antigen retrieval and treated with Proteinase K (Ambion, 2546) at 1 ug/mL concentration. Slides were then post-fixed. For RNA probes hybridization, slides were placed in a slide Rack with Kimwipes damped with 2xSSC lining the bottom. Each slide was treated with 200uL of NanoString Technologies whole transcriptome RNA probe Mix at a concentration of 4nM per probe in 1x Buffer R (NanoString Technologies). A Hybridslip (Grace Biolabs, 714022) was applied over each slide. Slides were incubated at 37°C overnight. After hybridization, slides were dipped in a 2x SSC + 0.1% Tween-20 (Teknova, T0710) to remove the coverslips. They were then washed twice in 2x SSC/50% formamide (ThermoFisher AM9342) at 37°C for 25 minutes followed by two washes in 2x SSC for 5 minutes each at room temperature. Slides were blocked in Buffer W (NanoString Technologies) at room temperature for 30 minutes, followed by application of 200uL morphology marker mix for 1 hour. Morphology markers details appear in Supplementary Table 14.

#### Sample collection

Sample collection was performed as indicated in the GeoMx DSP instrument user manual (MAN-10088-03). Slides were loaded to the GeoMx DSP instrument and scanned. For each tissue sample, we selected Regions of Interest (ROIs) corresponding to one of the following categories: artery (13), decidua (13), interstitial EVT (5), intravascular EVT (3); 34 ROIs in total were selected (Supplementary Table 15). Morphology markers for SMA and Vimentin were used in conjunction with a serial H&E section provide tissue context and to locate arteries and decidua on the platform. Artery, decidua, and intravascular EVT ROIs were selected using the geometric selection tool, and interstitial EVTs were selected using an HLA-G^+^ mask. Intravascular EVTs were identified as HLA-G^+^ cells located within arteries. Each ROI was collected into a single well in a 96 well plate.

#### GeoMx DSP NGS Library Preparation and Sequencing

Each GeoMx sample/well was uniquely indexed using Illumina’s i5 ⨯ i7 dual-indexing system. 4 uL of a GeoMx DSP sample was used in a PCR reaction with 1 uM of i5 primer, 1 uM i7 primer, and 1X NSTG PCR Master Mix. For PCR amplification reaction, each 96 well plate was placed in a thermocycler programmed with the following protocol: 37°C for 30 min, 50°C for 10 min, 95°C for 3 min, 18 cycles of 95°C for 15 sec, 65°C for 60 sec, 68°C for 30 sec, and final extension of 68°C for 5 min. PCR reactions were purified with two rounds of AMPure XP beads (Beckman Coulter) at 1.2x bead-to-sample ratio. Libraries were paired-end sequenced (2×75) on a NextSeq550 up to 400M total aligned reads.

#### Normalization and scaling of GeoMx counts data

Raw counts from each gene in each sample were extracted from the NanoString GeoMx NGS processing pipeline (Supplementary Table 15). QC was done according to the NanoString data analysis manual (MAN-10154-01) with default parameters as indicated in the manual. For each EVT sample, the counts were normalized using one of the manufacturer’s recommended approaches for normalizing GeoMx data: dividing all genes in each sample by the 75th percentile of expression in that sample, followed by multiplication by an identical scaling factor for all samples: the geometric mean of all 75th percentiles. This approach eliminates differences in counts between samples due to ROI specific properties such as size and RNA binding efficiency. The background due to non-specific binding per sample was approximated with the geometric mean of the 100 negative control probes included in the probe mix-as recommended by NanoString Technologies. The above-described normalization eliminated the correlation between background and ROI size for EVT samples. For artery and decidua samples, normalization is complicated by the fact that ROI size is tightly correlated with SAR stage and therefore biologically meaningful trends in the data. This led to the correlation between ROI size and background not being entirely eliminated by normalization. We therefore employed a background subtraction correction technique prior to normalization as recommended in the NanoString Technologies manual for such cases. The correction was performed by subtracting the geometric mean of negative probes from gene counts on a per sample basis and proceeding with normalization as previously described.

### Gene expression in artery as function of GA and SAR

Broadly, for each gene, we performed polynomial regressions of gene expression with δ and GA as the independent variables and used regression p-values to determine which genes were trending and the ratio of regression R^2^ values to classify the trends as detailed below.

The Nanostring technologies RNA probes panel contains probes for 18696 transcripts. For this analysis on artery samples, only genes with background subtracted, normalized counts ≥10 in at least two artery were considered. This resulted in 14471 expressed genes. Each artery sample was assigned a remodeling score δ, based on the δ of the sampled artery in the MIBI data. If several arteries were sampled-the assigned δ was the average δs of the sampled arteries. Endothelial loss and SMA loss per sample were calculated similarly, based on the corresponding MIBI values (Supplementary Table 15). The following steps were then performed on artery samples.

For all expressed genes, gene expression as a function of GA was defined as the background subtracted and normalized counts for that gene, as function of the GAs associated with the samples. Similarly, expression as a function of SAR for a gene was defined as the per-sample background subtracted and normalized counts of that gene, as function of the δ values per sample. A second-degree polynomial regression model was then fitted to the two above-described functions. The reason for using a second-degree polynomial instead of linear regression was to allow the regression models to capture non monotonic trends in gene expression. All regression models were implemented using the MATLAB function fitnlm. Expression fold change was defined as the ratio between maximum and minimum of expression values. The Center Of Mass, or COM of a gene’s expression trajectory as a function of t (t being either GA or δ) was defined as the weighted mean of t values, where the weights are the expression values at the respective t.

Genes with a p-value≤ 0.05 and fold change ≥ 2 for either GA or δ regression were classified as trending genes. The ratio between R^2^ in the two regression models was used to classify trending genes as GA-driven, SAR-driven or synchronized. Trending genes with log_2_(R^2^_δ_ /R^2^_GA_) ≥ 1 and R^2^_δ_ ≥ 0.05 were classified as SAR-driven, while genes with log_2_(R^2^_δ_ /R^2^_GA_) ≤ 1 and R^2^_GA_ ≥ 0.05 were classified as GA-driven. Other trending genes were classified as synchronized (Supplementary Table 16).

For visualization only, two fitted expression trajectories - one as a function of GA and another as function of δ were calculated per gene. These fitted expression trajectories were calculated as the values of the fitted 2nd degree polynomial model at 5 evenly spaced values of GA and δ respectively. To compare fitted expression trajectories between genes, they were normalized by z-scoring their value per gene (Fig. 3o, Extended Data Fig. S3k).

### Gene expression in decidua as function of GA, SAR and tissue composition

For this analysis on decidua samples, only genes with background subtracted, normalized counts ≥10 in at least two decidua samples were considered. This resulted in 13104 expressed genes.

In order to estimate the contribution of cell type frequencies in decidua samples to the observed changes in gene expression, cell-type abundancies were estimated in decidua ROIs collected using online version of the deconvolution algorithm CIBERSORTx^102, 103^. A custom signature matrix was created using the droplet-based 10x single-cell transcriptome profiles from 30 maternal-fetal tissue samples from Vento-Tormo et al.^4^ (dataset: https://www.ebi.ac.uk/biostudies/arrayexpress/studies/E-MTAB-6701#). Only cell annotations originating from decidua and placenta were used in creating the signature matrix. This signature matrix was used to impute cell fractions on the Nanostring decidua ROIs (Supplementary Table 16).

We then compared the cell type frequencies imputed by CIBERSORTx to the cell type frequencies we measured by MIBI in the decidua masks of the matched MIBI images. We limited this analysis to cell types that are abundant enough in decidua based on our MIBI measurements (frequency ≥1%). In order to match cell type annotations between the two different data sets, we grouped the cell types from the Vento-Tormo et al. dataset and matched them with MIBI as specified in Supplementary Table 16.

We then leveraged the deconvolved cell frequencies to predict which genes are trending with gestational age in decidual samples and which trends are better described by tissue composition or SAR. To do that, for each gene, we performed the following analysis:

1. Perform separate polynomial regressions (same as analysis described for artery samples above) of gene expression as a function of gestational age, SAR and each of the six cell type frequencies described in Supplementary Table S16
2. Considering only regression models with p-values ≤ 0.05 and fold change ≥ 2, rank models by their R^2^
3. Classify the gene as trending with gestational age, SAR or with one of the cell frequencies based on the model with maximal R^2^
4. If there were no regression models with p-values ≤ 0.05 and fold change ≥ 2 for the gene, classify it as non trending

### Coordinated gene expression by pathways in artery and decidua

As we set out to find gene pathways with temporally coordinated expression trends amongst two lists of genes of interest: genes trending with GA in decidua, genes trending with δ in arteries. For this analysis, in decidua samples we considered all genes for which GA trend was significantly better than SAR trend regardless of tissue composition trends. To find these coordinated pathways, we must first define the pathways and then define temporal coordination.

To define pathways, we used the R package msigdbr to get the lists of genes per pathway for the Gene Ontology by Biological Process (GO-BP) database (7481 pathways). We then cross referenced the list of genes for each pathway with the genes of interest and discarded pathways with an intersection of less than 10 genes. For the remaining pathways, we examined whether the pathway genes that appear in the gene set of interest exhibit temporal coordination in their expression.

A group of temporally coordinated genes was defined as a group of genes for which the COMs were significantly closer to each other in time than one would expect at random (time is either GA or δ, see previous section for definition of COM). Using the spread of COMs as a measure for temporal coordination allowed us to leverage the raw data rather than fitted gene expression trajectories, while still maintaining robustness to noise.

To calculate the extent of coordination between a group of N genes, we first calculated their median COM, denoted COM_med_. Then, their COM dispersal was defined as the median of the absolute deviations from COM_med_ for the N genes-denoted CD. In order to determine whether the CD for the gene group, CD_group_, is significantly smaller than expected at random, we calculated the randomly expected CD denoted CD_rand_. This was done by selecting N random genes without replacement and calculating their CD, 10^5^ times to estimate the null distribution. The random CD_rand_ was then calculated as the median over the CDs for randomized gene sets. The coordination score for our N genes group was then defined as log2(CD_rand_/CD_group_). The p-value for the coordination score was defined as the number of times a randomized CD was smaller than CD_group_, divided by the number of randomizations (10^5^). (1/number of randomizations) was then added to all p-values to account for the finite number of randomizations. q-values were calculated using the Benjamini and Hochberg method on p-values, implemented by the MATLAB function mafdr.

The CD, coordination scores, p-values and q-values were calculated as described above for the two lists of genes of interest, over all 7481 pathways. Pathways with coordination score≥1.5 and p-value≤0.05 were considered to be coordinated (Supplementary Table 5).

### Ridge regression for predicting GA from immune composition

Ridge regression was implemented using the sklearn Python package (sklearn.linear_model.Ridge, RidgeCV). Per-image immune frequencies were rescaled to the range 0-1 prior to model fitting, using the sklearn scaling function. Images with fewer than 10 immune cells were excluded (n=8). A randomly derived test-train split of 30/70 was used and GA distribution was verified to be equally represented in the test and train sets (Extended Data Fig. S4a). Ridge regression adds a regularization penalty to the loss function in order to prevent over or under representation of correlated variables, such as immune cell populations. The penalty used for the test set (0.81) was selected using Leave-One-Out Cross-Validation on the training set.

### Cell-cell and cell-artery spatial enrichment analysis

To identify preferential colocalization of maternal immune cells in decidua, we measured the spatial proximity enrichment for all cell type pairs, which evaluates the spatial organization of cell types relative to each other, as described previously^47–49^. Cells located in non-decidual feature masks (artery, gland, vessel, or cell column masks) were not included in this analysis. The distances in pixels (px) between all pairs of cells were calculated in each image. The resulting per image distance matrices were binarized with a distance threshold (100px or 39 μm in our case), pairs of cells closer than 100px from each other were considered a close interaction. To evaluate the number of close interactions between two cell types, this proximity matrix can be subset column wise by cell type A, and subset row wise by cell type B. The sum of the resulting submatrix quantifies the number of close interactions between the cells of types A and B. To evaluate the significance of the number of close interactions, given the total number of cells in the image, tissue architecture and composition across the cohort, and total number of cells of types A and B in the image-a bootstrapping approach was used. For each of 100 bootstrapping iterations, the location of cells of type A was randomized across all cell locations (of any type) in the image, while their total number was preserved. The number of close interactions with cells of type B is calculated for each randomized iteration. Repetitions of this process approach a null distribution for the number of close interactions between cells A and B. The enrichment score for cells A around cells B in the image is then calculated as the Z-score of the measured number of close interactions between A and B, when Z-scored together with the random bootstraps. This analysis was extended to incorporate enrichment of cell types around spiral arteries. For each cell, the distance to the nearest spiral artery was considered. An additional column was added to the proximity matrix described above which thresholded distances between cells and arteries with the same 100px threshold. The above-described bootstrapping approach provides a null distribution for artery proximity as well. Tools for this analysis were written in Python, with the bootstrapping accelerated via Cython. An intuitive, easy to use Jupyter Notebook interface was created to allow for easy implementation of this algorithm. For per image spatial enrichment scores, see Data Availability Statement. The code for this analysis is available on https://github.com/angelolab/ark-analysis.

### Cell-cell/cell-artery enrichment temporal trends: trending with GA/ SAR or constant

For examining cell-cell/ cell-artery enrichment within the decidua as function of GA and SAR (Fig. 4k, Extended Data Fig. S3j), per image enrichment score matrices E were calculated as described in the previous section, in which E_i,j_ is the enrichment score of cell type i around cell type j in the image. Enrichment as a function of GA was defined as the per image enrichment, as function of the GAs associated with the images. Similarly, enrichment as a function of SAR was defined as the per image enrichment, as function of the mean δ values per image. We fitted a linear regression model to the two above-described functions. All linear regression models were implemented using the MATLAB function fitlm. R^2^ and p-values for all δ and GA based regressions can be found in Supplementary Table 6. The ratio between R^2^ in the two regression models was used to classify trends as GA-driven, SAR-driven or synchronized like in Fig. 3n. Fig. 4k only shows points for which regression R^2^ ≥ 0.05, p-val ≤ 0.05, maximal absolute value of linear fit ≥2. Trends including muscle, fibroblast, myofibroblast, glandular, other and endothelial cells were not considered in this analysis. For determining trend sizes, the following calculation was used: denote the linear fit to per image enrichment scores as V, and the corresponding per image temporal stamps (either GA or mean image δ) as X. Trend size is then calculated as (V(max(X))-V(min(X))).

To determine whether two cell types were significantly enriched around each other throughout the cohort, we averaged their pairwise enrichment over all images. The pair was considered enriched if the absolute value of mean enrichment ≥2 (Supplementary Table 6).

In Fig. 4k, the following cell-cell enrichments were not plotted, for clarity:

1. Enrichment of a cell type around itself. i.e Tregs around Tregs
2. Enrichments including the following cell types: muscle, fibroblast, myofibroblast, glandular, other and endothelial
3. Enrichment trends that are SAR driven

### Functional markers positivity rate per cell type as function of GA and SAR

For examining cell type specific temporal trends in the expression of functional markers (Fig. 5a), 48 combinations of cell type-functional marker were selected. The selected combinations were those for which the positivity frequency Z-score exceeded 0.5 (Fig. 2a, right panel). For each of these combinations, the frequency of cells positive for the functional marker was calculated as the number of cells positive for the marker (see “Definition of thresholds for functional marker positivity”), out of the total number of cells of the same cell type in the image. All cells except those located within the cell column mask were included to focus the analysis on functional marker trends of maternal cells and EVTs that had infiltrated the decidua. For glandular cells, the location was further restricted to the glands mask. The frequency of cells positive for a functional marker as a function of GA, for a cell type, was defined as the per image positivity proportion values as function of the GAs associated with the images. Similarly, marker positivity frequency as a function of SAR for a cell type was defined as the per image proportions of that cell type positive for the marker, as function of the mean δ values per image. For the volcano plot in Fig. 5a, we fitted a linear regression model to the two above-described functions. All linear regression models were implemented using the MATLAB function fitlm and the volcano plot only shows points for which regression R^2^ ≥ 0.05. R^2^ and p-values for all δ and GA based regressions can be found in Supplementary Table 17. For determining trend sizes depicted in Fig. 5a, the following calculation was used: denote the linear fit to the per-image marker positivity proportion of a cell type as V, and the corresponding per image temporal stamps (either GA or mean image δ) as X. Trend size is then calculated as the difference between the first and last time point in units of the mean: 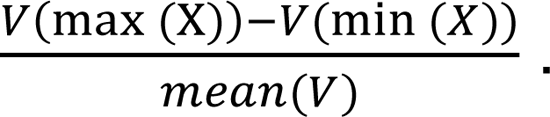

### Cellular microenvironments

For each cell in the dataset, we defined a “neighborhood” consisting of its 25 closest neighboring cells, as measured by Euclidean distance between X/Y centroids, excluding cells that were not in the decidua (i.e., cells that overlapped with any artery, gland, anchoring villous, or vessel feature masks). We clustered these cellular neighborhoods based on their composition of the 26 cell populations as identified previously using FlowSOM. For clustering, we used the scikit-learn implementation of k-Means algorithm with k=20 to identify neighborhoods characterized by the presence of rare cell populations. Selected clusters were merged based on similarity when hierarchically clustered, a threshold of 0.5 when comparing Euclidean distances between k-Means cluster centroids, and manual inspection of the cluster assignment when overlaid on the images. Based on these approaches we defined 12 distinct decidual cellular microenvironments, 10 of which are shown in Fig. 5f (microenvironments characterized by predominantly stromal cell populations, fibroblasts (3 in Supplementary Table 18) and myofibroblasts (6 in Supplementary Table 18), were not shown in the heatmap).

### Definition of anatomical EVT location and associated arteries

Cell column EVTs were defined as EVTs located within cell column masks, intravascular EVTs were located within artery masks, and interstitial EVTs were located in the decidua. Perivascular EVTs were defined as interstitial EVTs located within 50 pixels of the edge of an artery, as defined by radial expansion of the artery masks (Fig. 6b). Arteries were said to have perivascular or intravascular EVT (Fig. 6f, Extended Data Fig. S6c, d) if the number of EVT in the appropriate artery compartment was ≥5. For Extended Data Fig. S6f, only images that contained all four EVT types were considered and cell to artery distance was measured from the cell centroid as detected by segmentation to the border of the artery mask. For Extended Data Fig. S6e, one image was excluded (16_31762_20_8) due to abnormal tissue morphology.

### SMA and endothelium loss scores

The loss scores presented in Extended Data Fig. S6c, d were based on digitized morphological features. For SMA, the average feature was used and for endothelium, the radial coverage of CD31 (see “Automated digitization of artery morphological features”). The values for each of the two features were then divided by their maximum across arteries and subtracted from 1 to obtain a loss score. The resulting values were then linearly rescaled to the range 0-1 using the MATLAB function rescale.

### LDA of EVTs by compartment

For Extended Data Fig. S6g, a method similar to our calculation of the continuous SAR remodeling score δ was used for compartment-wise analysis of EVT types. The input table consisted of marker expression values per EVT. Lineage and functional markers expressed by EVTs were included: CD56, CD57, HLA-G, CK7, PD-L1 and Ki67 (Fig. 2a). EVTs were labeled by spatial compartment: cell column, interstitial, perivascular or intravascular (see “Definition of anatomical EVT location”). Marker expression values were standardized (mean subtracted and divided by the standard deviation) and cell column, interstitial, and intravascular location labels per EVT were used for training the LDA model. Perivascular EVTs were withheld as a test set. Due to the small number or features (markers) a one-dimensional LDA was calculated yielding a single coordinate LD1. LD1 was the optimal linear combination of a subset of markers, to maximize the separation by compartment between EVTs (Supplementary Table 19). LD1 values were subsequently calculated for the withheld test set of perivascular EVTs (Supplementary Table 20). To calculate the difference in similarity to intravascular EVT between interstitial and perivascular, the following calculation was used: intravascular-perivascular similarity was defined as *sim_intra-peri_ = mean(Ld1_intravascular_) - mean(Ld1_perivascular_)*. Similarly, intravascular-interstitial similarity was defined as *sim_intra-inter_ = mean(Ld1_intravascular_) - mean(Ld1_interstitial_)*. The difference in these similarities was then calculated as: 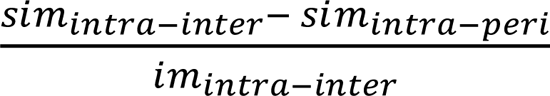 in %

### Origin of CD56+ EVTs in the intravascular compartment

The frequency of CD56 positivity for EVT increases with SAR both in the perivascular and intravascular compartment (Extended Data Fig. S6h, i). However, the increase in the intravascular compartment is steeper (Extended Data Fig. S6j). One potential explanation for the steep increase in EVT1c prevalence between the perivascular and intravascular compartment is that arterial intravasation of perivascular EVTs is accompanied by upregulation of CD56, such that EVT1a-b subsets would effectively become the EVT1c subset. We therefore hypothesized that such a process would involve an intermediate state between the two in which the EVT1a-b subsets moderately express of CD56 en-route to the high expression observed in the EVT1c subset. To test this hypothesis, we compared the average CD56 intensity of perivascular and intravascular EVT1a-b EVTs on a per-artery basis (for arteries that initiated remodeling: δ≥2). This analysis detected a statistically significant increase in CD56 expression between the perivascular and intravascular compartment by EVT1a-b subsets (sided Wilcoxon signed rank test p-value = 5e^-3^, Extended Data Fig. S6k). An alternative explanation for the disproportionate enrichment of EVT1c within vessels is that they are more proliferative. However, only 0.5% of intravascular EVT1c were Ki67+ compared to 9.6% and 1.8% of intravascular EVT1a and EVT1b cells, respectively (Extended Data Fig. S6l). This implies that the source of the multiplying CD56+ intravascular EVT1c are the perivascular EVT1a,b.

### Differentially expressed genes in EVT

Differentially expressed genes between intravascular and interstitial EVTs were identified using the Bioconductor package limma^104^ (Linear Models for Microarray Data) upon consulting with the NanoString statistics team. Using limma’s default parameters on 75 percentile normalized counts, 131 upregulated genes and 143 downregulated genes were found (FDR cutoff = 0.1, log Fold Change cutoff = 2). Genes with logFC ≥ 2.3 or ≤ 2.3 and adjusted p-value ≤ 0.05 are shown in the heatmap in Fig. 6h, the complete list of differentially expressed genes is shown in Extended Data Fig. S6m and Supplementary Table 21. For IHC validation of differential expression for selected genes, see Extended Data Fig. 6n.

### Niche-Net analysis

We used the NicheNet R package to predict ligand-receptor interactions between intravascular EVT and arteries. The analysis was performed by following the vignette: https://github.com/saeyslab/nichenetr/blob/master/vignettes/ligand_activity_geneset.md.

NicheNet requires three input gene lists, to predict ligands in sender cells which are likely to interact with receptors in receiver cells and impact the expression of genes of interest: genes of interest, genes expressed in sender cells and genes expressed in receiver cells.

For our analysis, we wanted to check which ligands expressed in the intravascular EVT are likely to be causing temporal gene expression trends with remodeling in arteries. To do so, we defined the genes of interest as all genes trending with remodeling in arteries (Fig. 3o). The genes expressed in receiver cells were defined as all genes expressed in arteries (see previous sections for definition of “expressed”) and genes expressed in sender cells were defined as genes differentially expressed between interstitial and intravascular EVT, and higher in intravascular (Supplementary Table 22). NicheNet analysis was performed as described in the vignette to prioritize ligands, infer corresponding receptors and downstream targets (Extended Data Fig. S6o). The inferred targets were manually classified according to their known function using the “Gene Cards” database (https://www.genecards.org) and literature survey. A list of references for all classifications can be found in Supplementary Table 7.

### Statistical analyses

Throughout the paper, the Kruskal-Wallis test was implemented using the MATLAB function KruskalWallis. All linear regression models were implemented using the MATLAB function fitlm unless stated otherwise. The sided Wilcoxon signed rank test for paired analysis in Extended Data Fig. S6 was implemented using the MATLAB function signrank. MATLAB version used throughout the paper for statistical analysis is MATLAB 2020b.

### Data availability

Patient block images with annotations, imaging data, segmentation masks, extracted features, CPMs, per cell-cell and cell-artery spatial image enrichment scores, a table that enumerates all single cells in this study and provides their location, morphological characteristics such as size and shape, marker expression, FlowSOM cluster assignment and cell type assignment are available for review at: https://datadryad.org/stash/share/kP1VKIm5ZalIF_CXZnpy0VAfTmkP_ei9t58uK84pl6U And will be made publicly available upon publication.

### Code availability

Software used for analysis in the paper: ImageJ, MAUI (https://github.com/angelolab/MAUI) for low level image processing, DeepCell (https://deepcell.readthedocs.io/en/master/index.html) for cell segmentation. Ark Analysis for cell-cell spatial enrichment (https://github.com/angelolab/ark-analysis). Custom code for this study is available for review at https://figshare.com/s/f78e3885d5afe241cd38 And will be made publicly available upon publication.

## Supporting information

All Supplementary Tables

## Acknowledgements

We thank M. Amouzgar, C. Liu, N. Vivanco, A. Moore, E. McCaffrey, D. Glass, T. Risom, J. P. Oliveria, K. O’Neill and C. Coutifaris for comments. M.A. is supported by 5U54CA20997105, 5DP5OD01982205, 1R01CA24063801A1, 5R01AG06827902, 5UH3CA24663303, 5R01CA22952904, 1U24CA22430901, 5R01AG05791504, and 5R01AG05628705 from NIH, W81XWH2110143 from DOD, and other funding from the Bill and Malinda Gates Foundation, Cancer Research Institute, the Parker Center for Cancer Immunotherapy, and the Breast Cancer Research Foundation. S.G is supported by the Bill and Melinda Gates Foundation OPP1113682. I.A. is an awardee of the Weizmann Institute of Science – Israel National Postdoctoral Award Program for Advancing Women in Science. E.S is supported by National Science Scholarship, Agency for Science, Technology, and Research (A*STAR), Singapore.

## Author contributions

S.G. assembled the tissue cohort, performed and designed experiments, annotated images, analyzed and interpreted data and wrote the manuscript. I.A. analyzed and interpreted data, wrote the manuscript. E.S. performed and designed experiments, annotated images, analyzed and interpreted data, wrote the manuscript. G.R. advised on cohort design, assembled tissue cohort, annotated images. A.B, N.G., A.K, G.M., M.S., W.G. and D.V.V. wrote software for image analysis. M.B., H.P and V.W advised on experimental design and reagent validation. E.J. assembled cohort patient metadata. L.K. advised on computational analysis. Z.K. Prepared and validated reagents. S.K. Constructed the tissue microarray. S.W. annotated images. T.H. validated reagents and advised on experimental design. M.R. oversaw tissue microarray construction. M.A. conceived the study, advised on experimental design and data analysis, wrote the manuscript.

## Supplementary table legends

**Supplementary Table 1 - Patients table.** Patient meta-data such as age, ethnicity, body mass index, parity and relevant medical conditions such as HIV.

**Supplementary Table 2 - Information on the histological characteristics of the blocks retrieved, including the presence of cell column anchoring villi.** Per patient’s block, indication whether cell column anchoring villi were present (1) or absent (0), and the number of regions containing spiral arteries annotated as appropriate for TMA construction by the pathologist (Methods). In blocks containing ≥ 2 distinct, separate pieces of tissue, cell column villi were considered present if they were present on any piece containing pathologist annotations.

**Supplementary Table 3 - Artery properties and staging.** Arteries meta-data, including their measured digitized morphological features (see Methods), manual stage and remodeling score δ.

**Supplementary Table 4 – Cell-artery spatial enrichment.** Linear regression results of cell types enrichment around arteries as function of remodeling score δ. Each row represents a cell type. The columns present regression R^2^, p-value, maximal absolute value of enrichment and trend size (see Methods).

**Supplementary Table 5 – Synchronized expression of GO-BP pathways in arteries and decidua.** List of the temporally synchronized gene ontology pathways for artery (first tab, as function of SAR) and decidua (second tab, as function of GA). See Methods.

**Supplementary Table 6 – Cell-cell spatial enrichment.** Information about temporal trends in cell-cell enrichment (first tab) and constant in time cell-cell enrichment (second tab). For temporal trends, each row represents a combination of 2 cell types in the format: cell type a-cell type b. The information in the row is for the enrichment of cell type a around cell type b. The columns show values for the linear regression on per image basis of enrichment scores as function of GA and remodeling score δ: R^2^, the maximal obtained regression R^2^, p-value, maximal enrichment (absolute value) and trend size (see Methods).

**Supplementary Table 7 – NicheNet derived target genes and their function.** Comprehensive list of all NicheNet predicted target genes in artery (see Methods) with function classification and references for classification.

**Supplementary Table 8 – Genes differentially expressed in decidua samples from women diagnosed with preeclampsia.** List taken from PMID: 28618048.

**Supplementary Table 9 - Information on antibodies, metal reporters, staining concentrations, and parameters used for low-level processing of MIBI data.** Per marker antibody information, including: clone, vendor, vendor ID, channel and elemental reporter, and final staining titers used. The parameters used for marker-specific low-level processing of MIBI data (background removal, denoising, and aggregate removal steps as previously described) are also shown.

**Supplementary Table 10 – Number of segmented events per image.** Number of single cells per image, after segmentation, post processing and exclusion of small non-cellular objects. See Methods.

**Supplementary Table 11 - Positivity binary threshold for functional markers.** Binary expression thresholds per functional marker-used to determine whether cells are positive for that marker (See methods).

**Supplementary Table 12 - LDA coefficients for artery morphological features.** Per feature coefficients for the featurse that define the LDA space used for digitized artery staging (Fig. 3m). For the features selected by the algorithm, their ld1 and ld2 coefficients are listed. Additional columns show the Z scored absolute values of these coefficients.

**Supplementary Table 13 - Regression results for cell type proportions as a function of GA and δ.** Values plotted in Fig. 3n. Each row represents a cell proportion with those starting with R_ indication proportion out of all cells in the image, I_ - proportion out of immune cells in image, N_- proportion out of NK cells in image, M_ proportion out of macrophages in image, T_- proportion out of T cells in image, F_- proportion out of EVT in image, S_ proportion out of structural cells in image (Fig. 2b). The columns show values for the linear regression on per image proportions as function of GA and remodeling score δ: the log transformed ratio of R^2^, the maximal obtained regression R^2^ (maximal between GA and δ), the minimal obtained regression p-value and trend size (see Methods).

**Supplementary Table 14 – Nanostring morphology markers information.** Per marker antibody information including clone, channel and final staining titers used.

**Supplementary Table 15 – Nanostring ROI meta-data and raw counts.** Meta-data for ROIs (first tab), mapping Nanostring ROIs to their matching MIBI images of (second tab) and raw counts data per ROI (third tab).

**Supplementary Table 16 – Regression results for gene expression in artery and decidua as a function of GA and δ.** Per gene second degree polynomial regression R^2^, p-values and fold change as function of GA and δ for artery ROIs (first tab). Decidua ROIs deconvolution results using CIBERSORTx (second tab), match between deconvolved cell types and MIBI (Fig. 2a,b) cell types (third tab) and per gene second degree polynomial regression R^2^, p-values and fold change as function of GA, δ and cell frequencies for decidua ROIs (fourth tab), see Methods.

**Supplementary Table 17 - Regression results for functional markers expression as a function of GA and δ.** Values plotted in Fig. 5a. Each row represents a combination of a cell type and a functional marker. The columns show values for the linear regression on per image marker positivity rates for the marker-cell type as function of GA and remodeling score δ: the log transformed ratio of R^2^, the maximal obtained regression R^2^ (maximal between GA and δ), the minimal obtained regression p-value and trend size (see Methods).

**Supplementary Table 18 – Cell microenvironment lineage composition.** Fraction of cells from each lineage, for each of the 12 cell microenvironments. See Methods.

**Supplementary Table 19 - LD1 coefficients for markers expressed by EVT.** Coefficients per EVT expressed marker that define the LDA space used for measuring similarity between anatomical tissue compartments (see Methods, Extended Data Fig. 6g). For the markers selected by the algorithm, their ld1 coefficients are listed. An additional column shows the Z scored absolute values of these coefficients.

**Supplementary Table 20 - LD1 values per EVT.** ld1 values per single EVT with LDA input-standardized marker expression values (see Methods, Extended Data Fig. 6g). Additional metadata such as EVT type, anatomical location, the image the cell was taken from is also provided.

**Supplementary Table 21 – Differentially expressed genes between interstitial and intravascular EVTs.** See methods.

**Supplementary Table 22 – NicheNet input gene lists.** First tab-Genes Of Interest (GOI) – SAR trending genes in artery, second tab: genes expressed in receiver cells – all genes expressed in artery, third tab: genes in sender cells – DEG between intravascular and interstitial EVTs, higher in intravascular. See methods.

## Notes

### Competing Interest Statement

MA is a cofounder and stockholder in Ionpath Inc which is commercializing instrumentation for MIBI-TOF analysis.

### Summary of Updates

Addition and integration of a new coregistered spatial transcriptomics dataset to interrogate transcriptional changes with respect to spiral artery remodeling and EVT migration. Addition of 2 new spatial analyses that give insight into microenvironment dynamics in the decidua.

