## Extended Data Figure 1 for "Spatiotemporal coordination at the maternal-fetal interface promotes trophoblast invasion and vascular remodeling in the first half of human pregnancy"

### Extended Data Figures

#### Figure S1

Antibody Panel Single Channel Immune Controls

Tonsil

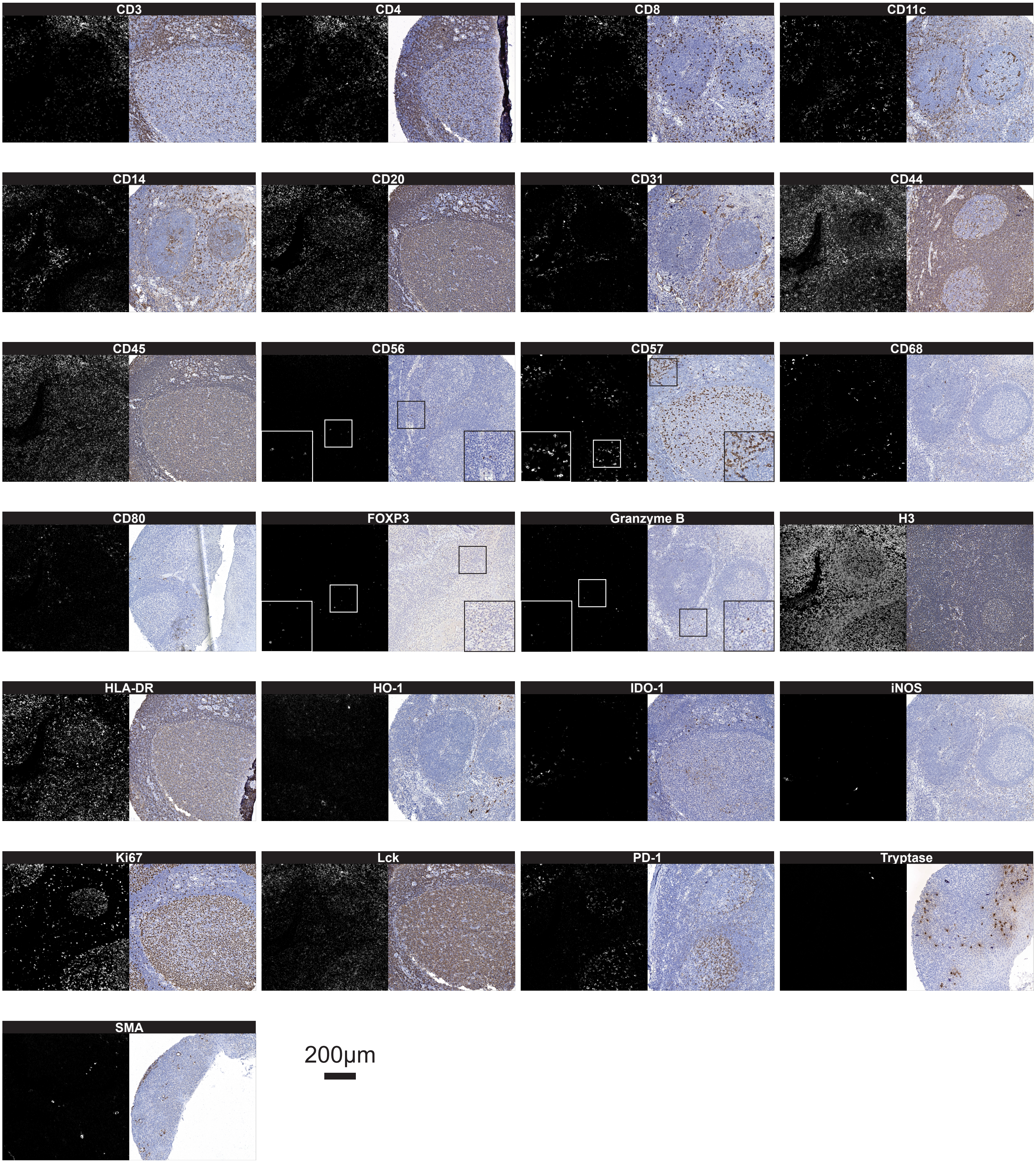

Placenta/decidua

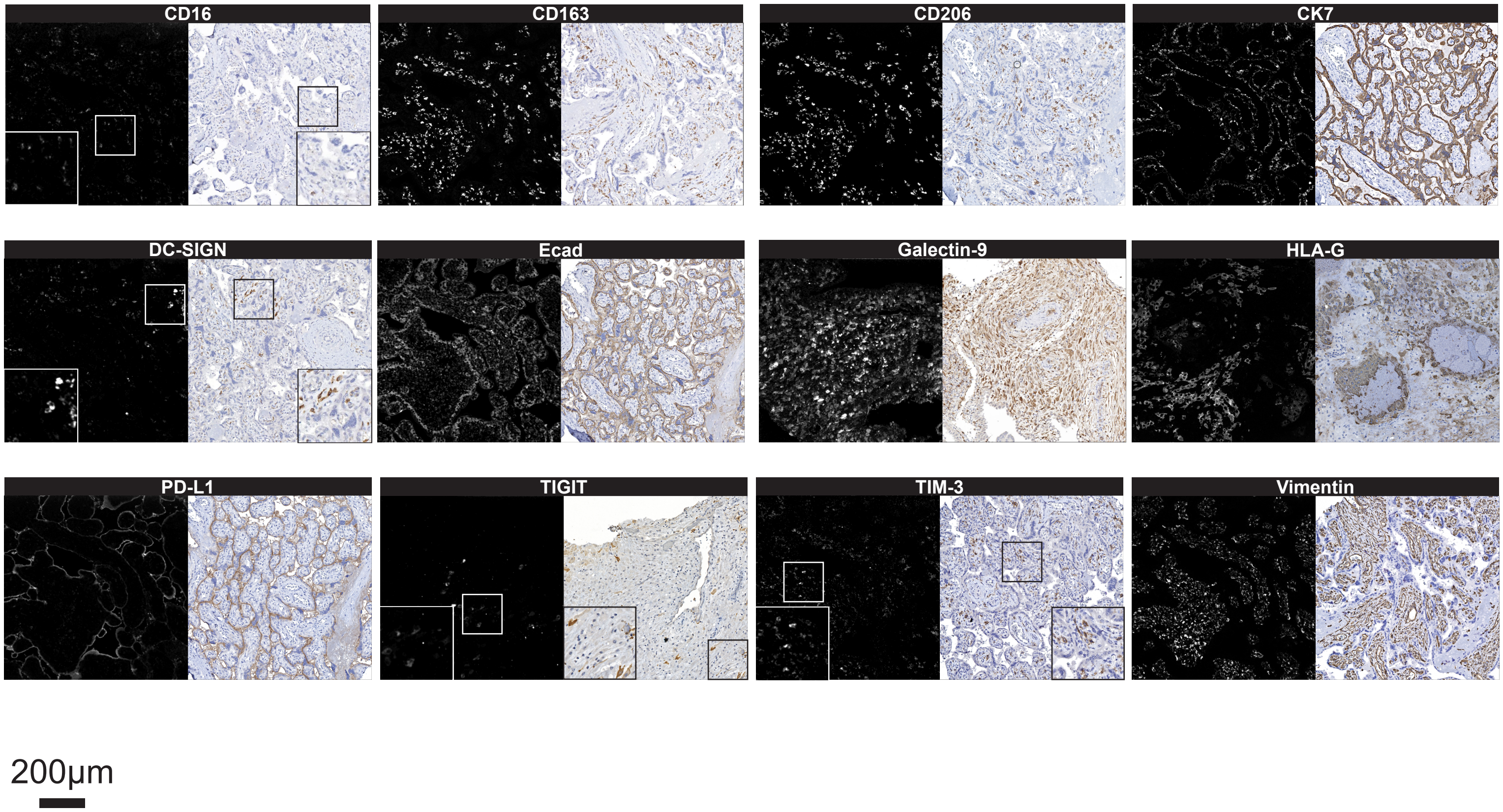

**Extended Data Figure S1 | MIBI-TOF antibody panel.** Representative images of MIBI conjugate staining for all markers along with serial single channel immunohistochemistry images, with immune control tissues (tonsil and placenta).
