## Extended Data Figures 2-6 for "Spatiotemporal coordination at the maternal-fetal interface promotes trophoblast invasion and vascular remodeling in the first half of human pregnancy"

#### Figures S2 – S6

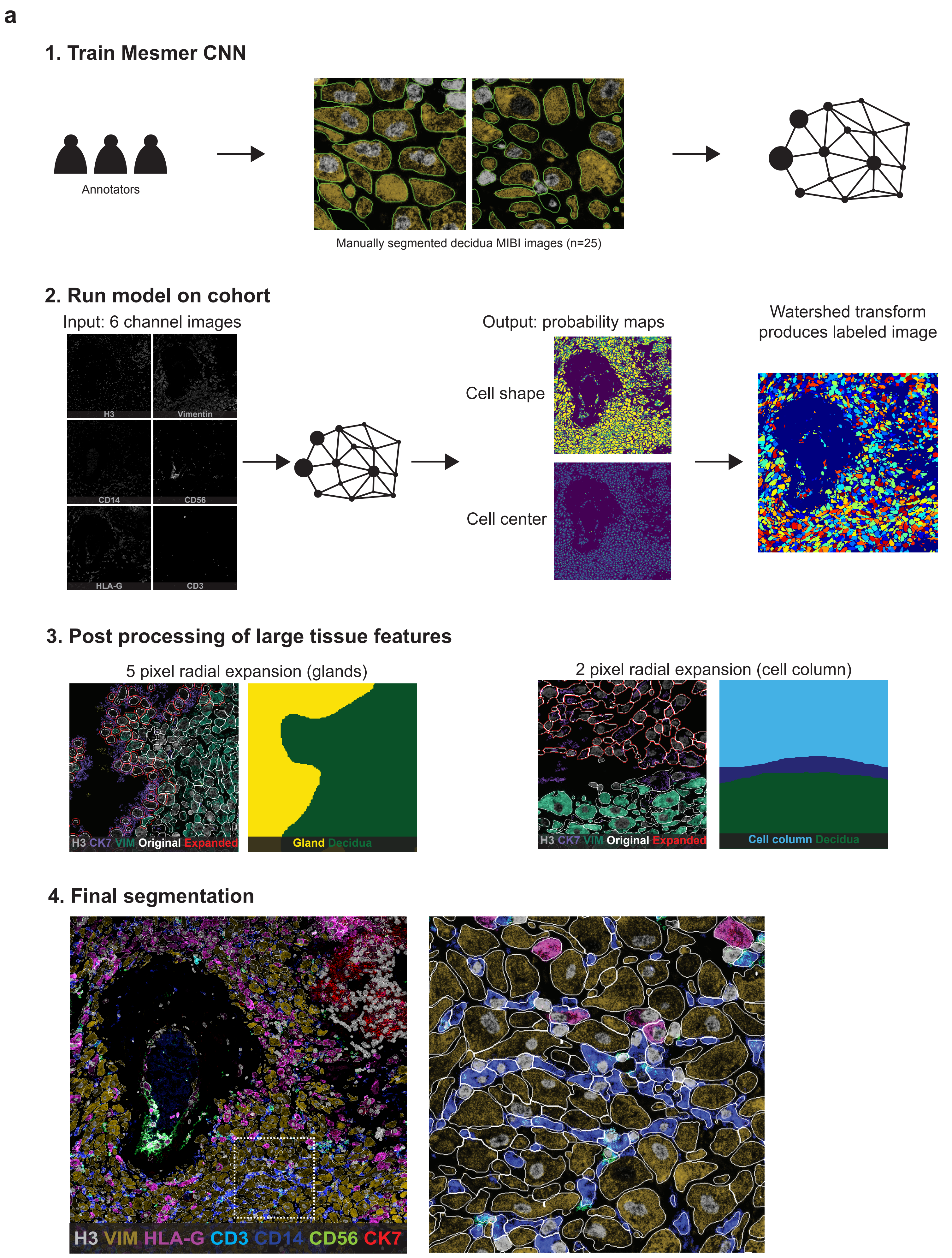

a
1. Train Mesmer CNN
Annotators
Manually segmented decidua MIBI images (n=25)
2. Run model on cohort
Input: 6 channel images
H3
Vimentin
CD14
CD56
HLA-G
CD3
Output: probability maps
Cell shape
Cell center
Watershed transform produces labeled image
3. Post processing of large tissue features
5 pixel radial expansion (glands)
H3 CK7 VIM Original Expanded
Gland Decidua
2 pixel radial expansion (cell column)
H3 CK7 VIM Original Expanded
Cell column Decidua
4. Final segmentation
H3 VIM HLA-G CD3 CD14 CD56 CK7

b

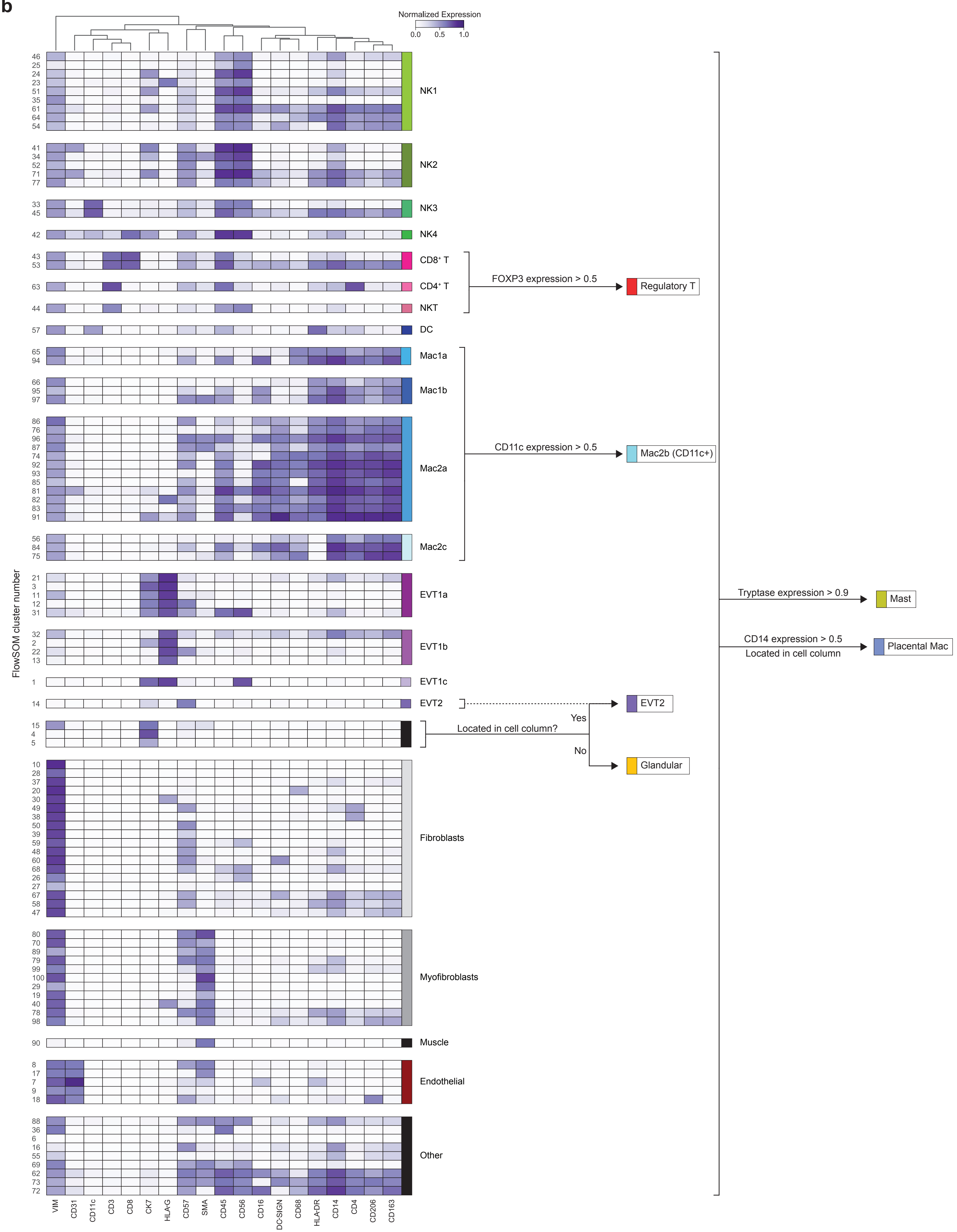
Normalized Expression
0.0
0.5
1.0
FlowSOM cluster number
NK1
NK2
NK3
NK4
CD8⁺ T
CD4⁺ T
NKT
DC
Mac1a
Mac1b
Mac2a
Mac2c
EVT1a
EVT1b
EVT1c
EVT2
Fibroblasts
Myofibroblasts
Muscle
Endothelial
Other
FOXP3 expression > 0.5
Regulatory T
CD11c expression > 0.5
Mac2b (CD11c+)
Tryptase expression > 0.9
Mast
CD14 expression > 0.5
Located in cell column
Placental Mac
EVT2
Yes
Located in cell column?
No
Glandular
VIM
CD31
CD11c
CD3
CD8
CK7
HLA-G
CD57
SMA
CD45
CD56
CD16
DC-SIGN
CD68
HLA-DR
CD14
CD4
CD206
CD163

**c**

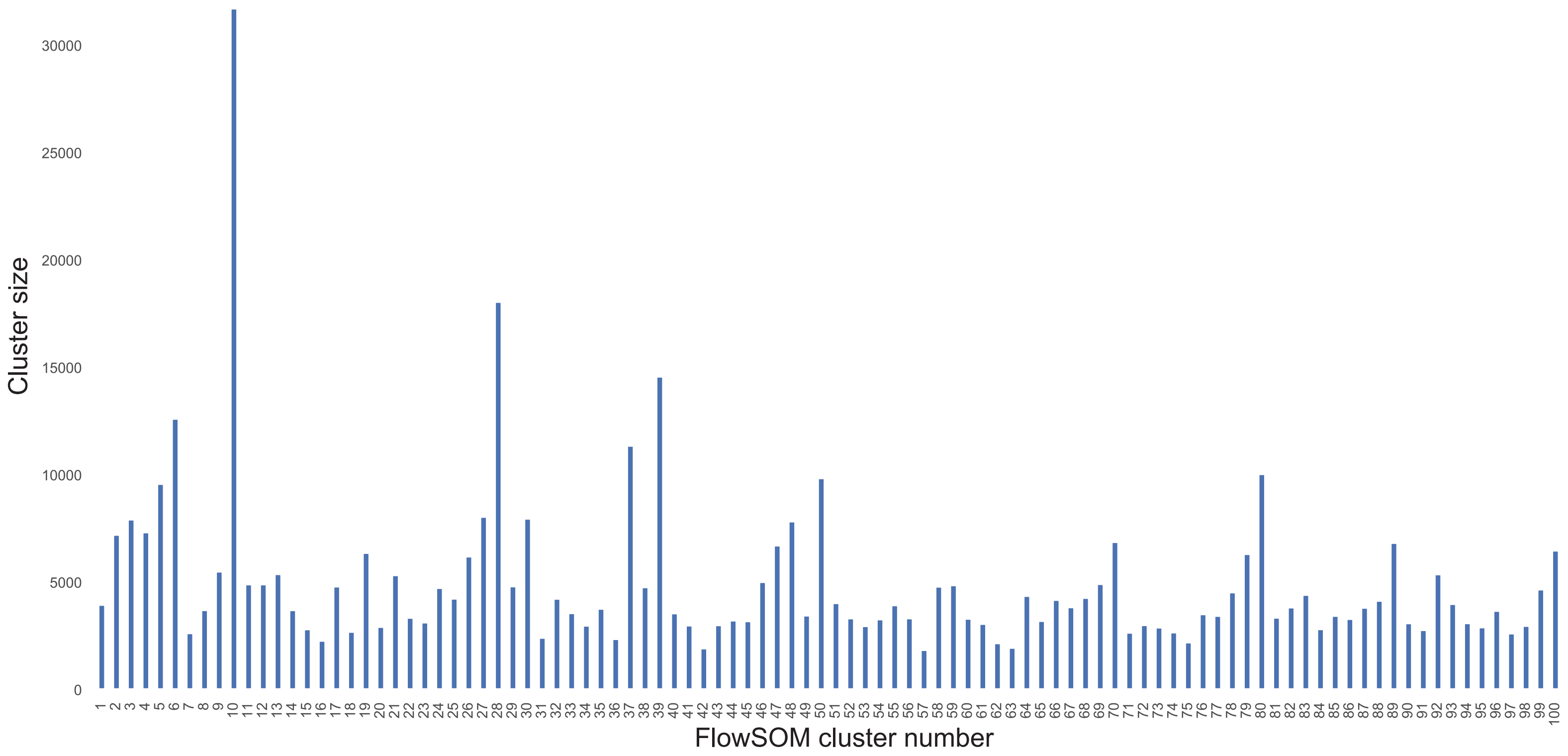
Cluster size
FlowSOM cluster number
30000
25000
20000
15000
10000
5000
0

**d**

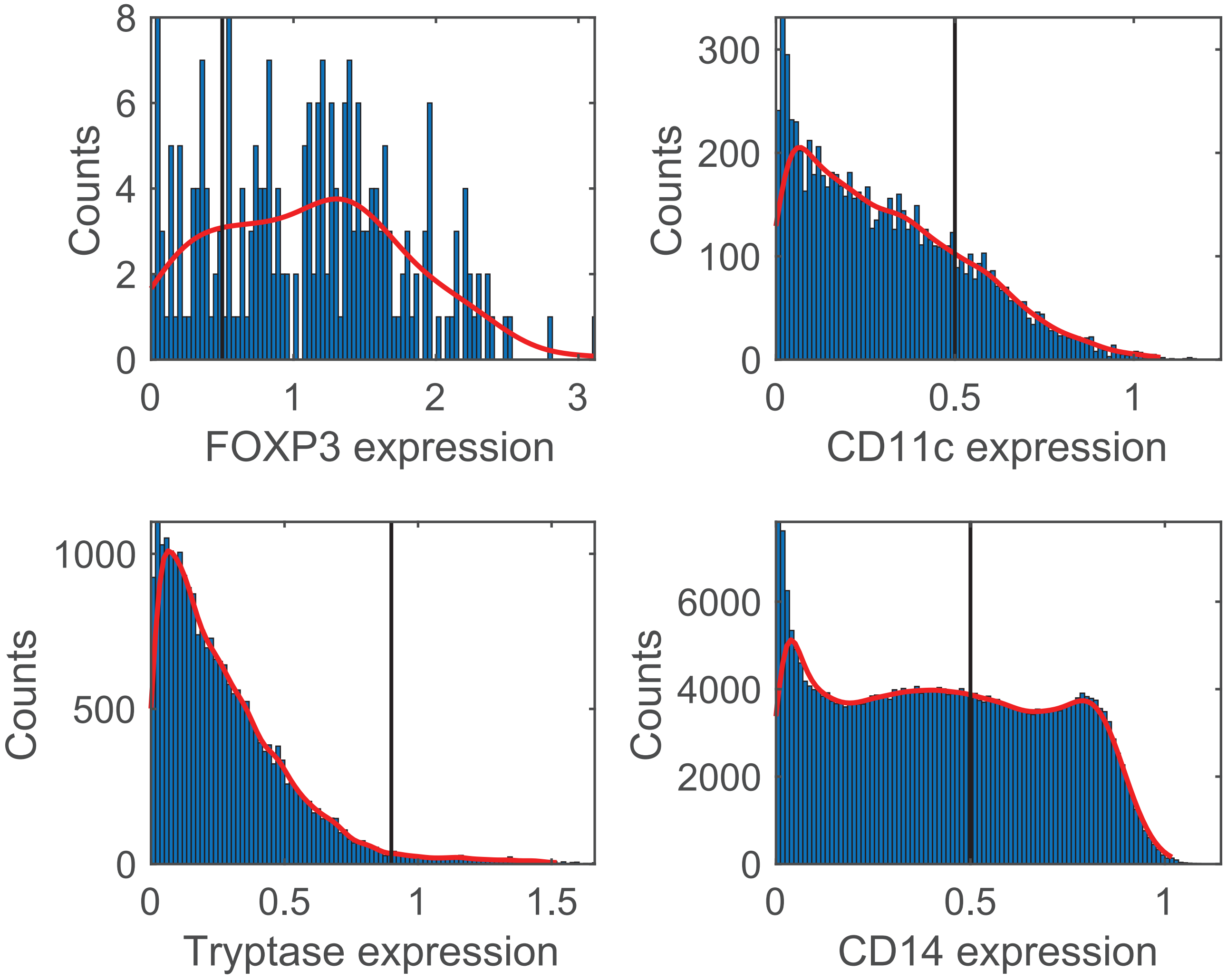
Counts
FOXP3 expression
Counts
CD11c expression
Counts
Tryptase expression
Counts
CD14 expression

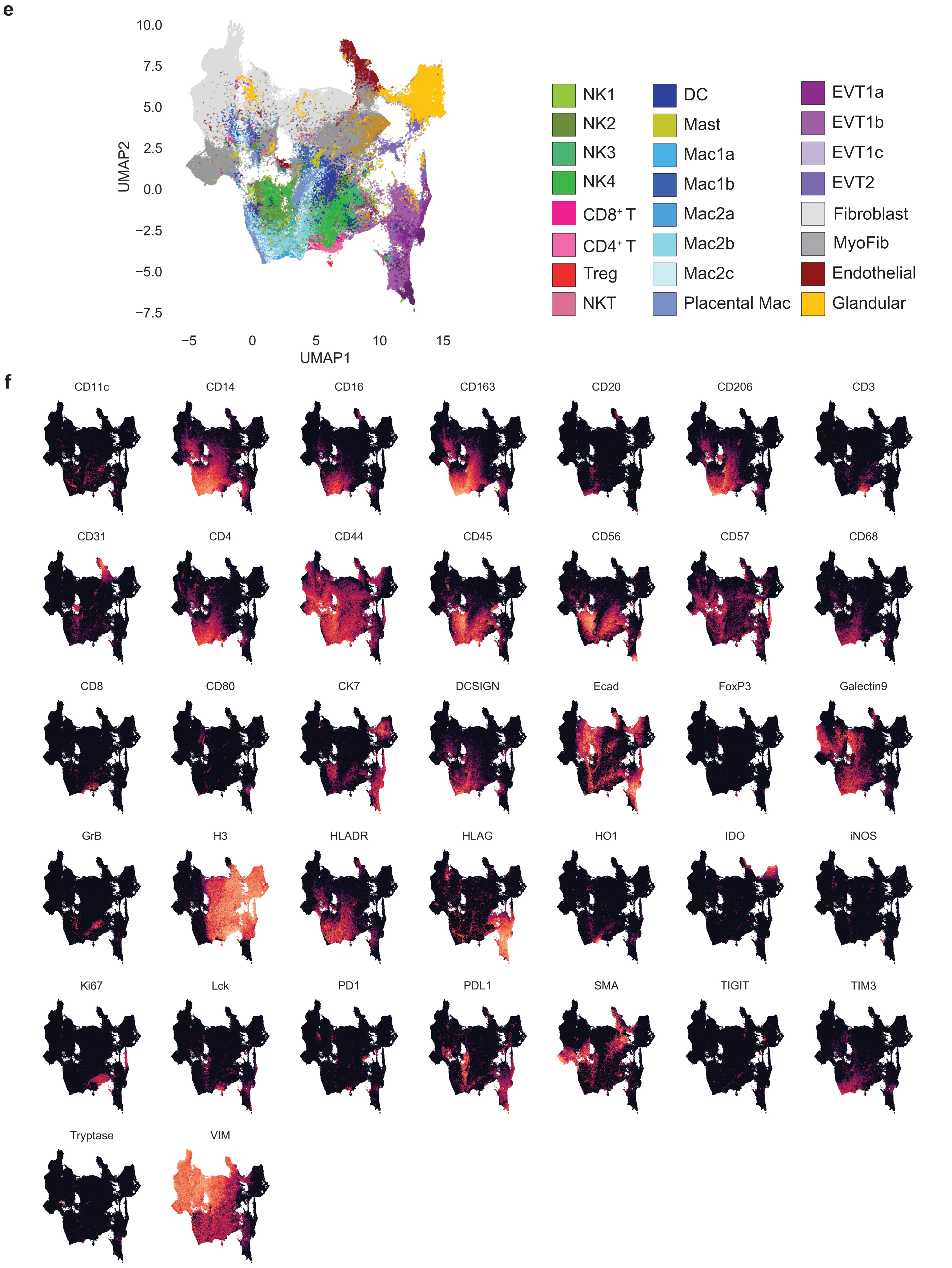
e
10.0
7.5
5.0
2.5
0.0
−2.5
−5.0
−7.5
UMAP2
−5
0
5
10
15
UMAP1
NK1
NK2
NK3
NK4
CD8⁺ T
CD4⁺ T
Treg
NKT
DC
Mast
Mac1a
Mac1b
Mac2a
Mac2b
Mac2c
Placental Mac
EVT1a
EVT1b
EVT1c
EVT2
Fibroblast
MyoFib
Endothelial
Glandular
f
CD11c
CD14
CD16
CD163
CD20
CD206
CD3
CD31
CD4
CD44
CD45
CD56
CD57
CD68
CD8
CD80
CK7
DCSIGN
Ecad
FoxP3
Galectin9
GrB
H3
HLADR
HLAG
HO1
IDO
iNOS
Ki67
Lck
PD1
PDL1
SMA
TIGIT
TIM3
Tryptase
VIM

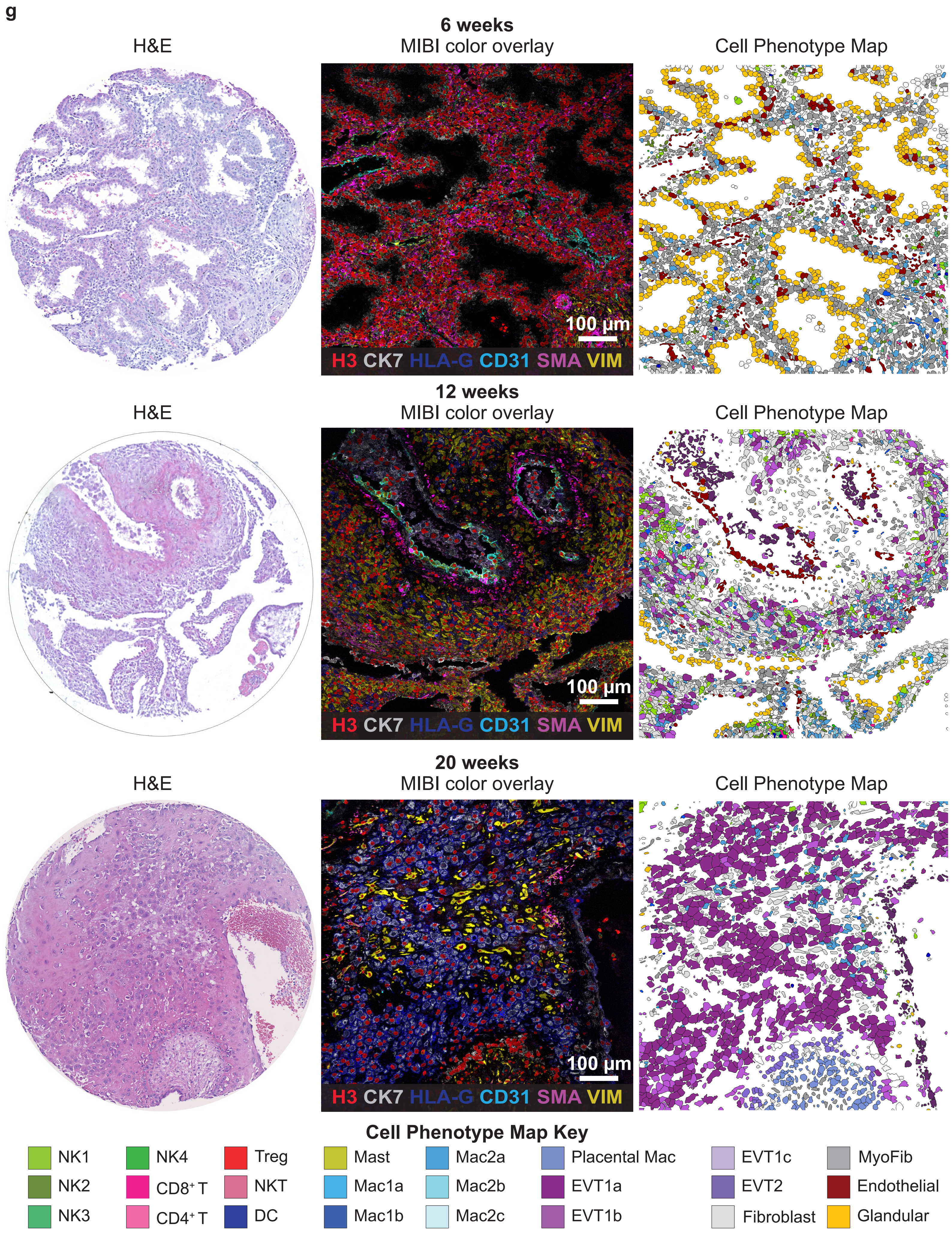
g
6 weeks
H&E
MIBI color overlay
Cell Phenotype Map
100 µm
H3 CK7 HLA-G CD31 SMA VIM
12 weeks
H&E
MIBI color overlay
Cell Phenotype Map
100 µm
H3 CK7 HLA-G CD31 SMA VIM
20 weeks
H&E
MIBI color overlay
Cell Phenotype Map
100 µm
H3 CK7 HLA-G CD31 SMA VIM
Cell Phenotype Map Key
NK1
NK2
NK3
NK4
CD8⁺ T
CD4⁺ T
Treg
NKT
DC
Mast
Mac1a
Mac1b
Mac2a
Mac2b
Mac2c
Placental Mac
EVT1a
EVT1b
EVT1c
EVT2
Fibroblast
MyoFib
Endothelial
Glandular

h

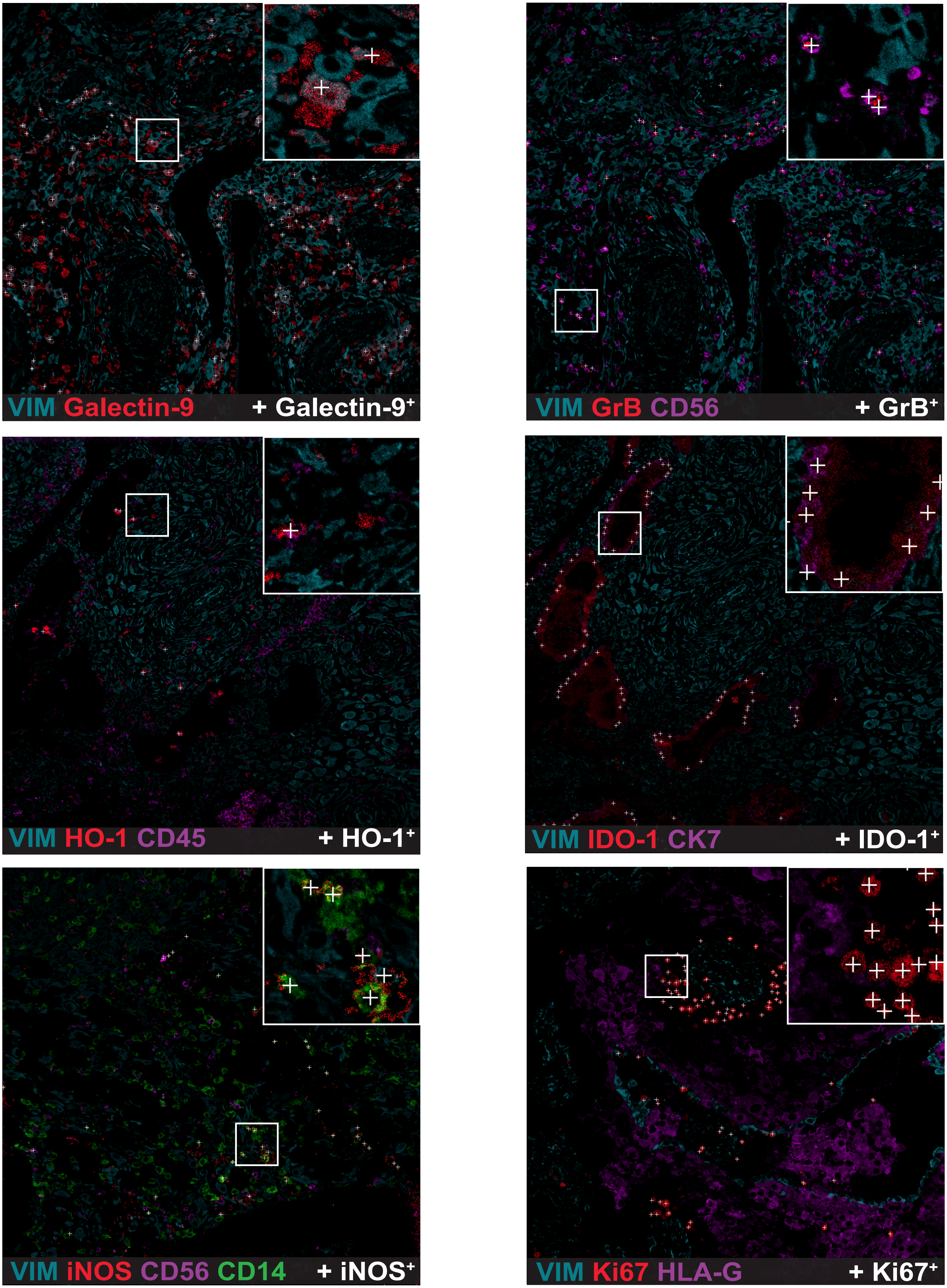
VIM Galectin-9
+ Galectin-9+
VIM GrB CD56
+ GrB+
VIM HO-1 CD45
+ HO-1+
VIM IDO-1 CK7
+ IDO-1+
VIM iNOS CD56 CD14
+ iNOS+
VIM Ki67 HLA-G
+ Ki67+

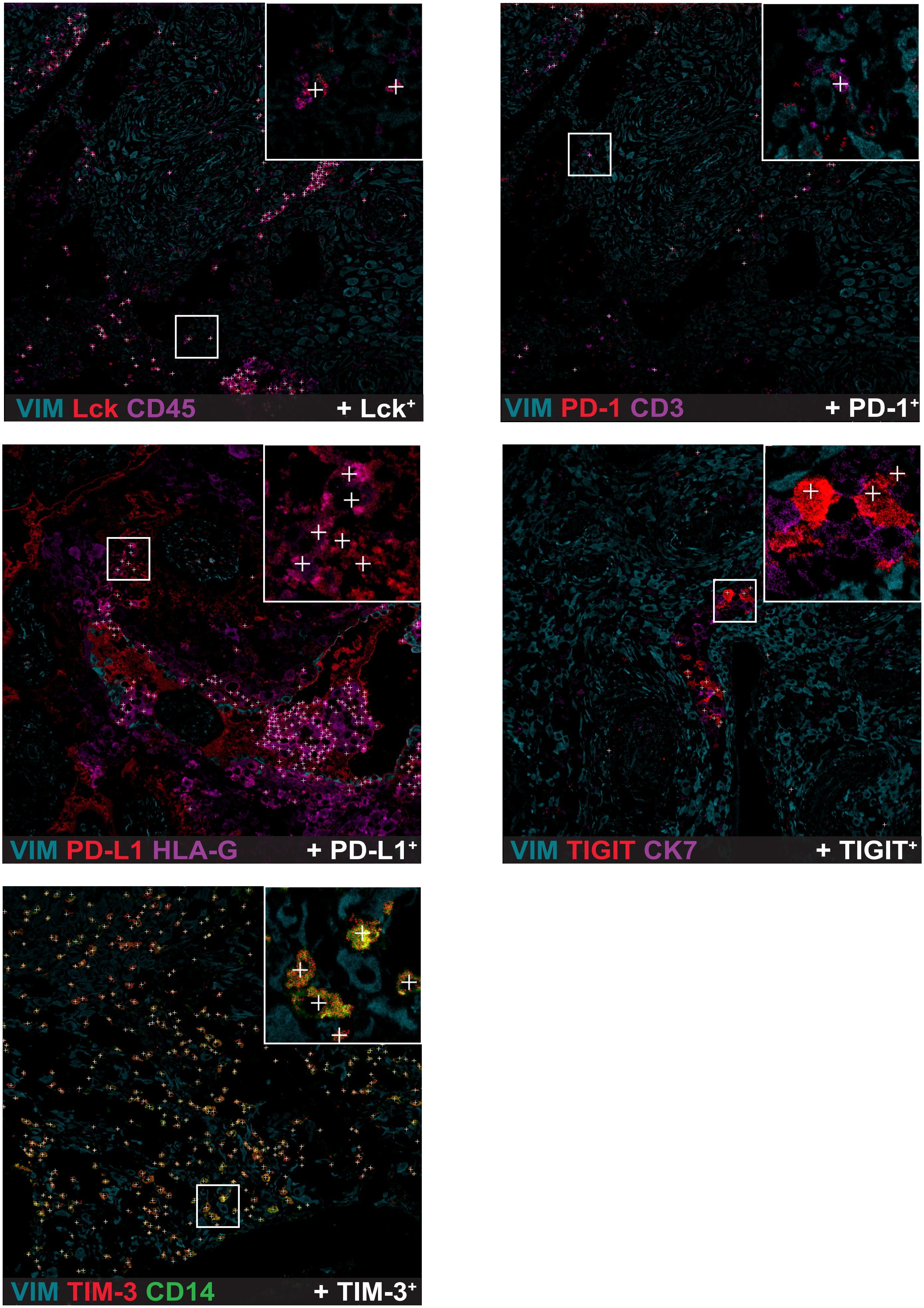
VIM Lck CD45
+ Lck+
VIM PD-1 CD3
+ PD-1+
VIM PD-L1 HLA-G
+ PD-L1+
VIM TIGIT CK7
+ TIGIT+
VIM TIM-3 CD14
+ TIM-3+

i

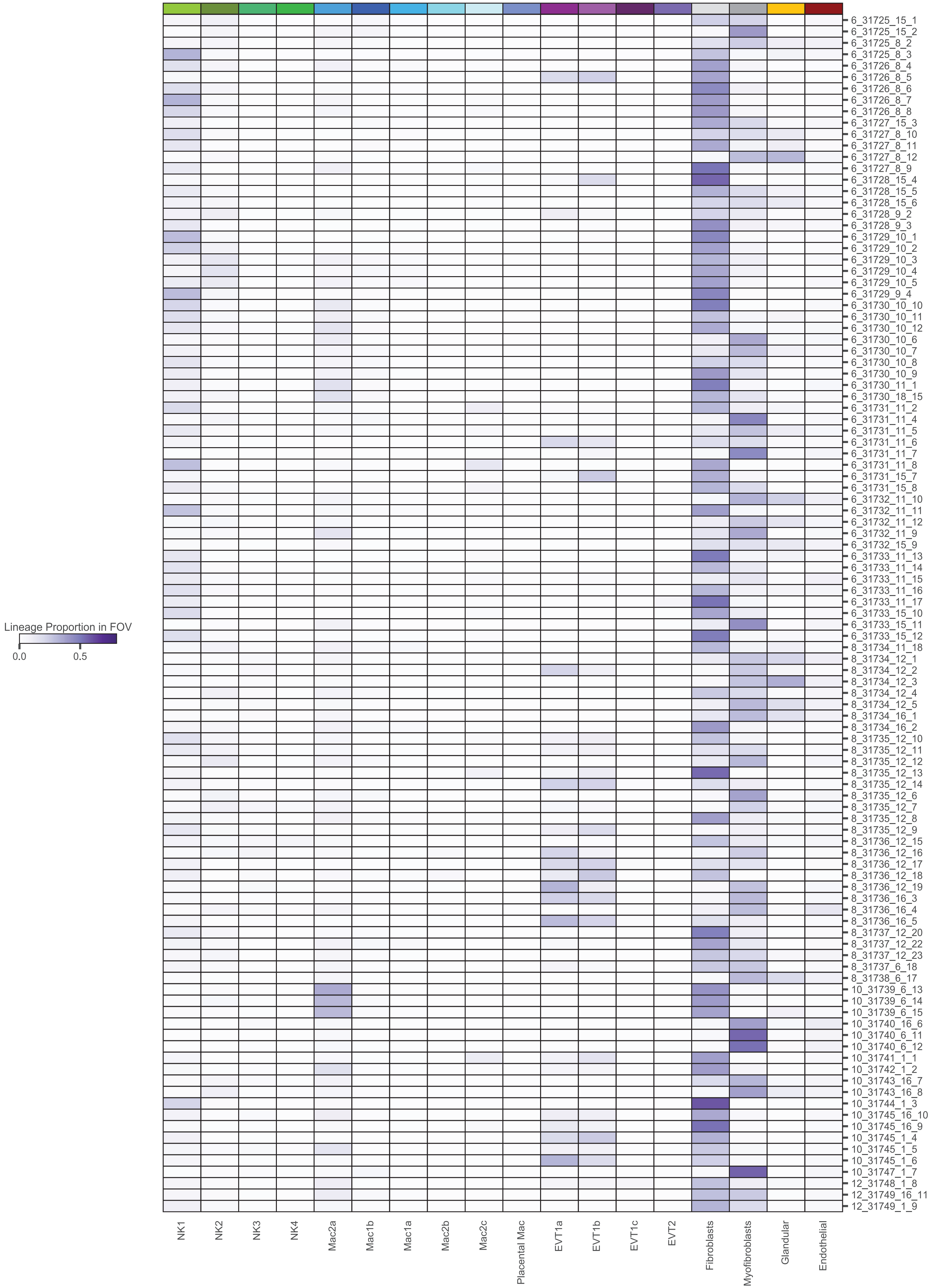
Lineage Proportion in FOV
0.0
0.5
NK1
NK2
NK3
NK4
Mac2a
Mac1b
Mac1a
Mac2b
Mac2c
Placental Mac
EVT1a
EVT1b
EVT1c
EVT2
Fibroblasts
Myofibroblasts
Glandular
Endothelial
6_31725_15_1
6_31725_15_2
6_31725_8_2
6_31725_8_3
6_31726_8_4
6_31726_8_5
6_31726_8_6
6_31726_8_7
6_31726_8_8
6_31727_15_3
6_31727_8_10
6_31727_8_11
6_31727_8_12
6_31727_8_9
6_31728_15_4
6_31728_15_5
6_31728_15_6
6_31728_9_2
6_31728_9_3
6_31729_10_1
6_31729_10_2
6_31729_10_3
6_31729_10_4
6_31729_10_5
6_31729_9_4
6_31730_10_10
6_31730_10_11
6_31730_10_12
6_31730_10_6
6_31730_10_7
6_31730_10_8
6_31730_10_9
6_31730_11_1
6_31730_18_15
6_31731_11_2
6_31731_11_4
6_31731_11_5
6_31731_11_6
6_31731_11_7
6_31731_11_8
6_31731_15_7
6_31731_15_8
6_31732_11_10
6_31732_11_11
6_31732_11_12
6_31732_11_9
6_31732_15_9
6_31733_11_13
6_31733_11_14
6_31733_11_15
6_31733_11_16
6_31733_11_17
6_31733_15_10
6_31733_15_11
6_31733_15_12
8_31734_11_18
8_31734_12_1
8_31734_12_2
8_31734_12_3
8_31734_12_4
8_31734_12_5
8_31734_16_1
8_31734_16_2
8_31735_12_10
8_31735_12_11
8_31735_12_12
8_31735_12_13
8_31735_12_14
8_31735_12_6
8_31735_12_7
8_31735_12_8
8_31735_12_9
8_31736_12_15
8_31736_12_16
8_31736_12_17
8_31736_12_18
8_31736_12_19
8_31736_16_3
8_31736_16_4
8_31736_16_5
8_31737_12_20
8_31737_12_22
8_31737_12_23
8_31737_6_18
8_31738_6_17
10_31739_6_13
10_31739_6_14
10_31739_6_15
10_31740_16_6
10_31740_6_11
10_31740_6_12
10_31741_1_1
10_31742_1_2
10_31743_16_7
10_31743_16_8
10_31744_1_3
10_31745_16_10
10_31745_16_9
10_31745_1_4
10_31745_1_5
10_31745_1_6
10_31747_1_7
12_31748_1_8
12_31749_16_11
12_31749_1_9

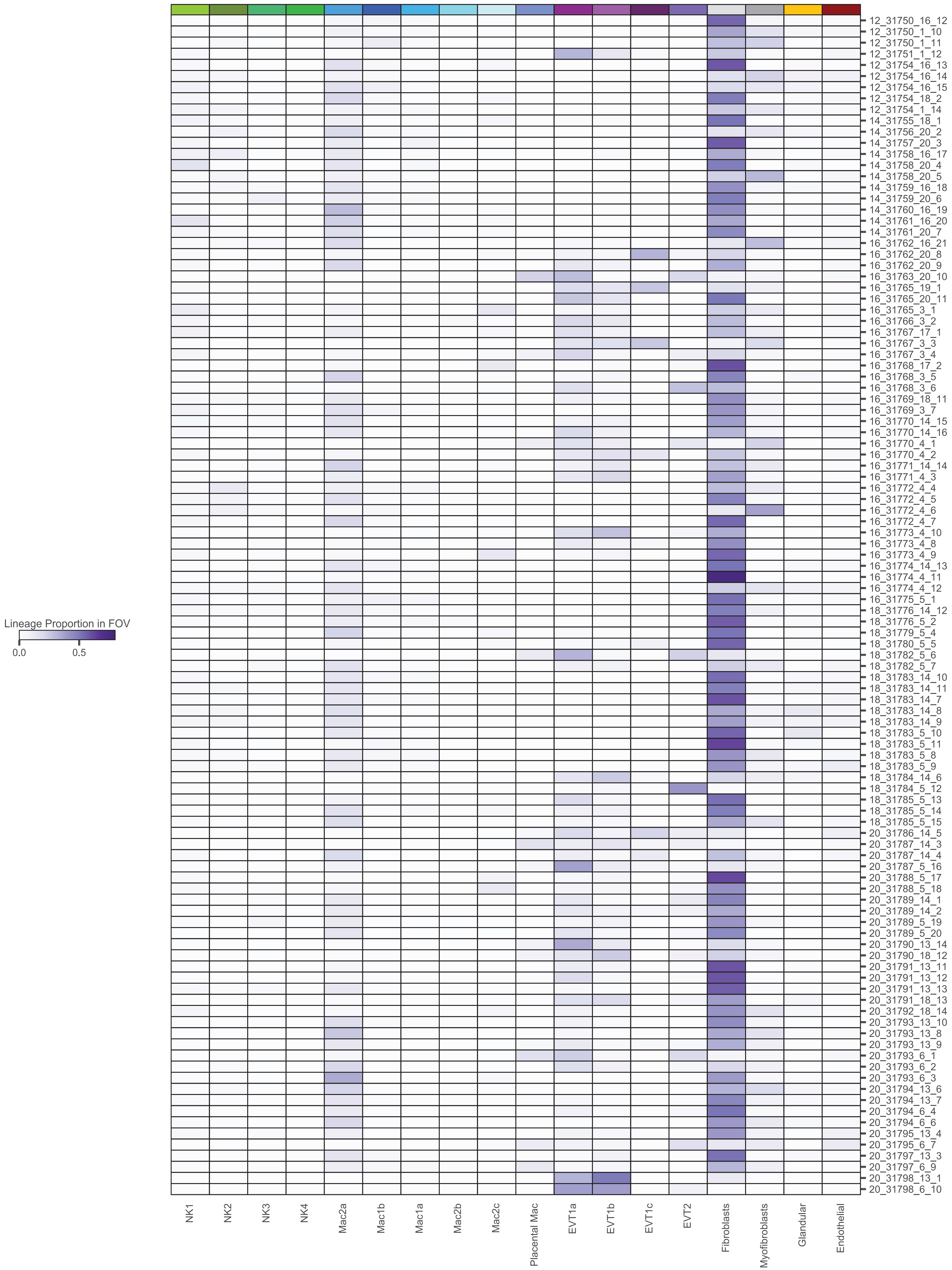
Lineage Proportion in FOV
0.0
0.5
12_31750_16_12
12_31750_1_10
12_31750_1_11
12_31751_1_12
12_31754_16_13
12_31754_16_14
12_31754_16_15
12_31754_18_2
12_31754_1_14
14_31755_18_1
14_31756_20_2
14_31757_20_3
14_31758_16_17
14_31758_20_4
14_31758_20_5
14_31759_16_18
14_31759_20_6
14_31760_16_19
14_31761_16_20
14_31761_20_7
16_31762_16_21
16_31762_20_8
16_31762_20_9
16_31763_20_10
16_31765_19_1
16_31765_20_11
16_31765_3_1
16_31766_3_2
16_31767_17_1
16_31767_3_3
16_31767_3_4
16_31768_17_2
16_31768_3_5
16_31768_3_6
16_31769_18_11
16_31769_3_7
16_31770_14_15
16_31770_14_16
16_31770_4_1
16_31770_4_2
16_31771_14_14
16_31771_4_3
16_31772_4_4
16_31772_4_5
16_31772_4_6
16_31772_4_7
16_31773_4_10
16_31773_4_8
16_31773_4_9
16_31774_14_13
16_31774_4_11
16_31774_4_12
16_31775_5_1
18_31776_14_12
18_31776_5_2
18_31779_5_4
18_31780_5_5
18_31782_5_6
18_31782_5_7
18_31783_14_10
18_31783_14_11
18_31783_14_7
18_31783_14_8
18_31783_14_9
18_31783_5_10
18_31783_5_11
18_31783_5_8
18_31783_5_9
18_31784_14_6
18_31784_5_12
18_31785_5_13
18_31785_5_14
18_31785_5_15
20_31786_14_5
20_31787_14_3
20_31787_14_4
20_31787_5_16
20_31788_5_17
20_31788_5_18
20_31789_14_1
20_31789_14_2
20_31789_5_19
20_31789_5_20
20_31790_13_14
20_31790_18_12
20_31791_13_11
20_31791_13_12
20_31791_13_13
20_31791_18_13
20_31792_18_14
20_31793_13_10
20_31793_13_8
20_31793_13_9
20_31793_6_1
20_31793_6_2
20_31793_6_3
20_31794_13_6
20_31794_13_7
20_31794_6_4
20_31794_6_6
20_31795_13_4
20_31795_6_7
20_31797_13_3
20_31797_6_9
20_31798_13_1
20_31798_6_10
NK1
NK2
NK3
NK4
Mac2a
Mac1b
Mac1a
Mac2b
Mac2c
Placental Mac
EVT1a
EVT1b
EVT1c
EVT2
Fibroblasts
Myofibroblasts
Glandular
Endothelial

**Extended Data Figure S2 | Workflow for Mesmer whole-cell segmentation of single cells from MIBI images and cell phenotyping. a.** Workflow. (1) Input of manually annotated decidua MIBI images to train our custom convolutional neural network (CNN) Mesmer. (2) Six-channel inputs to Mesmer and outputs (probability maps for cell shape and the center of each cell) with subsequent watershed transformation producing labeled, segmented images. (3) Post-processing parameters used to more accurately represent marker expression by cell populations located in the glands and anchoring (cell column) villi. VIM, vimentin. (4) Pseudocolored MIBI overlay showing final whole-cell segmentation of a representative MIBI image. **b.** Overview of steps involved in single-cell phenotyping. (Left to right) Output from FlowSOM (100 clusters) and subsequent assignment to cell phenotypes. **c.** Number of cells assigned to each FlowSOM cluster **d.** Histograms of normalized marker expression distributions for FOXP3, CD11c, tryptase, and CD14, with thresholds (vertical line) used for cell phenotyping. **e.** UMAP of dataset with cell phenotype assignments overlaid **f.** UMAPs of dataset with marker expression overlaid. **g.** Representative H&E and MIBI serial sections with cell phenotype maps. Each cell is colored by its phenotype assignment for GAs 6, 12, and 20 weeks. VIM, vimentin; SMA, smooth muscle actin; Placental Mac, Placental Macrophage; MyoFib, Myofibroblast. **h.** Representative MIBI overlays showing positive functional marker expression. White crosses indicate cells positive for each functional marker according to the set binary threshold. **i.** Heatmap showing proportion of cells belonging to each assigned lineage in each FOV.

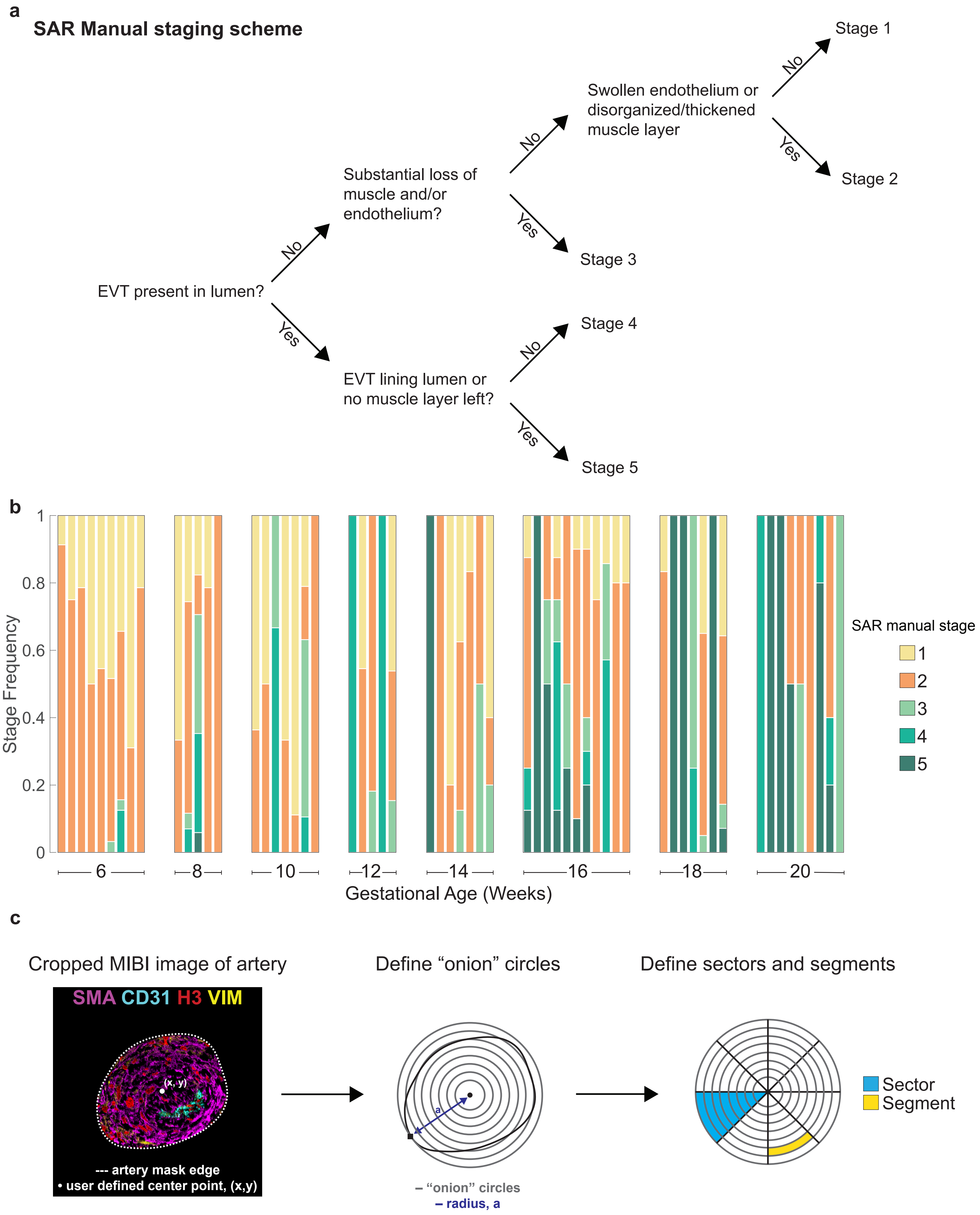
a
SAR Manual staging scheme
EVT present in lumen?
No
Yes
Substantial loss of muscle and/or endothelium?
EVT lining lumen or no muscle layer left?
Swollen endothelium or disorganized/thickened muscle layer
Stage 1
Stage 2
Stage 3
Stage 4
Stage 5
b
Stage Frequency
Gestational Age (Weeks)
6
8
10
12
14
16
18
20
SAR manual stage
1
2
3
4
5
c
Cropped MIBI image of artery
SMA CD31 H3 VIM
(x, y)
--- artery mask edge
• user defined center point, (x,y)
Define "onion" circles
– "onion" circles
– radius, a
Define sectors and segments
Sector
Segment

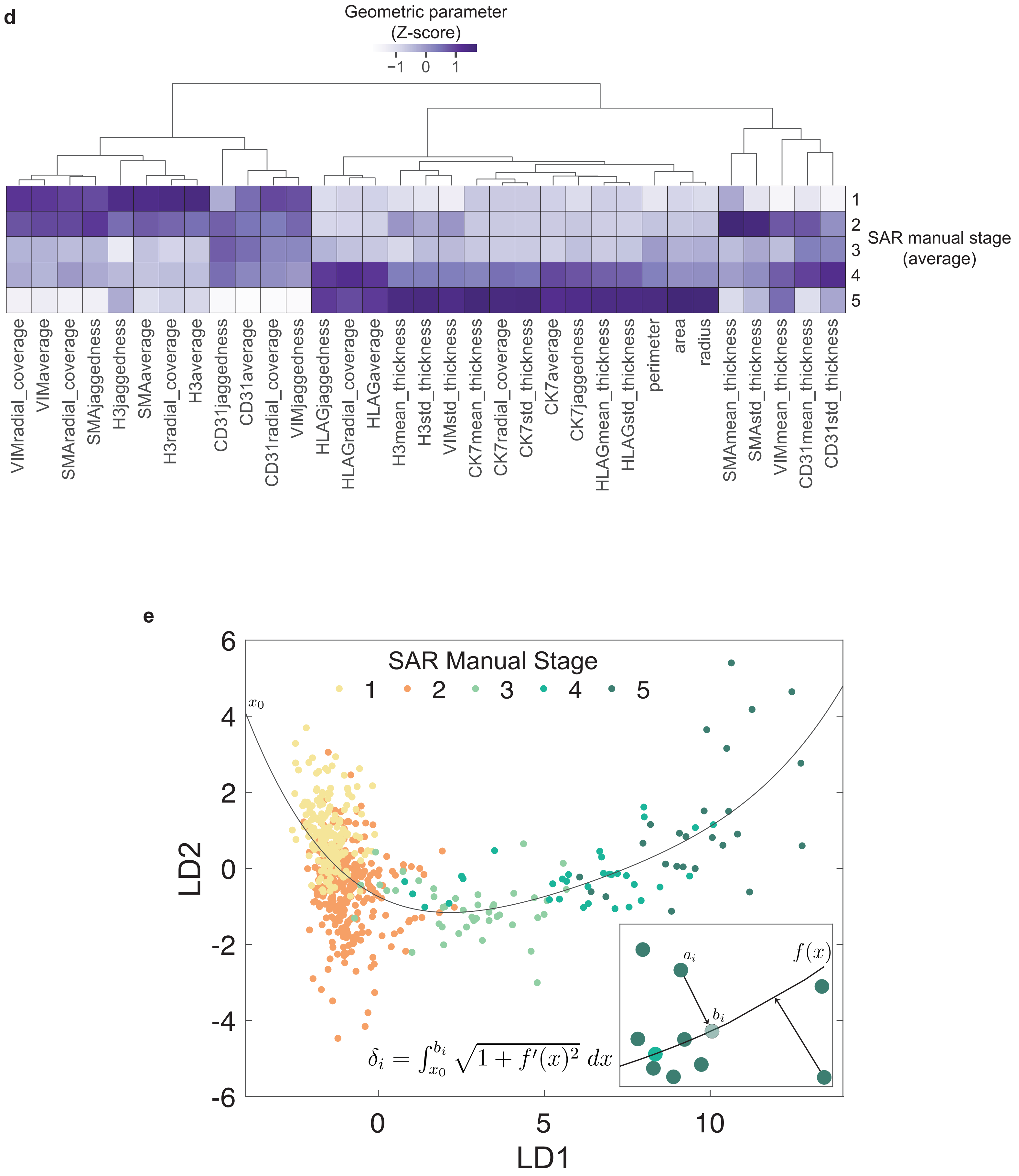

d
Geometric parameter (Z-score)
−1 0 1
SAR manual stage (average)
1
2
3
4
5
VIMradial_coverage
VIMaverage
SMAradial_coverage
SMAjaggedness
H3jaggedness
SMAaverage
H3radial_coverage
H3average
CD31jaggedness
CD31average
CD31radial_coverage
VIMjaggedness
HLAGjaggedness
HLAGradial_coverage
HLAGaverage
H3mean_thickness
H3std_thickness
VIMstd_thickness
CK7mean_thickness
CK7radial_coverage
CK7std_thickness
CK7average
CK7jaggedness
HLAGmean_thickness
HLAGstd_thickness
perimeter
area
radius
SMAmean_thickness
SMAstd_thickness
VIMmean_thickness
CD31mean_thickness
CD31std_thickness
e
SAR Manual Stage
1 2 3 4 5
LD2
LD1
6
4
2
0
-2
-4
-6
0
5
10
$x_0$
$a_i$
$b_i$
$f(x)$
$\delta_i = \int_{x_0}^{b_i} \sqrt{1 + f'(x)^2}\, dx$

**f**

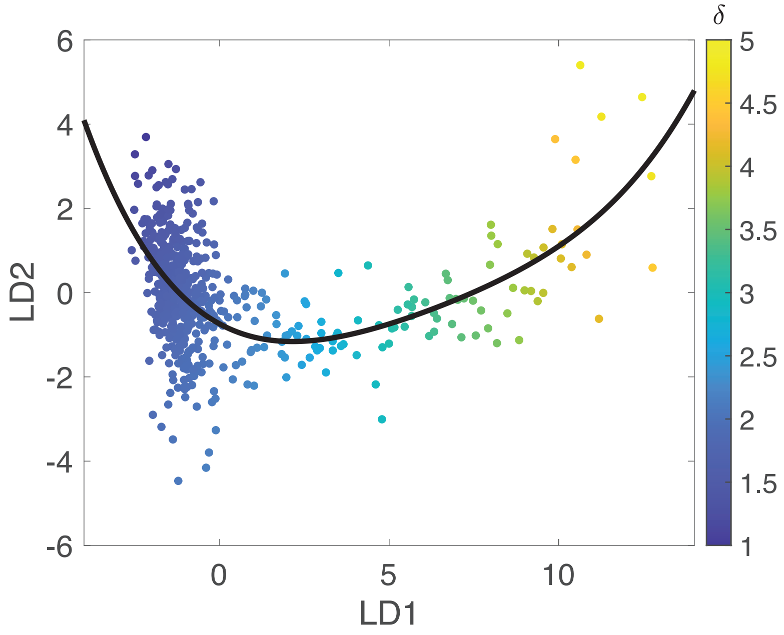

δ
6
4
2
0
-2
-4
-6
LD2
0
5
10
LD1
5
4.5
4
3.5
3
2.5
2
1.5
1


**g**

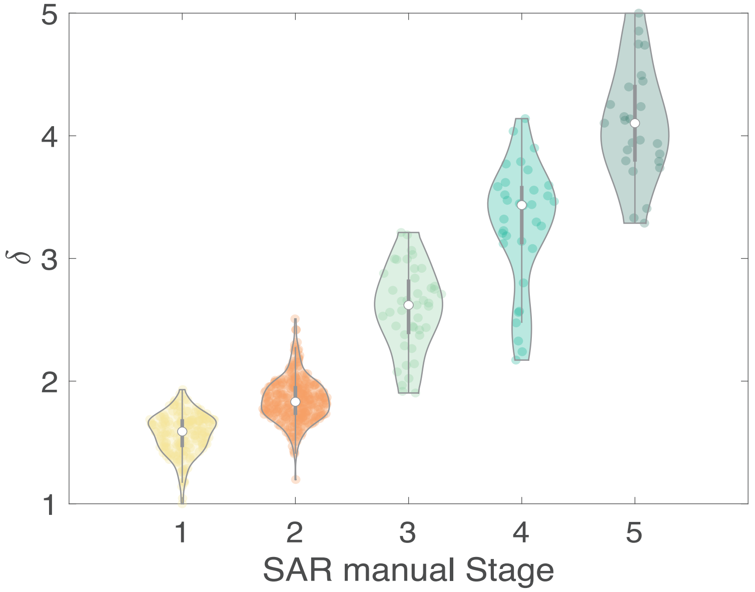

5
4
3
2
1
δ
1
2
3
4
5
SAR manual Stage

**h**

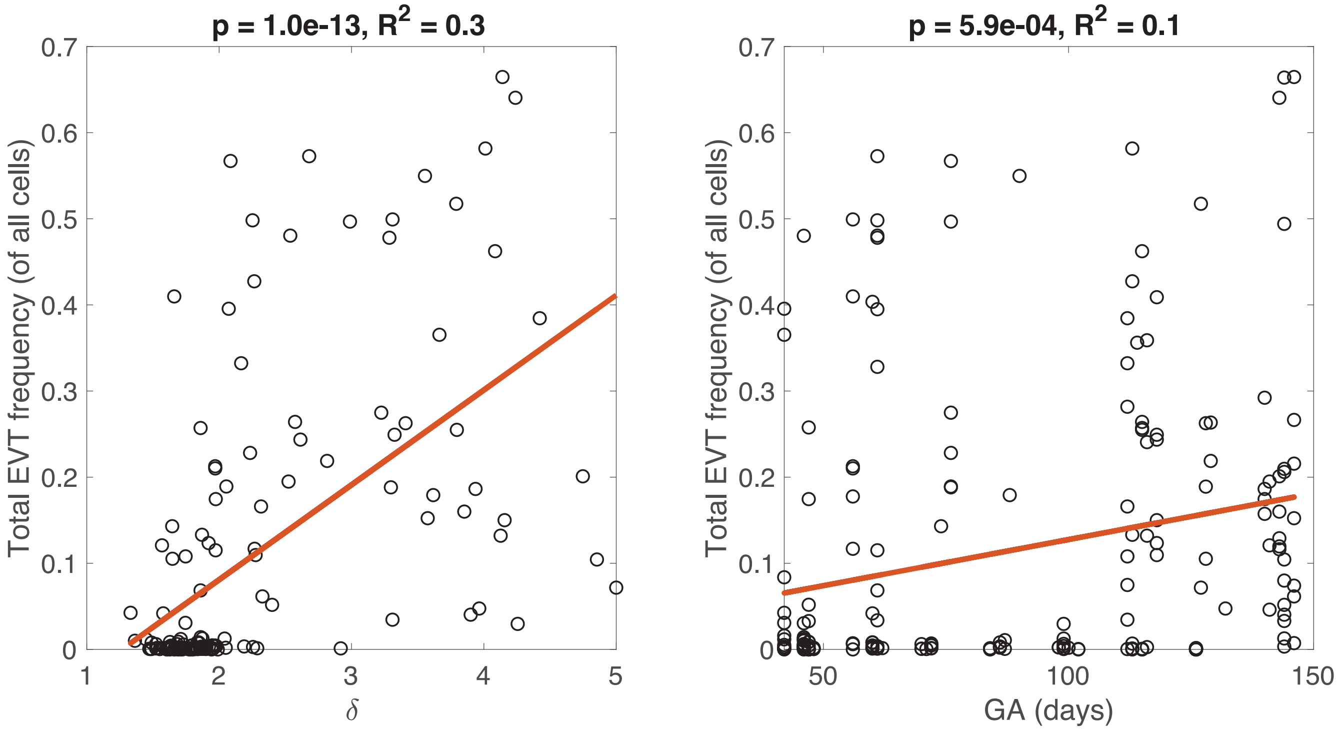
p = 1.0e-13, R2 = 0.3
Total EVT frequency (of all cells)
δ
p = 5.9e-04, R2 = 0.1
Total EVT frequency (of all cells)
GA (days)

**i**

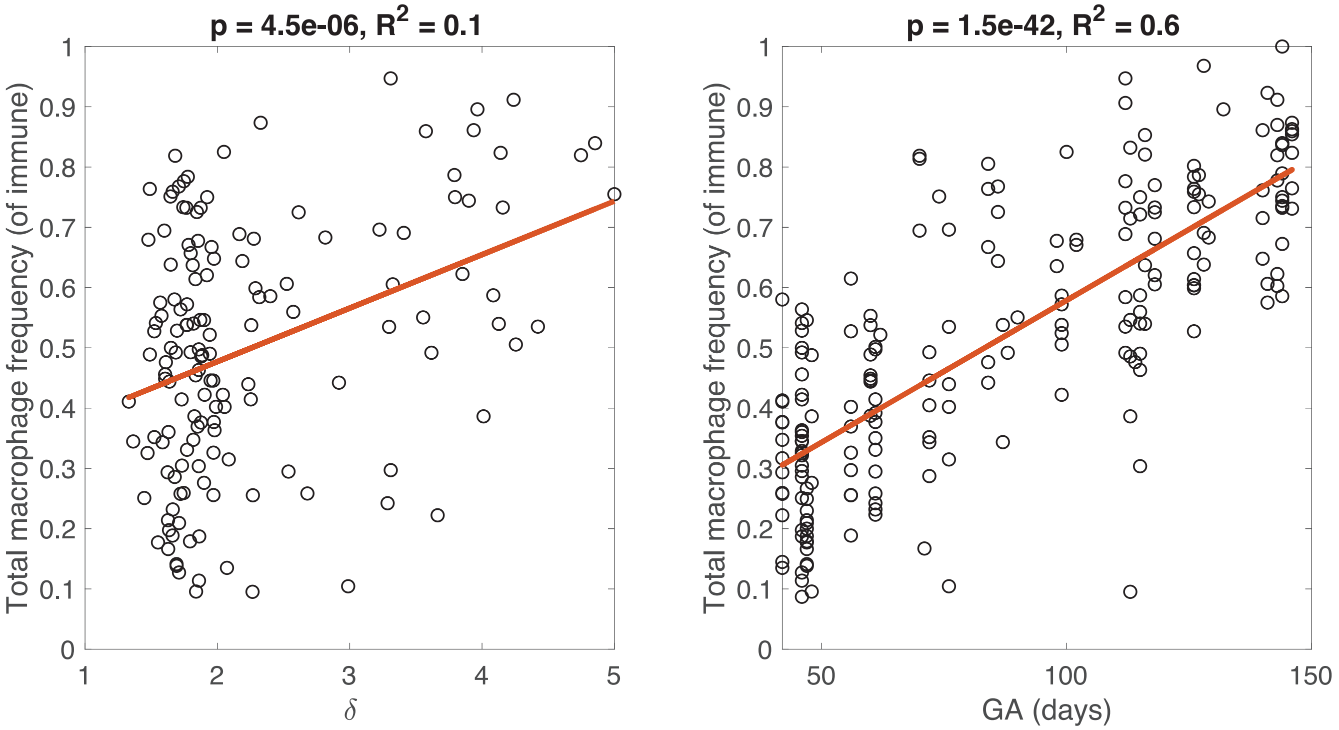
p = 4.5e-06, R2 = 0.1
Total macrophage frequency (of immune)
δ
p = 1.5e-42, R2 = 0.6
Total macrophage frequency (of immune)
GA (days)

**j**

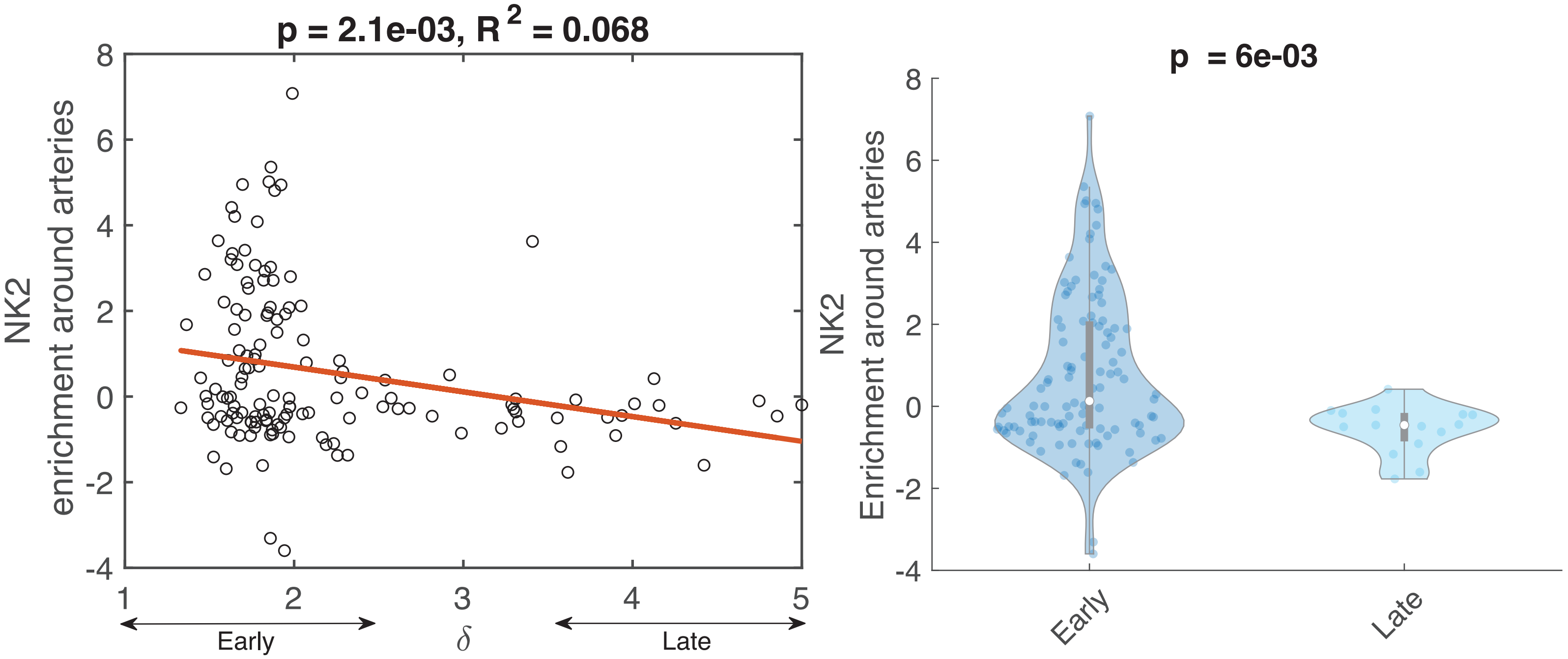
p = 2.1e-03, $R^2$ = 0.068
NK2
enrichment around arteries
Early
$\delta$
Late
p = 6e-03
NK2
Enrichment around arteries
Early
Late

**k**

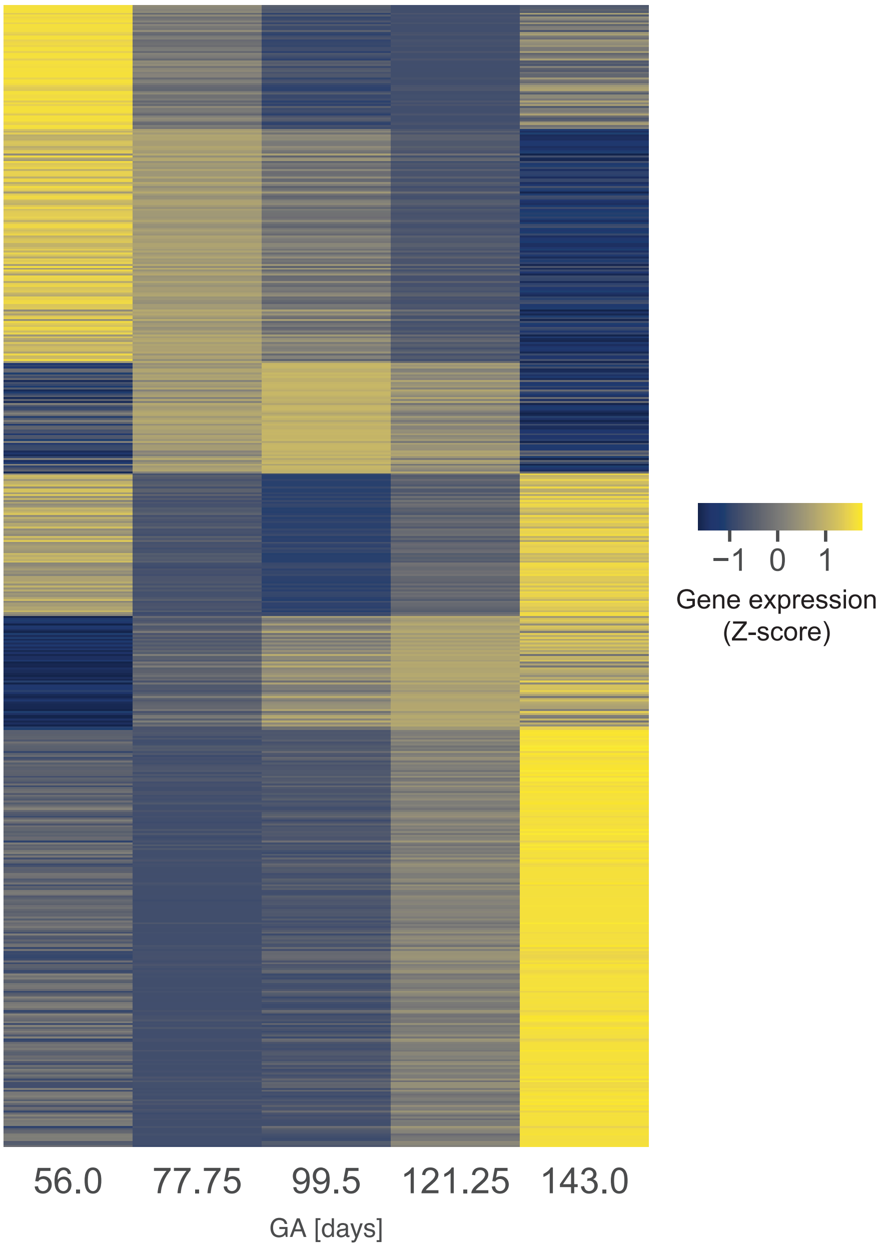
−1 0 1
Gene expression
(Z-score)
56.0
77.75
99.5
121.25
143.0
GA [days]

**I**

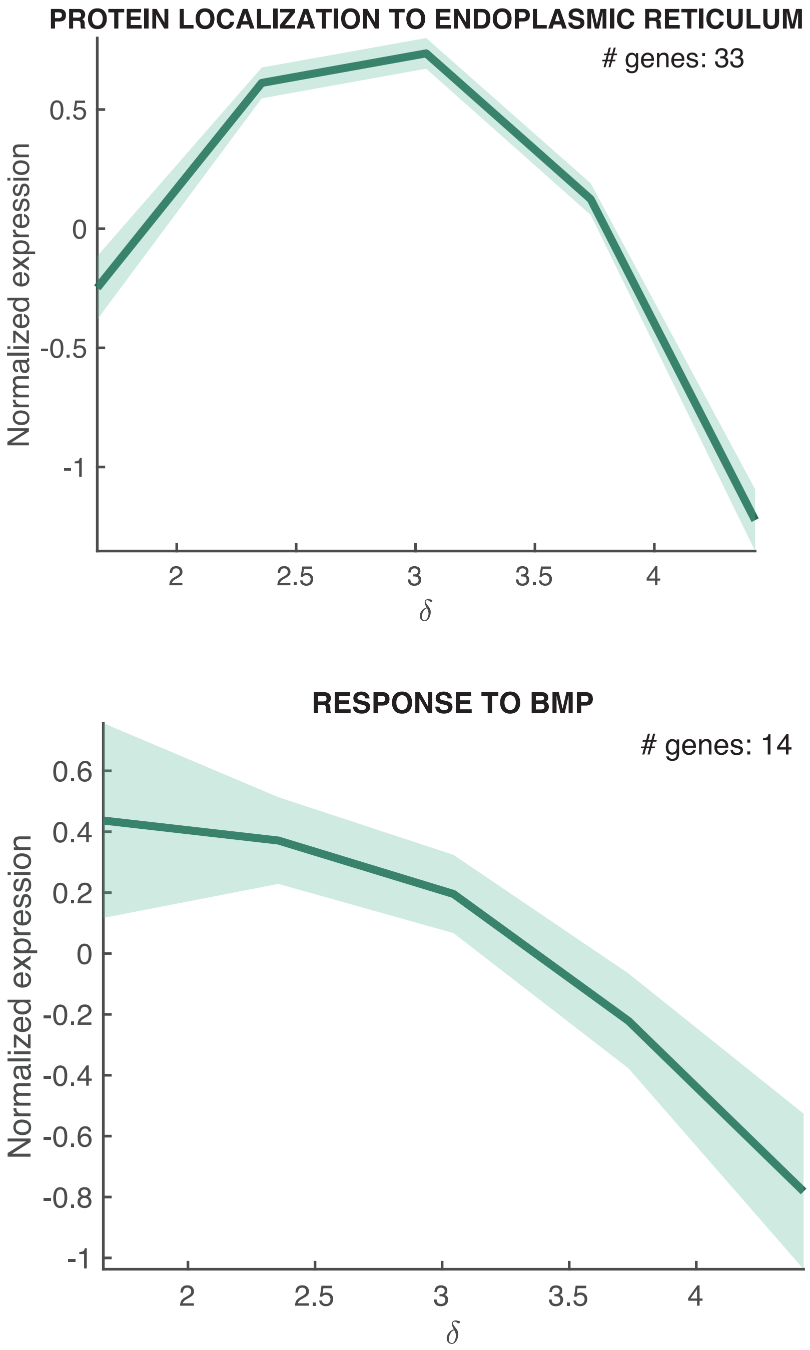
PROTEIN LOCALIZATION TO ENDOPLASMIC RETICULUM
### genes: 33
Normalized expression
0.5
0
-0.5
-1
2
2.5
3
3.5
4
δ
RESPONSE TO BMP
### genes: 14
Normalized expression
0.6
0.4
0.2
0
-0.2
-0.4
-0.6
-0.8
-1
2
2.5
3
3.5
4
δ

**Extended Data Figure S3 | Spiral Artery Remodeling (SAR) manual and digitized staging schemes. a.** Flowchart for manual staging of arteries used by a blinded reviewer for staging arteries. Criteria were modified for MIBI images from Smith et al[42]. **b.** Bar plot of the frequency of artery stages manually determined per patient. **c.** Schematic showing digitization of artery morphology features. **d.** Heatmap of quantified digitized morphological features, mean across manual SAR stages. Columns were Z-scored and hierarchically clustered. **e.** Scatter of arteries in LDA space color coded by manually assigned stage. The polynomial fit depicts the remodeling trajectory. Inset: matching each artery point $a_i$ to the SAR trajectory by finding the nearest point along trajectory $b_i$. The continuous SAR score $\delta$ was then defined as the distance from origin $x_0$ to $b_i$ along the trajectory curve. **f.** Scatter plot of arteries in LDA space color coded by SAR continuous score $\delta$. **g.** Violin plot of the distribution of SAR continuous score $\delta$ by manual stage, p-value comparing the 5 distributions **h.** Scatter plot of total EVT frequency (out of all cells) by SAR continuous score $\delta$ (left) and GA (right). Red line, fitted linear regression model. **i.** Scatter plot of total macrophage frequency (out of immune cells) by SAR continuous score $\delta$ (left) and GA (right). Red line, fitted linear regression model. **j.** Pairwise enrichment between NK2 cells and arteries. Left: mean enrichment per image by SAR continuous score $\delta$. Right: Distribution of NK2-artery enrichment scores by early ($1 \leq \delta < 3$) and late ($3 \leq \delta < 5$) remodeling stage. **k.** GA-correlated genes in decidua showing mean normalized expression (Z-score) by binned GA. Genes shown are only those most correlated with GA after comparing each gene against all predictors (GA, SAR, cell type frequencies; Supplemental Table 16) **l.** 2 SAR ($\delta$)-trending in artery gene ontology pathways, showing normalized expression of genes in the GO pathway by SAR ($\delta$).

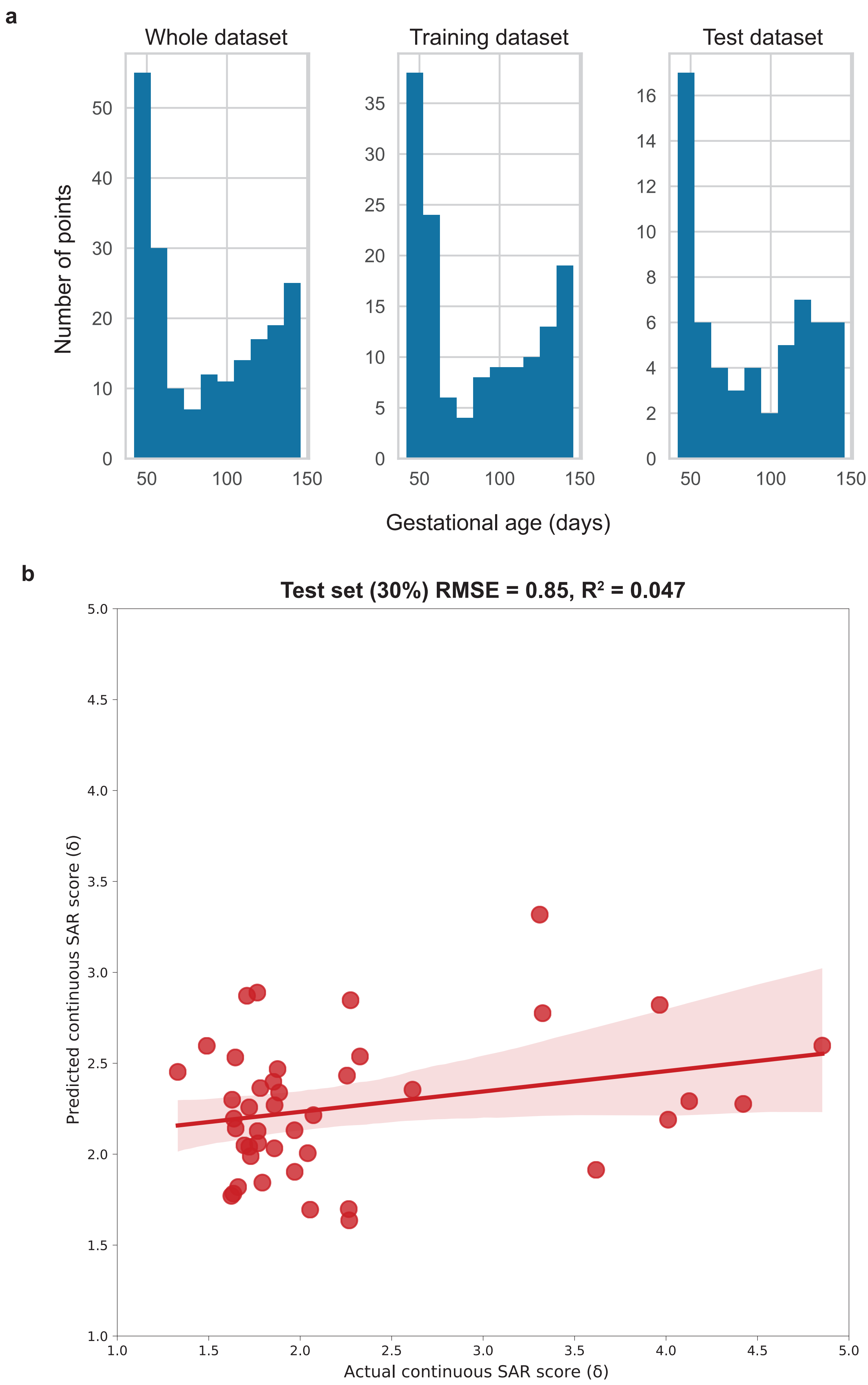
a
Whole dataset
Training dataset
Test dataset
Number of points
Gestational age (days)
b
Test set (30%) RMSE = 0.85, R² = 0.047
Predicted continuous SAR score (δ)
Actual continuous SAR score (δ)

**Extended Data Figure S4 | Ridge regression for predicting gestational age (GA) from immune composition. a.** Distribution of GA (in days) across the whole dataset, training (70%) set, and test (30%) set. **b.** Predicted versus actual SAR continuous score $\delta$ for the ridge regression model (analogous to the model used to predict GA in days) trained on GA-associated immune features, for the withheld test set (30%). Shaded region, 1 standard deviation.

**a**

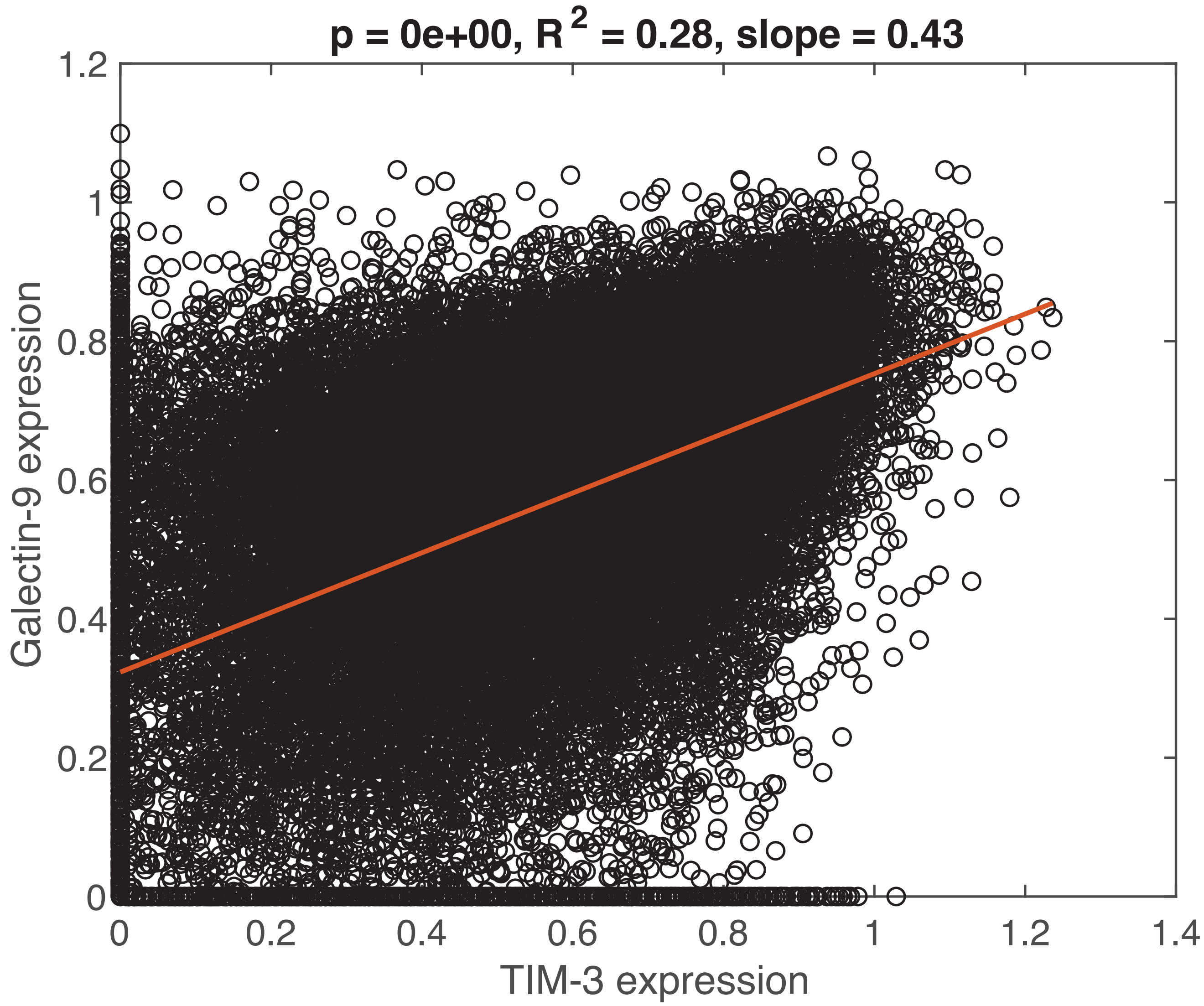

p = 0e+00, R 2 = 0.28, slope = 0.43
1.2
1
0.8
0.6
0.4
0.2
0
Galectin-9 expression
0
0.2
0.4
0.6
0.8
1
1.2
1.4
TIM-3 expression

**Extended Data Figure S5 | Single-cell expression of immunoregulatory markers. a.** Scatter plot of the expression of TIM-3 and galectin-9 in individual Mac2a and Mac2b cells from all images. Red line, fitted linear regression model.

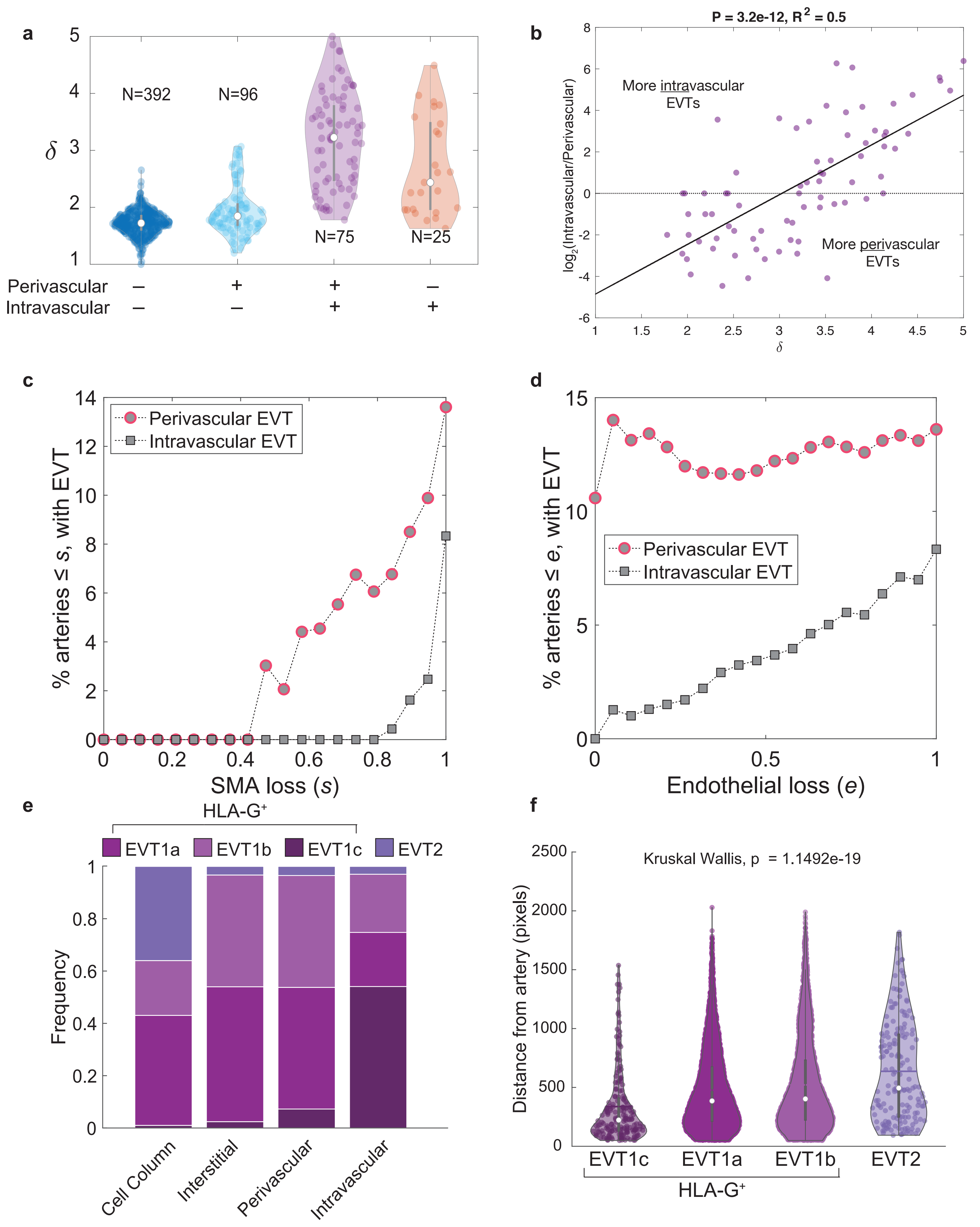

a
b
c
d
e
f
N=392
N=96
N=75
N=25
δ
Perivascular
Intravascular
P = 3.2e-12, R² = 0.5
More intravascular EVTs
More perivascular EVTs
log2(Intravascular/Perivascular)
% arteries ≤ s, with EVT
SMA loss (s)
Perivascular EVT
Intravascular EVT
% arteries ≤ e, with EVT
Endothelial loss (e)
HLA-G⁺
EVT1a
EVT1b
EVT1c
EVT2
Frequency
Cell Column
Interstitial
Perivascular
Intravascular
Kruskal Wallis, p = 1.1492e-19
Distance from artery (pixels)
EVT1c
EVT1a
EVT1b
EVT2
HLA-G⁺

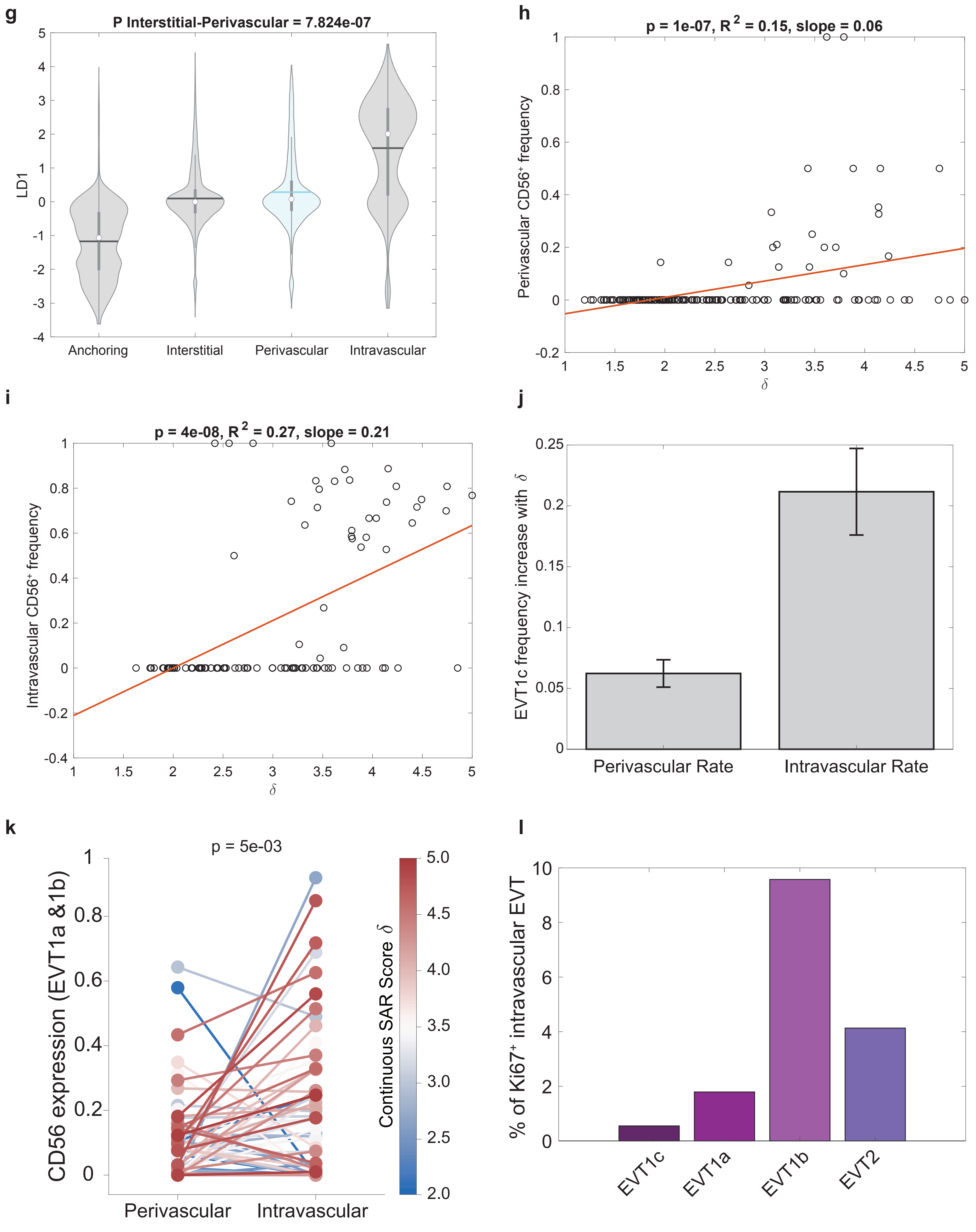
g
P Interstitial-Perivascular = 7.824e-07
LD1
Anchoring
Interstitial
Perivascular
Intravascular
h
p = 1e-07, R 2 = 0.15, slope = 0.06
Perivascular CD56⁺ frequency
δ
i
p = 4e-08, R 2 = 0.27, slope = 0.21
Intravascular CD56⁺ frequency
δ
j
EVT1c frequency increase with δ
Perivascular Rate
Intravascular Rate
k
p = 5e-03
CD56 expression (EVT1a &1b)
Continuous SAR Score δ
Perivascular
Intravascular
l
% of Ki67⁺ intravascular EVT
EVT1c
EVT1a
EVT1b
EVT2

**m**

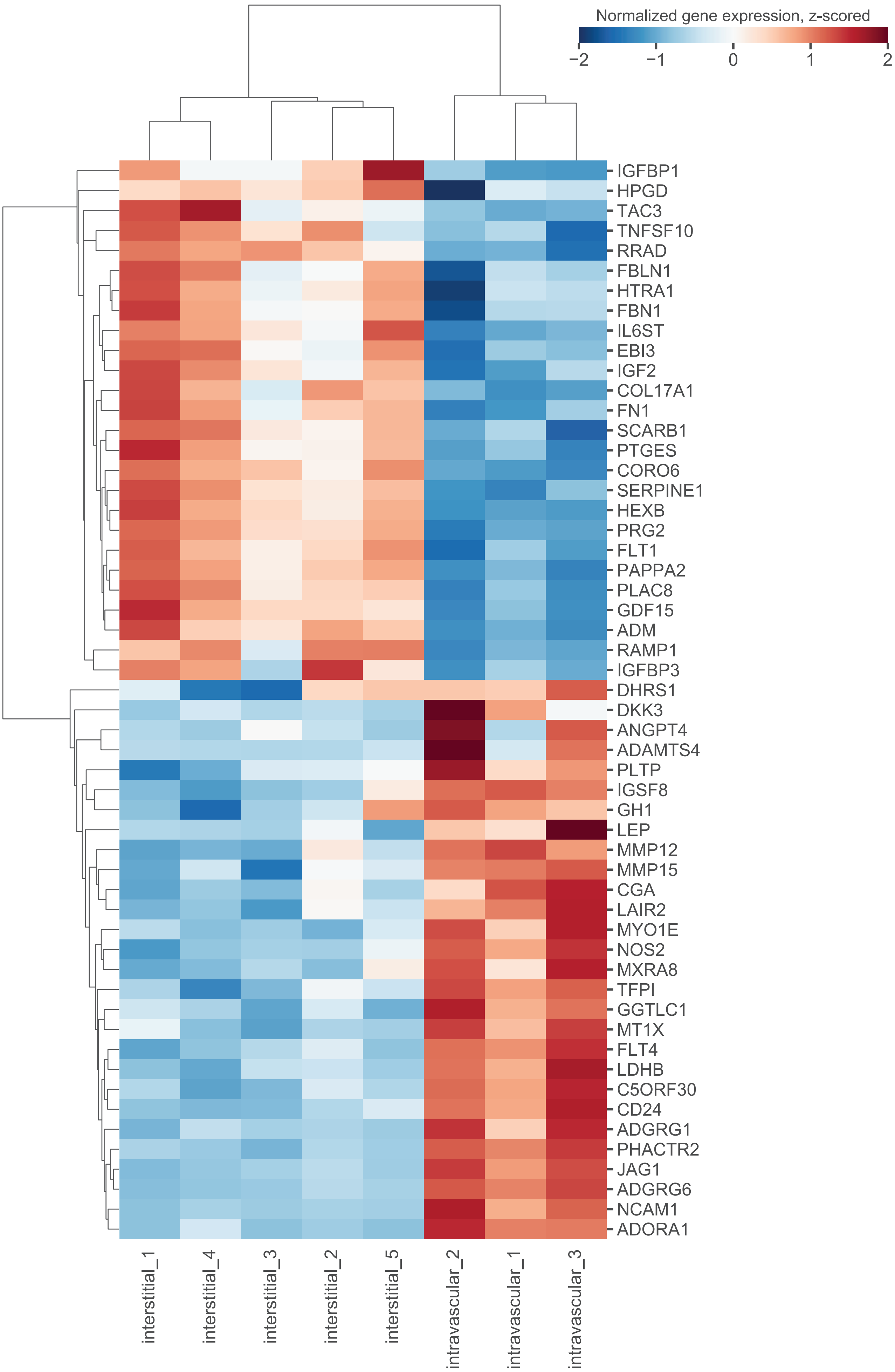

Normalized gene expression, z-scored
−2
−1
0
1
2
IGFBP1
HPGD
TAC3
TNFSF10
RRAD
FBLN1
HTRA1
FBN1
IL6ST
EBI3
IGF2
COL17A1
FN1
SCARB1
PTGES
CORO6
SERPINE1
HEXB
PRG2
FLT1
PAPPA2
PLAC8
GDF15
ADM
RAMP1
IGFBP3
DHRS1
DKK3
ANGPT4
ADAMTS4
PLTP
IGSF8
GH1
LEP
MMP12
MMP15
CGA
LAIR2
MYO1E
NOS2
MXRA8
TFPI
GGTLC1
MT1X
FLT4
LDHB
C5ORF30
CD24
ADGRG1
PHACTR2
JAG1
ADGRG6
NCAM1
ADORA1
interstitial_1
interstitial_4
interstitial_3
interstitial_2
interstitial_5
intravascular_2
intravascular_1
intravascular_3

**n**

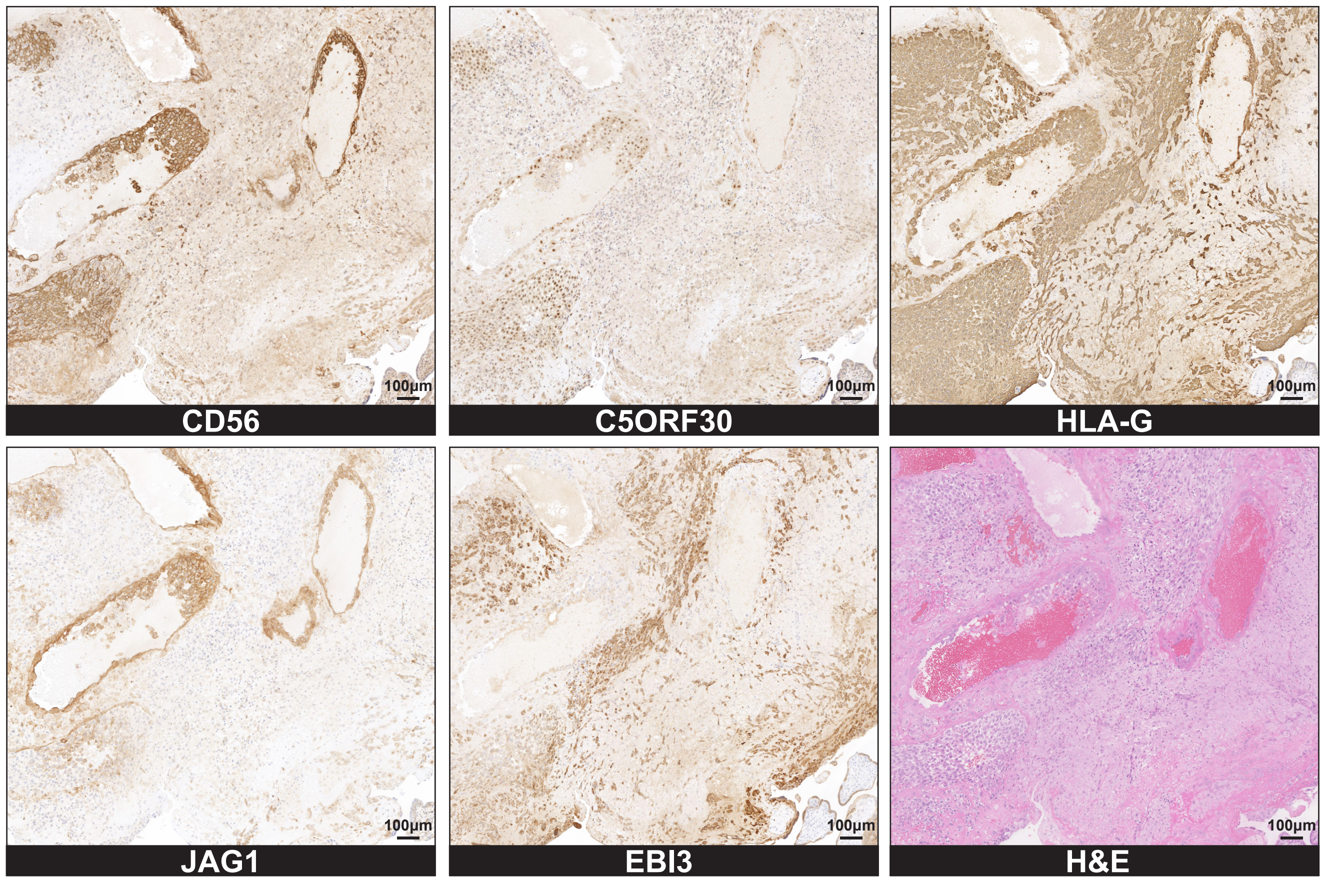
CD56
C5ORF30
HLA-G
JAG1
EBI3
H&E
100µm

**o**

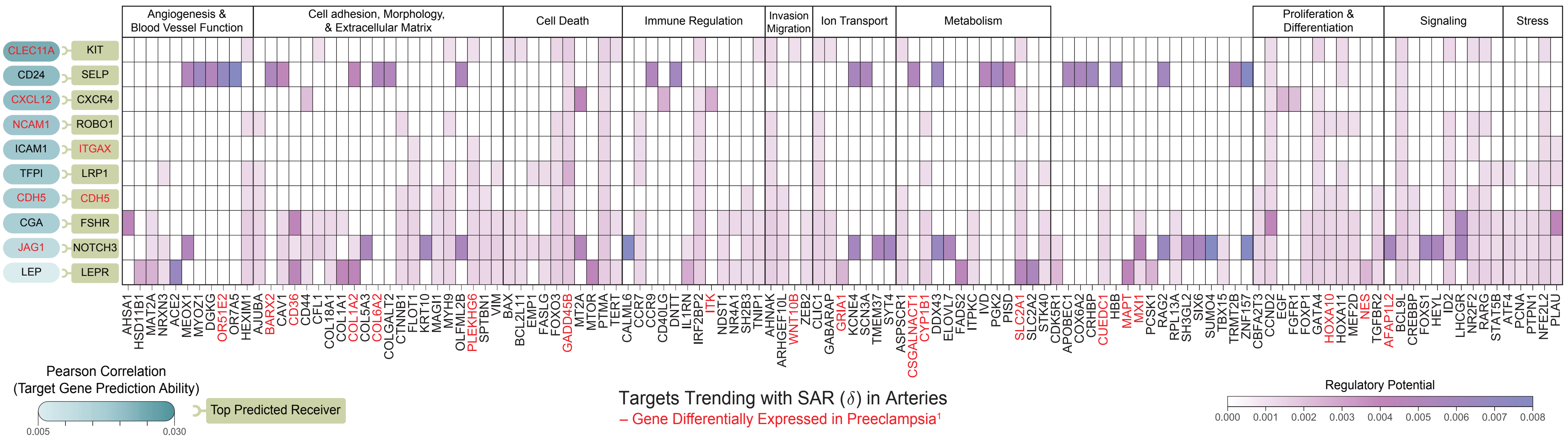
Angiogenesis & Blood Vessel Function
Cell adhesion, Morphology, & Extracellular Matrix
Cell Death
Immune Regulation
Invasion Migration
Ion Transport
Metabolism
Proliferation & Differentiation
Signaling
Stress
CLEC11A KIT
CD24 SELP
CXCL12 CXCR4
NCAM1 ROBO1
ICAM1 ITGAX
TFPI LRP1
CDH5 CDH5
CGA FSHR
JAG1 NOTCH3
LEP LEPR
AHSA1 HSD11B1 MAT2A NRXN3 ACE2 MEOX1 MYOZ1 DGKG OR51E2 OR7A5 HEXIM1
AJUBA BARX2 CAV1 CD36 CD44 CFL1 COL18A1 COL1A1 COL1A2 COL5A3 COL6A2 COLGALT2 CTNNB1 FLOT1 KRT10 MAGI1 MYH9 OLFML2B PLEKHG6 SPTBN1 VIM
BAX BCL2L11 EMP1 FASLG FOXO3 GADD45B MT2A MTOR PTMA TERT
CALML6 CCR7 CCR9 CD40LG DNTT IL1RN IRF2BP2 ITK NDST1 NR4A1 SH2B3 TNIP1
AHNAK ARHGEF10L WNT10B ZEB2
CLIC1 GABARAP GRIA1 KCNE4 SCN3A TMEM37 SYT4
ASPSCR1 CSGALNACT1 CYP1B1 DDX43 ELOVL7 FADS2 ITPKC IVD PGK2 PISD SLC2A1 SLC2A2 STK40
CDK5R1 APOBEC1 COX6A2 CRHBP CUEDC1 HBB MAPT MXI1 PCSK1 RAG2 RPL13A SH3GL2 SIX6 SUMO4 TBX15 TRMT2B ZNF157
CBFA2T3 CCND2 EGF FGFR1 FOXP1 GATA4 HOXA10 HOXA11 MEF2D NES
TGFBR2 AFAP1L2 BCL9L CREBBP FOXS1 HEYL ID2 LHCGR NR2F2 RARG STAT5B
ATF4 PCNA PTPN1 NFE2L2 PLAU
Pearson Correlation
(Target Gene Prediction Ability)
0.005
0.030
Top Predicted Receiver
Targets Trending with SAR (δ) in Arteries
– Gene Differentially Expressed in Preeclampsia¹
Regulatory Potential
0.000 0.001 0.002 0.003 0.004 0.005 0.006 0.007 0.008

**Extended Data Figure S6 | EVT distribution. a.** Distribution of SAR continuous score $\delta$ per artery based on the presence of perivascular and/or intravascular EVTs. **b.** Scatter plot of $\log_2$(Intravascular/Perivascular) ratio by SAR continuous score $\delta$, for arteries with both perivascular and intravascular EVTs present. Black line, fitted linear regression model. **c.** Violin plot of the distribution of LD1 for EVTs, by anatomical location. Horizontal lines inside violins indicate mean LD1 value. **d.** Scatter plot of perivascular EVT1c ($CD56^+$) frequency by SAR continuous score $\delta$. Red line, fitted linear regression model. **e.** Percentage of arteries with scores less than or equal to a given SMA loss (s) threshold, by perivascular or intravascular EVTs present. Arteries were considered to have perivascular or intravascular EVT if the number of EVT in the appropriate artery compartment was >5. **f.** Percentage of arteries with scores less than or equal to a given endothelial loss (e) threshold, by perivascular or intravascular EVTs present. Arteries were considered to have perivascular or intravascular EVT if the number of EVT in the appropriate artery compartment was >5. **g.** Frequency of EVT populations by anatomical location. **h.** Violin plot of distance from artery (in pixels) of EVTs grouped by EVT type. **i-l:** see Methods "Origin of CD56+ EVTs in the intravascular compartment" **i.** Scatter plot of intravascular EVT1c ($CD56^+$) frequency by SAR continuous score $\delta$. Red line, fitted linear regression model. **j.** Bar plot of the EVT1c frequency increase rate, as a function of SAR continuous score $\delta$, by anatomical location. Rates presented are the linear regression slopes from **i**. Error bars, 95% confidence interval for regression slopes. **k.** Paired-by-artery plot of CD56 expression in EVT1a&b, comparing the perivascular and intravascular compartments. Arteries with $\delta \geq 2$ were included. p = 5e-03, Wilcoxon signed rank test. **l.** Proportion of $Ki67^+$ intravascular EVTs, by EVT type. **m.** Full heatmap for differentially expressed genes between intravascular and interstitial EVTs by Nanostring showing gene expression (Z-score), (logFC > 2, adj p-value <0.05) **n.** IHC validation on serial human decidua sections for protein counterparts for 3 genes found to be differentially expressed by Nanostring (JAG1, C5ORF30, and EBI3) along with controls HLA-G, CD56, and H&E. **o.** Full output for outcome of NicheNet's ligand activity prediction on differentially expressed genes on intravascular EVTs: results are shown for the 10 EVT-ligands best predicting receivers expressed in arteries, ranked by Pearson correlation coefficient or the EVT ligand activity ranking metric. Ligands, receivers, and targets also differentially expressed in Preeclamptic decidua samples[1] indicated in red.
